# Embryonic cortical layer 5 pyramidal neurons form an active, transient circuit motif perturbed by autism-associated mutations

**DOI:** 10.1101/2022.08.31.506080

**Authors:** Martin Munz, Arjun Bharioke, Georg Kosche, Verónica Moreno-Juan, Alexandra Brignall, Alexandra Graff-Meyer, Talia Ulmer, Tiago M. Rodrigues, Stephanie Haeuselmann, Dinko Pavlinic, Nicole Ledergeber, Brigitte Gross-Scherf, Balázs Rózsa, Jacek Krol, Simone Picelli, Cameron S. Cowan, Botond Roska

## Abstract

Cortical circuits are composed predominantly of pyramidal-to-pyramidal neuron connections, yet their assembly during embryonic development is not well understood. We show that embryonic layer 5 pyramidal neurons, identified through single cell transcriptomics, display two phases of circuit assembly in vivo. At E14.5, a multi-layered circuit motif, composed of a single layer 5 cell type, forms. This motif is transient, switching to a second circuit motif, involving all three types, by E17.5. In vivo targeted single cell recordings and two-photon calcium imaging of embryonic layer 5 neurons reveal that, in both phases, neurons have active somas and neurites, tetrodotoxin-sensitive voltage-gated conductances, and functional glutamatergic synapses. Embryonic layer 5 neurons strongly express autism-associated genes, and perturbing these genes disrupts the switch between the two motifs. Hence, layer 5 pyramidal neurons form transient active pyramidal-to-pyramidal circuits, at the inception of neocortex, and studying these circuits could yield insights into the etiology of autism.

## Introduction

The neocortex contains neuronal circuits that are highly interconnected, with the most common class of neurons, pyramidal neurons (Harris and Shepherd, 2015), receiving a majority of their connections from other pyramidal neurons (Brown et al., 2021; Kim et al., 2015; Vélez-Fort et al., 2014; Wertz et al., 2015; Young et al., 2021). Pyramidal neurons are of different types with distinct transcriptomic identities and are organized into distinct layers (Harris and Shepherd, 2015; Tasic et al., 2016, 2018). While the activity and communication within pyramidal-to-pyramidal neuron circuits have been intensively investigated, in vivo, in the adult (Cossell et al., 2015; Han et al., 2018; Keller et al., 2020; Kondo et al., 2016; Lee et al., 2016; Morishima and Kawaguchi, 2006; Petreanu et al., 2009; Wertz et al., 2015), when and how the first active pyramidal neuron circuits assemble in vivo are not known. Answers to these questions are central to understanding cortical circuit development. Moreover, since neurodevelopmental disorders are associated with defects within cortical circuits (Del Pino et al., 2018; Foss-Feig et al., 2017; Golden et al., 2018; Mullins et al., 2016; Robertson and Baron-Cohen, 2017), insights into pyramidal circuit formation may also be relevant for understanding the mechanisms of diseases such as autism spectrum disorder.

During cortical development, pyramidal neurons migrate to their final locations in neocortex, with layers forming in an inside-out fashion (Angevine and Sidman, 1961; Berry and Rogers, 1965; Greig et al., 2013; Jabaudon, 2017; Molyneaux et al., 2007; Rakic, 1972). Deep layer (layers 5 and 6) pyramidal neurons are born first, appearing in mice between embryonic day (E) 11.5 and E14.5, and starting to migrate into the developing cortex from E12.5 onwards (Cadwell et al., 2019; Greig et al., 2013; Kriegstein and Noctor, 2004; Molyneaux et al., 2007). Among deep layer neurons, layer 6 pyramidal neurons receive the majority of their inputs from upper layer neurons (Vélez-Fort et al., 2014), which arrive in cortex later, from E15.5 onwards. Therefore, most of the layer 6 pyramidal neuron circuitry likely forms only after E15.5. In contrast, layer 5 pyramidal neurons – which are composed of three cell types: near-projecting neurons (NP), intratelencephalic neurons (IT), and pyramidal tract neurons (PT) – receive 50% to 70% of their inputs, recurrently, from other layer 5 pyramidal neurons (Kim et al., 2015; Young et al., 2021). Therefore, the majority of layer 5 pyramidal neuron connectivity could form early in embryonic cortical development.

Neuronal activity starts during embryonic development (Allene and Cossart, 2010; Allène et al., 2008; Antón-Bolaños et al., 2019; Bortone and Polleux, 2009; Corlew et al., 2004; Galli and Maffei, 1988; Huang et al., 2020; Luhmann et al., 2016; Maffei and Galli-Resta, 1990; Mayer et al., 2019; McCabe et al., 2006; Ohtaka-Maruyama et al., 2018; Owens and Kriegstein, 1998; Owens et al., 1996; Yuryev et al., 2016, 2018). Cajal-Retzius cells, as early as E14.5 (Yuryev et al., 2016, 2018), and pyramidal neurons, from E15.5 onwards, show calcium transients in their somas, in vivo (Antón-Bolaños et al., 2019; Huang et al., 2020). Moreover, at E14.5, thalamic neurons display correlated activity in vitro, and this correlated activity can reach cortex by E16.5, when thalamic axons arrive in the cortex (Guillamón-Vivancos et al., 2022; Moreno-Juan et al., 2017). However, the time when cortical pyramidal neurons first assemble into circuits with other cortical pyramidal neurons, in vivo, and the point at which activity appears and becomes correlated within these circuits, remains unknown.

Neurons within circuits transmit activity using active conductances via neurites across synapses. Therefore, to understand if early born pyramidal neurons form active circuits, it is necessary to measure activity within their individual neurites, to find active conductances, and to detect functional synapses, in vivo. While in vivo two-photon imaging in embryos has been performed (Ang et al., 2003; Higuchi et al., 2016; Huang et al., 2020; Kawasoe et al., 2020; Yanagida et al., 2012; Yuryev et al., 2016, 2018), these recordings did not resolve activity within individual neurites and were not targeted to cell types. Further, thus far, no patch clamp recordings, capable of revealing active conductances, have been performed in embryonic neurons, in vivo.

Insight into the timing and assembly of the first active pyramidal neuron circuits is also important in the context of neurodevelopmental disorders such as autism spectrum disorder and schizophrenia (Marín, 2016). Indeed, common circuit dysfunctions have been hypothesized to underlie the phenotypic similarities across the heterogeneous mixture of genetic abnormalities associated with these neurodevelopmental disorders (Courchesne and Pierce, 2005; Del Pino et al., 2018; Foss-Feig et al., 2017; Golden et al., 2018; Kaiser and Feng, 2015; Mullins et al., 2016; Robertson and Baron-Cohen, 2017). Further, patches of disorganized cortical tissue have been observed in brains of both children with autism as well as in mouse models of the disorder, around the time of birth (Choi et al., 2016; Dufour et al., 2022; Kim et al., 2017; Orosco et al., 2014; Peñagarikano et al., 2011; Rabelo et al., 2022; Stoner et al., 2014; Yim et al., 2017). However, it remains unknown if mutations in autism-related genes may perturb the development of pyramidal-to-pyramidal neuron circuits during embryonic development.

Here, we developed a method with sufficient mechanical stability to perform both two-photon imaging from individual neurites and two-photon guided patch clamp recordings, in healthy living embryos connected to the dam. For imaging, we targeted embryonic layer 5 pyramidal neurons, identified through single cell RNA sequencing, since these neurons likely form the earliest pyramidal-to-pyramidal circuits. Embryonic layer 5 pyramidal neurons displayed calcium transients in vivo starting at E13.5, with two distinct phases of increased activity, separated by a transition phase. The first phase of increased activity showed a peak of activity at E14.5, while the second phase extended from E17.5 onwards. In the first phase of increased activity, embryonic layer 5 pyramidal neurons formed two layers composed of a single embryonic type, transcriptionally closest to the adult NP cell type. During the transition phase, the upper layer disappeared through programmed cell death, and neurons from the lower layer migrated to form a third layer. This third layer formed the adult layer 5, and is composed of three types of embryonic neurons, corresponding to the adult NP, IT, and PT layer 5 pyramidal neuron types. Already in the first phase of increased activity, at E14.5, layer 5 pyramidal neurons of the embryonic NP type showed signatures of active connected circuits: First, these neurons formed synapses, including synapses between pairs of embryonic layer 5 pyramidal neurons, constituting recurrent connectivity. Second, neurons displayed tetrodotoxin-(TTX) sensitive active conductances, and applying TTX decreased the activity of the neurons. Third, applying α-amino-3-hydroxy-5-methyl-4-isoxalone propionic acid (AMPA) and N-methyl-D-aspartate (NMDA) resulted in increased calcium fluorescence in embryonic layer 5 pyramidal neurons. Fourth, neurons displayed excitatory postsynaptic potentials. Fifth, the activity of neurons within and across the two layers was highly correlated. In addition, disrupting activity, from E14.5 onwards, shifted the spatial location of the embryonic neurons, within the developing cortex. Hence, embryonic cortical pyramidal neurons form two distinct multi-layered, active circuit motifs during embryonic development that switch after E15.5. The first constitutes a new circuit motif at the inception of the neocortex. Layer 5 pyramidal neurons also showed increased expression of genes associated with autism spectrum disorder during embryonic development compared to the adult. Remarkably, perturbing two different autism-related genes, selectively in embryonic layer 5 pyramidal neurons, resulted in a similar disruption of the circuit architecture: a disruption of the transition between the two layer 5 circuit motifs, with the first circuit motif persisting past E15.5, as well as the formation of patches of displaced embryonic layer 5 pyramidal neurons prior to birth.

## Results

### A population of genetically labelled embryonic cortical neurons has layer 5 pyramidal neuron identity

A mouse line expressing Cre in adult layer 5 pyramidal neurons, Rbp4-Cre KL100 (Gerfen et al., 2013; Gong et al., 2007), when crossed with a fluorescent Cre-reporter line, encoding either tdTomato (Madisen et al., 2010) or GCaMP6s (Madisen et al., 2015), marked cells with neuronal morphology close to the surface of the neocortex during embryonic development (Fig. 1A, B). Rbp4-Cre neurons first appeared at E13.5, a time when layer 5 pyramidal neurons first enter the preplate, the precursor to the cortical plate (Greig et al., 2013) (Fig. 1A). From E13.5 onwards, Rbp4-Cre neurons expressed Bcl11b, the expression of which is restricted in the adult cortex to layer 5 (Arlotta et al., 2005), suggesting that Rbp4-Cre neurons have layer 5 pyramidal neuronal identity already early in development (Fig. 1B, C). Since neuronal identity is more comprehensively defined by transcriptomic identity, assessed through single cell sequencing (Bandler et al., 2021; Di Bella et al., 2021; Loo et al., 2019; Tasic et al., 2016, 2018), we performed single cell RNA sequencing of cells from the developing cortex every day from E14.5 to E18.5, and identified Rbp4-Cre neurons, via tdTomato expression (Fig. S1). To compare the identity of embryonic Rbp4-Cre neurons with adult cortical neurons, we used a panel of 26 genes with layer-specific expression, selected from a published atlas of adult cortical cell type transcriptomes (Tasic et al., 2016, 2018). On all studied embryonic days, Rbp4-Cre neurons expressed marker genes associated with adult cortical layer 5 pyramidal neurons (Fig. 1D). Moreover, using those genes that best distinguished neurons in individual adult cortical layers (Tasic et al., 2016, 2018), the gene expression profile of embryonic Rbp4-Cre neurons was most highly correlated to that of adult layer 5 pyramidal neurons, for up to 120 genes (Fig. 1E). These experiments suggest that, during embryonic development, Rbp4-Cre neurons are embryonic layer 5 pyramidal neurons.

**Fig. 1.**
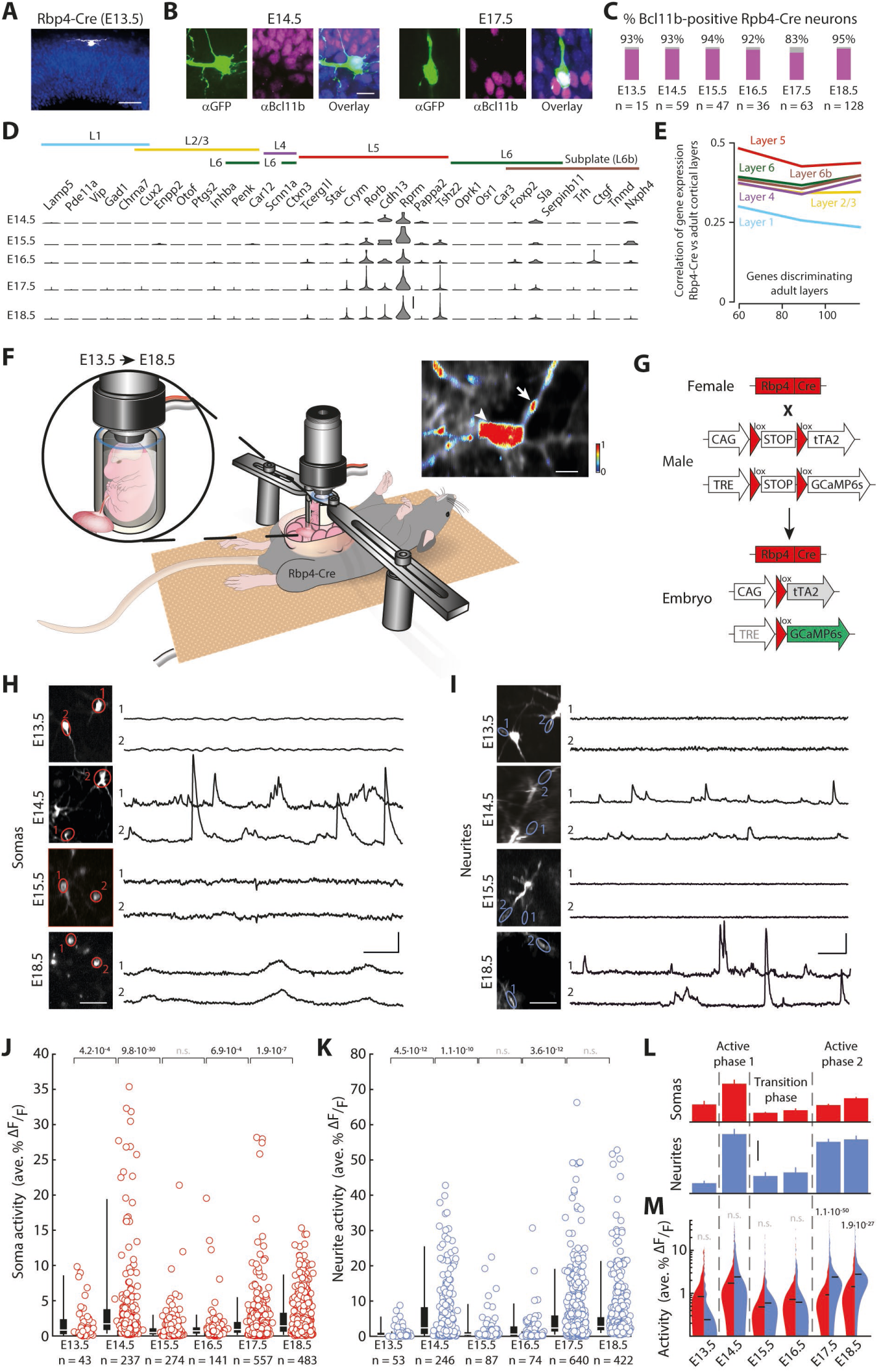
Rbp4-Cre neurons show two phases of increased spontaneous activity in both somas and neurites. (**A**) Rbp4-cre positive neuron at E13.5, near surface of developing cortex, immunostained against tdTomato (white), counterstained with Hoechst (blue). (**B**) Rbp4-Cre neurons (stained using GFP antibody labeling GCaMP6s, green) co-labeled with Bcl11b antibody (a marker of layer 5 neurons) at E14.5 (left) and E17.5 (right), counterstained with Hoechst (blue). (**C**) Fraction of Rbp4-Cre neurons expressing Bcl11b from E13.5 to E18.5. n = number of Rbp4-Cre neurons on each embryonic day. (**D**) Single cell RNA sequencing shows expression profile of layer-specific genes in Rbp4-Cre neurons from E14.5 to E18.5. Genes, selected as in (Schuman et al., 2019; Tasic et al., 2016, 2018), were ordered according to their specificity within adult cortical layers (L1 – L6b). (**E**) Correlation of gene expression profiles of embryonic Rbp4-Cre neurons compared to individual adult layers for up to 120 layer-specific genes, separately selected for each adult layer (Tasic et al., 2016, 2018) (**F**) Schematic diagram of embryonic para-uterine stabilization during in vivo two-photon calcium imaging in individual neurons’ somas and neurites, from E13.5 to E18.5. Embryo is immobilized within a holder matched to the size of the embryo, while maintaining its umbilical connection to the dam. The holder is partially immersed in the abdomen and stabilized by external supports. Top right: single embryonic neuron; arrowhead: soma activity; arrow: neurite activity; color: normalized calcium activity. (**G**) Mating strategy to drive GCaMP6s expression in embryonic cortical neurons. Female Rbp4-Cre mice (Gerfen et al., 2013; Gong et al., 2007) were crossed with male double transgenic mice, with two floxed STOP genes (tTA2 under the control of a CAG promoter (Miyamichi et al., 2011), and GCaMP6s under the control of a tTA2 responsive TRE promoter (Madisen et al., 2015)), resulting in embryos expressing GCaMP6s in Rbp4-Cre neurons. (**H** and **I**) Two-photon imaging of somas (red, **H**) and neurites (blue, **I**) of embryonic Rbp4-Cre neurons with two regions of interest (ROIs) (left) and their recorded activity traces (right) shown for each selected embryonic day (filtered to denoise and adjust for baseline shifts in the traces). (**J** and **K**) Activity of individual somas (**J**) and neurites (**K**) of Rbp4-Cre neurons from E13.5 to E18.5. Circles: activity of each neuron; box-and-whiskers: distributions across all cells at each embryonic day shown as box (25-75 percentile) and whisker (5-95 percentile); white line: median; n = number of neurons. (**L**) Mean activity across all somas (top, red) and neurites (bottom, blue) recorded on each embryonic day. Dotted line: separation of active phases and transition phase. (**M**) Comparison of distributions of calcium activity across somas (left, red) and neurites (right, blue) of Rbp4-Cre neurons from E13.5 to E18.5. (**J, K,** and **M**) Probability: Wilcoxon rank-sum test. Recordings from 3 (E13.5), 9 (E14.5), 5 (E15.5), 4 (E16.5), 5 (E17.5), and 6 (E18.5) embryos (activity in individual embryos labeled in Fig. S9). Scale bars: 50 μm (A), 10 μm (B), 20 transcripts (D), 10 μm (inset, F), 40 μm (left, H), 25s and 25 %ΔF/F (right, H), 40 μm (left, I), 25s and 50 %ΔF/F (right, I), 2 average %ΔF/F (M).

### Para-uterine imaging enables recordings of activity from neurites of pyramidal neurons in living embryos

To understand if embryonic layer 5 pyramidal neurons form active circuits in vivo, we developed a method that provides enhanced mechanical stability to the embryo, and thereby the resolution necessary to image individual neurites in vivo. To increase mechanical stability, we placed the embryo within a holder matched to the embryo’s size, filled the holder with agar, and placed a cover glass on top (Fig. 1F). Additionally, to isolate the embryo from the anesthetized dam’s movements due to breathing and beating of the dam’s heart, the holder was externally supported. In this way, embryos, ranging in weight from 0.17 g at E13.5 to 1.3 g at E18.5 (Fig. S2), were stable enough such that the amount of movement between imaging frames during two-photon imaging of the embryonic cortex was similar to that observed in adult mice during two photon imaging (Fig. S3). Moreover, to keep the embryo healthy during imaging, we held the embryo in the holder within the dam’s abdominal cavity adjacent to the uterus (“para-uterine”) (Fig. 1F), decreasing strain on the umbilical cord. Blood flow within the embryonic cortex was constant for at least 5 hours, as opposed to deteriorating within 10 minutes upon disruption of the umbilical cord (Videos S1, S2, Fig. S4). Further, the para-uterine location maintained the embryo at physiological temperature (Fig. S5) (Refinetti, 2010; Reitman, 2018). These improvements allowed the imaging of Rbp4-Cre neurons for at least 5 hours, with a resolution sufficient to resolve individual neurites.

We tested several systems for driving expression of calcium sensors in Rbp4-Cre neurons: adeno-associated viruses, herpes simplex viruses, electroporation, and mouse lines. Amongst these systems, GCaMP6s-tTA2 reporter mice, when crossed with Rbp4-Cre mice, generated GCaMP6s expression in Rbp4-Cre neurons throughout cortex, with sufficient signal-to-noise ratio for two-photon imaging during embryonic cortical development (Fig. 1F, G, S6). GCaMP6s-tTA2 mice were double transgenics that we created by crossing mice expressing tetracycline-controlled transactivator protein (tTA2) (Miyamichi et al., 2011) with mice expressing tTA2-dependent GCaMP6s (Chen et al., 2013; Madisen et al., 2015), both under the control of Cre (Fig. 1G). Rbp4-Cre neurons showed an average fluorescence that was 109% above the background, from E13.5 to E18.5 (Fig. S6). Therefore, the combination of “para-uterine imaging” and the use of GCaMP6s-tTA2 mice enabled the in vivo imaging of activity in both somas and neurites of Rbp4-Cre neurons, across the embryonic development of the cortical plate (Fig. 1H, I).

### Embryonic layer 5 pyramidal neurons show two phases of increased activity

At E14.5 and from E17.5 onwards, we observed spontaneous calcium events in both Rbp4-Cre somas and neurites (Fig. 1H, I, Videos S3-6) in the posterior part of cortex (posterior dorsal pallium; Fig. S7). Calcium event properties, such as the length and size, differed across embryonic days (Fig. S8). Therefore, to assess the overall change of activity, we quantified the total change of GCaMP6s fluorescence (%ΔF/F) across each recording, normalized per second (Fig. 1J, K, S9). In both Rbp4-Cre somas and neurites, spontaneous calcium activity significantly increased from E13.5 to E14.5. This was followed by a significant decrease in activity from E14.5 to E15.5 (Video S7). Activity stayed at a low level from E15.5 to E16.5. From E16.5 to E17.5, activity in both Rbp4-Cre somas and neurites significantly increased. In somas, the activity then increased further from E17.5 to E18.5 while, in neurites, the increased activity plateaued from E17.5 to E18.5. Although the dam was anesthetized during imaging, different anesthetics did not change the overall amplitude of spontaneous calcium activity in either active phase (Fig. S10), despite their distinct modes of action (Hemmings et al., 2019; Pavel et al., 2020). Therefore, we found two periods of time with increased spontaneous calcium activity, independent of the dam’s anesthesia: the first at E14.5, and the second starting at E17.5, separated by a transition phase from E15.5 to E16.5, where Rbp4-Cre neurons showed reduced activity (Fig. 1L). These two phases were also reflected in the fraction of active neurons (Fig. S8).

In the first phase of increased activity, somas and neurites showed no significant difference in activity from each other (Fig. 1M). In contrast, in the second phase of increased activity, neurites showed significantly higher activity than that in the somas. Therefore, the two phases of increased activity show a qualitative difference between the relative activity of somas and neurites of embryonic layer 5 pyramidal neurons.

### Embryonic layer 5 pyramidal neurons have active conductances

To determine if neurons that are active in the two phases have active conductances, we performed two-photon targeted patch clamp recordings, in vivo, from Rbp4-Cre neurons labeled with tdTomato in the current clamp configuration (Fig. 2A). Interestingly, all recorded Rbp4-Cre neurons, at E14.5 and E18.5, displayed active conductances as indicated by a nonlinear increase in peak voltage, with increasing current steps (Fig. 2B, C). These conductances disappeared upon application of a blocker of voltage-gated sodium channels, TTX (Fig. S11). In contrast, none of the recorded unlabeled cells in the vicinity of labeled Rbp4-Cre neurons displayed active conductances at E14.5 (Fig. S12). Even at E18.5, only half of nearby unlabeled cells had active conductances. Therefore, during both phases of increased calcium activity, Rbp4-Cre neurons display TTX-sensitive active conductances, suggesting that voltage-gated sodium channels may contribute to the increase in calcium activity.

**Fig. 2.**
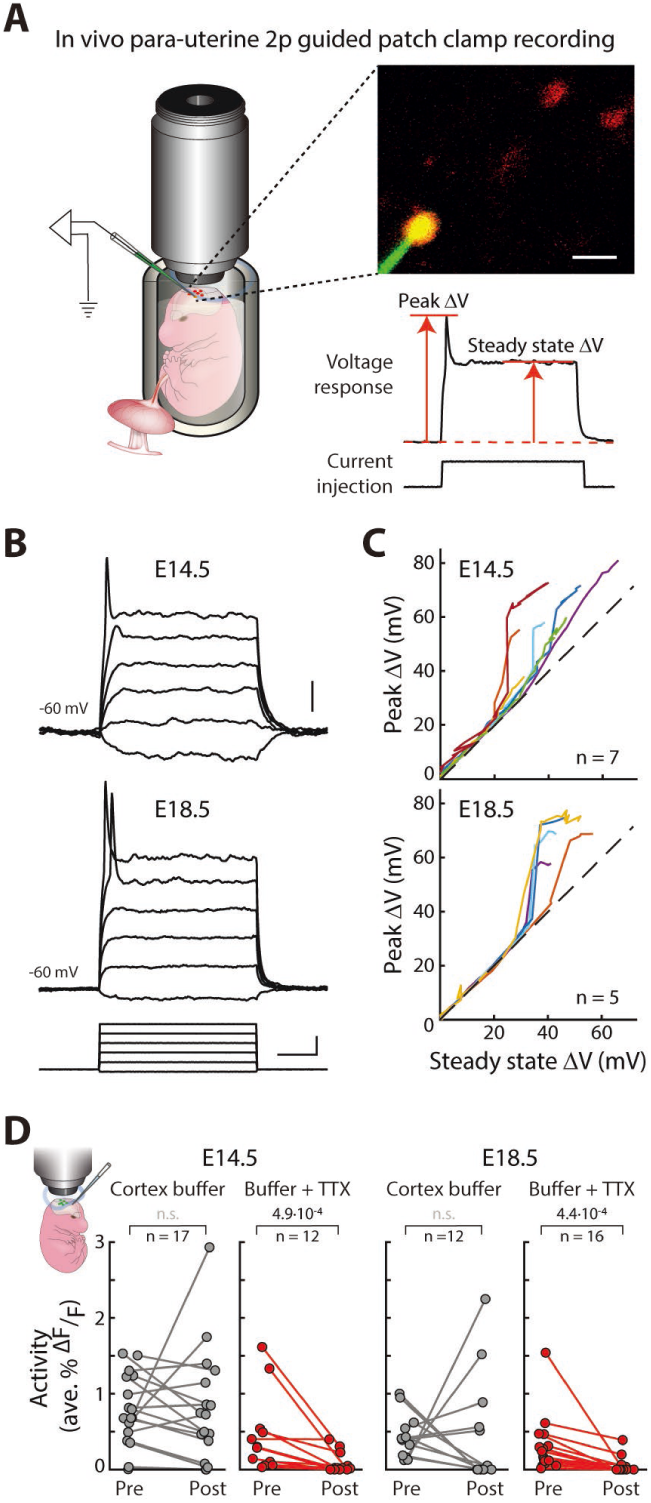
Rbp4-Cre neurons display active conductances during both active phases of embryonic development. (**A**) Left: schematic of para-uterine imaging combined with patch clamp recording to perform in vivo two-photon targeted electrophysiology from Rbp4-cre neurons. Right top: imaging field during patch clamp recording shows Rbp4-cre neurons (red), Alexa Fluor 488 filled pipette (green) and patched neuron (yellow) that is both Rbp4-cre positive and Alexa Fluor 488 filled. Right bottom: voltage trace showing peak and steady state voltage measurements, in response to current injection. (**B**) Voltage responses of an Rbp4-cre neuron (at E14.5 (top) and E18.5 (middle)) to graded intracellular current injections (bottom), recorded in the current clamp configuration. (**C**) Peak versus steady state voltage, recorded for different intracellular current injections, at E14.5 (top) and E18.5 (bottom), to visualize nonlinearities in Rbp4-Cre neurons (for current-voltage curves and comparisons with unlabeled cells in the vicinity of Rbp4-Cre neurons, see Fig. S12). n = number of neurons recorded from 5 (E14.5) and 4 (E18.5) embryos. (**D**) Calcium activity of embryonic Rbp4-Cre neurons at E14.5 and E18.5 before and after application of cortex buffer (control) or cortex buffer with TTX to the surface of cortex, obtained via in vivo para-uterine two-photon imaging. Probability: Wilcoxon signed rank test. n = number of Rbp4-Cre neurons recorded from 3 (E14.5) and 3 (E18.5) embryos. Scale bars: 20 μm (inset, A), 10 mV (top, B), 50 ms and 40 pA (bottom, B).

To test this, we imaged the change in spontaneous calcium activity in Rbp4-Cre neurons, before and after the application of TTX. At both E14.5 and E18.5, calcium activity significantly decreased following the application of TTX (Fig. 2D). In contrast, following the application of cortex buffer, as a control, Rbp4-Cre neurons showed no significant change in the spontaneous activity on either embryonic day. This suggests that active conductances, mediated by voltage-gated sodium channels, contribute to the recorded calcium activity during embryonic development.

### Embryonic layer 5 pyramidal neurons form transient layers

Given the two phases of increased spontaneous calcium activity across Rbp4-Cre neurons, we asked if the spatial organization of these neurons may also change across the same phases during embryonic development. To reveal the location of GCaMP6s-expressing Rbp4-Cre neurons, we collected brain sections every day from E13.5 to E18.5 and stained them with a GFP antibody (Fig. 3A). Rbp4-Cre neurons were distributed throughout the depth of the developing preplate at E13.5. By E14.5, Rbp4-Cre neurons had organized into two distinct layers: an upper layer near the surface of cortex, and a lower layer below the cortical plate. The number of Rbp4-Cre neurons within the upper layer decreased with time, with the maximum decrease occurring from E14.5 to E15.5, and all upper layer neurons completely disappearing before E17.5 (Fig. S13). This decrease was not due to GCaMP6s expression, since we found a similar decrease in Rbp4-Cre mice crossed with tdTomato reporter mice (Fig. S14), with a greater fraction of upper layer neurons staining positive for cleaved Caspase-3, a marker of apoptosis, than lower layer neurons (Fig. S13). From E15.5 onwards, a middle layer appeared, between the upper and lower layers. By E18.5, neurons in the middle layer adopted a morphology akin to adult layer 5 pyramidal neurons, with neurites extending to the surface of cortex (Fig. 3A). Therefore, Rbp4-Cre neurons exist in two different spatial configurations during the embryonic development of cortex: a lower layer and a transient upper layer around E14.5, and the same lower layer and a middle layer from E17.5 to E18.5. The transition between these two spatial configurations occurs from E15.5 to E16.5.

**Fig. 3.**
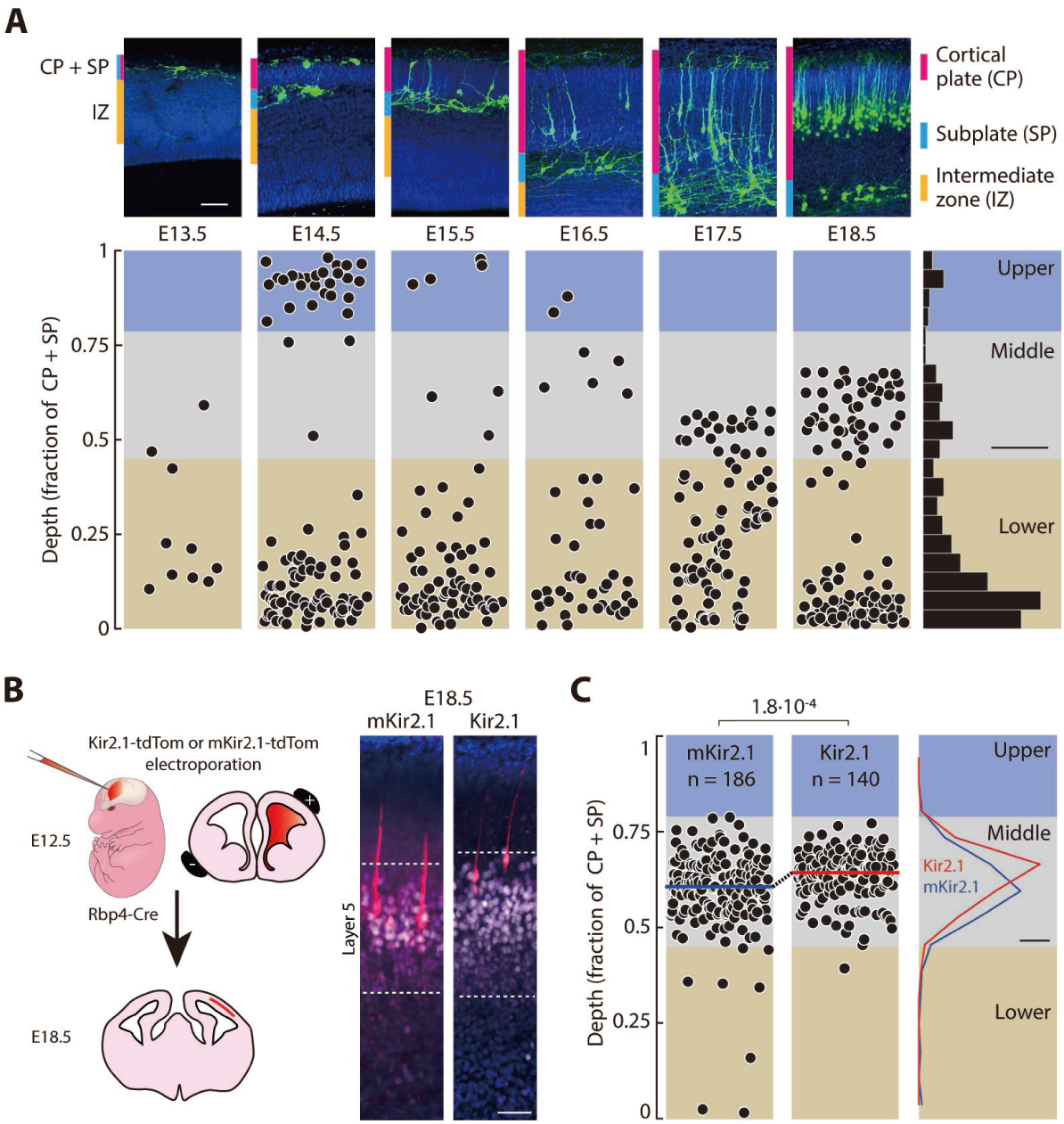
Rbp4-Cre neurons show changes in spatial organization aligned with the changes in activity during embryonic development. (**A**) Top: Rbp4-Cre neurons (stained using GFP antibody labeling GCaMP6s, green) from E13.5 to E18.5, within the cortical plate (magenta), subplate (cyan), and intermediate zone (yellow), counterstained with Hoechst (blue). Bottom left: distribution of Rbp4-Cre neuronal locations as a fraction of the cortical plate and subplate thickness, from E13.5 to E18.5. Bottom right: distribution of all Rbp4-Cre neuronal locations across days was used to identify distinct clusters by anatomical depth (blue: upper layer; gray: middle layer; beige: lower layer). (**B**) Left: schematic description of electroporations. E12.5 embryos were electroporated with either a plasmid conditionally expressing Kir2.1-tdTomato or mKir2.1-tdTomato. Expression was observed at E18.5. Right: Example electroporated embryos at E18.5 stained using tdTomato antibody (red: mKir2.1-tdTomato (left); Kir2.1-tdTomato (right)), Bcl11b (white), counterstained with Hoechst (blue). (**C**) Distribution of mKir2.1- or Kir2.1-positive neurons’ locations (from 10 mKir2.1-tdTomato electroporated embryos and 10 Kir2.1-tdTomato electroporated embryos), as a fraction of the cortical plate and subplate thickness at E18.5. Right: Distribution of Rbp4-Cre neurons (blue: mKir2.1; red: Kir2.1). Blue: upper layer; gray: middle layer; beige: lower layer. Boundaries set as from wildtype distribution from (A). n = number of electroporated Rbp4-Cre neurons. Scale bars: 50 μm (top, A), 10% (bottom right, A), 20 µm (B), 10% (C).

From E16.5 onwards, the number of neurons in the middle layer increased as the number of neurons in the lower layer decreased (Fig. 3A, S15). To test if this change in the proportion of neurons in the two layers is due to neurons in the lower layer migrating to the middle layer, we performed in vivo time-lapse imaging of Rbp4-Cre neurons across 5 hours, at E16.5. We observed neurons moving from the lower layer towards the middle layer, with an average speed of 3 µm/hour, a similar speed to what has been reported before (Boitard et al., 2015; Noda et al., 2019; Simó et al., 2010). Each hour, 4% of lower layer neurons moved into the middle layer. This suggests that neurons in the lower layer migrate to form the middle layer.

During development, the cortical plate is bordered above by the marginal zone, containing Cajal-Retzius cells (Kilb and Frotscher, 2016; Marín-Padilla, 1998; Sekine et al., 2014), and below by the subplate, containing subplate neurons (Allendoerfer and Shatz, 1994; Ohtaka-Maruyama, 2020). Throughout embryonic development, upper layer Rbp4-Cre neurons appear close to the surface of cortex, while lower layer neurons are localized within the subplate, raising the possibility that some Rbp4-Cre neurons may have Cajal-Retzius or subplate identity. However, we found that genes previously identified as markers of Cajal-Retzius cells (Bandler et al., 2021; Di Bella et al., 2021; Kuang et al., 2010) identified a population of neurons with distinct gene expression profiles from Rbp4-Cre neurons (Fig. S16). Similarly, genes identified as markers of subplate neurons (Di Bella et al., 2021; Tasic et al., 2016) also overlapped in a population of neurons distinct from Rbp4-Cre neurons, and Rbp4-Cre neurons did not express the common subplate marker, Nr4a2 (Fig. S17). Instead, at E14.5, in both upper and lower layers, Rbp4-Cre neurons express Bcl11b (Fig. S18). Similarly, at E18.5, Rbp4-Cre neurons in both the middle and lower layers also express Bcl11b. Therefore, Rbp4-Cre neurons have layer 5 neuronal identity, independent of their physical depth within cortex, and do not show Cajal-Retzius nor subplate identity, despite their physical location close to, or within, the marginal zone and subplate.

### Perturbation of activity alters the layered organization of embryonic layer 5 pyramidal neurons

Since both the activity and spatial organization of Rbp4-Cre neurons changed in a coordinated way across embryonic days, we asked if perturbing the activity of Rbp4-Cre neurons may change their spatial localization. At E12.5, we performed in utero electroporations of embryos with a plasmid expressing, in a Cre-dependent manner, either the inward rectifier potassium channel Kir2.1 (Arcangeli et al., 1995) or an engineered non-conducting Kir2.1 channel (mKir2.1) as a control (Tinker et al., 1996; Xue et al., 2014), both fused to a tdTomato fluorescent protein (Fig. 3B). The electroporated plasmids enter progenitor neurons at E12.5 and, once Cre becomes active in postmitotic Rbp4-Cre neurons, should result in the hyperpolarization of the Cre-expressing postmitotic neurons (Angevine and Sidman, 1961; Berry and Rogers, 1965; Greig et al., 2013; Jabaudon, 2017; Molyneaux et al., 2007; Rakic, 1972). To reveal the location of Kir2.1-tdTomato- and mKir2.1-tdTomato-expressing Rbp4-Cre neurons, we collected brain sections at E18.5 and stained them with a tdTomato antibody (Fig. 3B). Compared to mKir2.1-tdTomato expressing Rbp4-Cre neurons, Kir2.1-tdTomato expressing Rbp4-Cre neurons localized significantly closer to the surface of cortex (Fig. 3C). Therefore, hyperpolarization of postmitotic embryonic layer 5 pyramidal neurons affects their location within the neocortex at E18.5.

### Embryonic layer 5 pyramidal neurons divide into three cell types

Adult layer 5 pyramidal neurons divide into three cell types: intratelencephalic neurons (IT), pyramidal tract neurons (PT), and near-projecting neurons (NP) (Groh et al., 2010; Kim et al., 2015). To understand if embryonic layer 5 pyramidal neurons also divide into distinct cell types, we performed clustering of the transcriptomes of individual Rbp4-Cre neurons (Becht et al., 2019; Traag et al., 2019). We found three distinct clusters of transcriptomic identity (Fig. 4A). In each embryonic cluster, most of the neurons had gene expression profiles corresponding to one of the three adult types, with conditional probability >99% (Fig. 4B, S19). This implies a one-to-one correspondence between the adult types and embryonic clusters, and hence, we named cluster 1 as embryonic NP, cluster 2 as embryonic IT, and cluster 3 as embryonic PT. The three embryonic types showed distinct relative proportions on each embryonic day (Fig. 4C). At E14.5, all embryonic layer 5 pyramidal neurons were of the embryonic NP type. From E15.5 onwards, the fraction of embryonic IT and PT neurons gradually increased. These results suggest that embryonic layer 5 pyramidal neurons divide into three embryonic cell types, and this cell type composition is developmentally regulated.

**Fig. 4.**
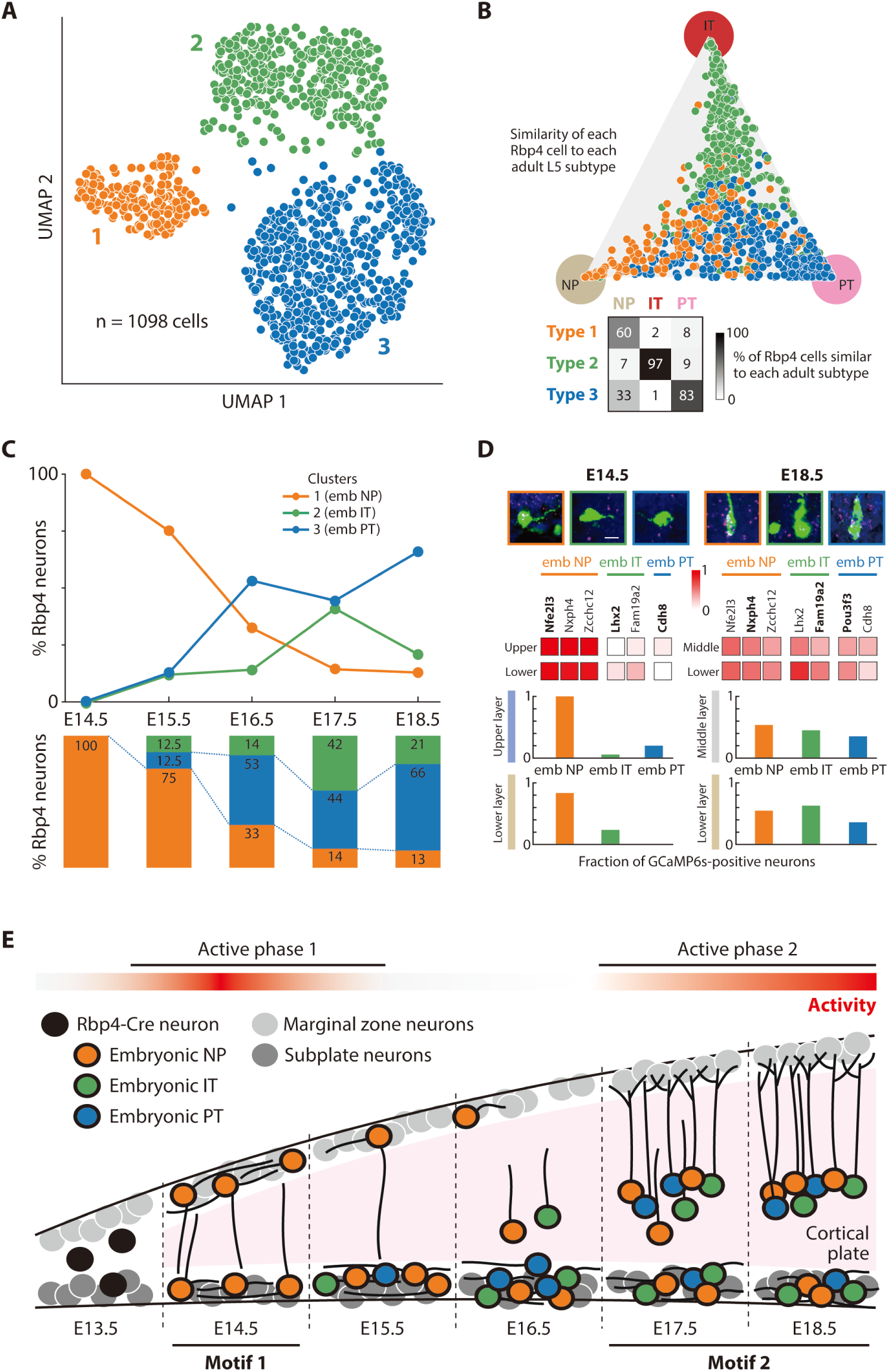
Rbp4-Cre neurons show changes in cell type composition aligned with the changes in activity and spatial organization during embryonic development. (**A**) UMAP embedding of 1098 Rbp4-Cre neurons’ transcriptomes (Becht et al., 2019). Color: Leiden clusters (Traag et al., 2019). (**B**) Top: Embryonic Rbp4-Cre types (colored as in (**A**)) embedded in a triangular representation representing the conditional probability of each cell’s expression profile being sampled from the three adult layer 5 types: near-projecting neurons (NP) (beige), intratelencephalic neurons (IT) (red), pyramidal tract neurons (PT) (pink) (Tasic et al., 2018), computed through a Bayesian approach. Adult layer 5 types are defined through the gene expression profiles of 24 differentially expressed genes (see Fig. S19 for the result of varying the number of differentially expressed genes). Bottom: Percent of Rbp4-Cre neurons of each embryonic type associated with each adult type with probability greater than 0.99. (**C**) Percent of neurons from each embryonic type on each embryonic day (colored as in (**A**)). Types defined by their closest adult type in (**B**); type 1: embryonic NP; type 2: embryonic IT; type 3: embryonic PT. (**D**) Top: Example in situ hybridizations of type-specific markers (magenta) in Rbp4-Cre neurons (green), counterstained with Hoechst (blue); box color: embryonic types colored as in (**A**). Middle: Fraction of GCaMP6s-positive Rbp4-Cre neurons containing embryonic NP, embryonic IT, and embryonic PT-specific in situ hybridization markers, in each layer, at E14.5 (left) and E18.5 (right); bold: genes shown in example in situ hybridizations, above. Bottom: combining all type-specific markers reveals fractions of embryonic NP, embryonic IT, and embryonic PT Rbp4-Cre neurons in each layer at E14.5 (left) and E18.5 (right). (**E**) Schematic of development of layer 5 pyramidal neurons from embryonic day 13.5 to 18.5 depicting alignment of phases of activity, changes in cell type composition, and spatial organization. Scale bars: 10 μm (top, D).

So far, our results show that embryonic layer 5 pyramidal neurons divide into three transcriptomic cell types and are organized into three distinct layers. To reveal how these three cell types divide into the three layers, we performed in situ hybridization against genes that are preferentially expressed in the embryonic NP, embryonic IT, and embryonic PT types (Fig. 4D, S20). At E14.5, both upper and lower layers were composed predominantly of embryonic NP neurons. This confirms the previous finding that, at E14.5, all Rbp4-Cre neurons formed a single cluster corresponding to the embryonic NP type. At E18.5, both middle and lower layers show all three embryonic types. Therefore, upper layer neurons are always embryonic NP type. At E14.5, lower layer neurons are also embryonic NP type, but the cell type composition gradually changes to include all three cell types by E18.5. Similarly, the middle layer is a mixture of all three types (Fig. 4D, E).

Taken together, during embryonic development, layer 5 pyramidal neurons show a switch in activity, spatial organization, and cell type identity, that are coordinated in time (Fig. 4E). Initially, embryonic layer 5 pyramidal neurons are present in a highly active lower-and-upper layer configuration composed of a single layer 5 pyramidal neuron type, embryonic NP neurons. This constitutes the first organizational motif, which persists until E15.5. From E15.5 to E16.5, activity significantly decreases, coinciding with a decrease in upper layer neurons, and a migration to form a middle layer. By E17.5, embryonic layer 5 pyramidal neurons switch to a new, lower-and-middle layer configuration, which persists until birth. This second motif again shows increased activity, but contains all three embryonic layer 5 types.

### Embryonic layer 5 pyramidal neurons form synapses

To identify whether embryonic layer 5 pyramidal neurons could form cortical circuits with other embryonic layer 5 pyramidal neurons, we explored the expression of the molecular machinery underlying neuronal communication in Rbp4-Cre neurons of all three embryonic types. Many genes associated with synaptic function and active neuronal membrane properties were expressed within all three embryonic cell types, across developmental days E14.5 to E18.5 (Fig. 5A, S21). This included genes involved in chemical synaptic transmission, such as AMPA receptor subunits and NMDA receptor subunits, genes involved in electrical synaptic transmission, such as Connexin 45 (*Gjc1*), genes involved in pre- and postsynaptic signaling, such as *Erc1* and PSD-95 (*Dlg4*), genes associated with the synaptic vesicle cycle, such as *Snap25* and *Vamp2*, as well as genes necessary for active neuronal membrane properties, such as voltage-gated calcium, sodium, and potassium channels (Fig. 5A, S21) (Kanehisa and Goto, 2000; Siddiqui and Craig, 2011). Therefore, the molecular components underlying neuronal communication are present within all three embryonic types of layer 5 pyramidal neurons across embryonic days, from E14.5 to E18.5.

**Fig. 5.**
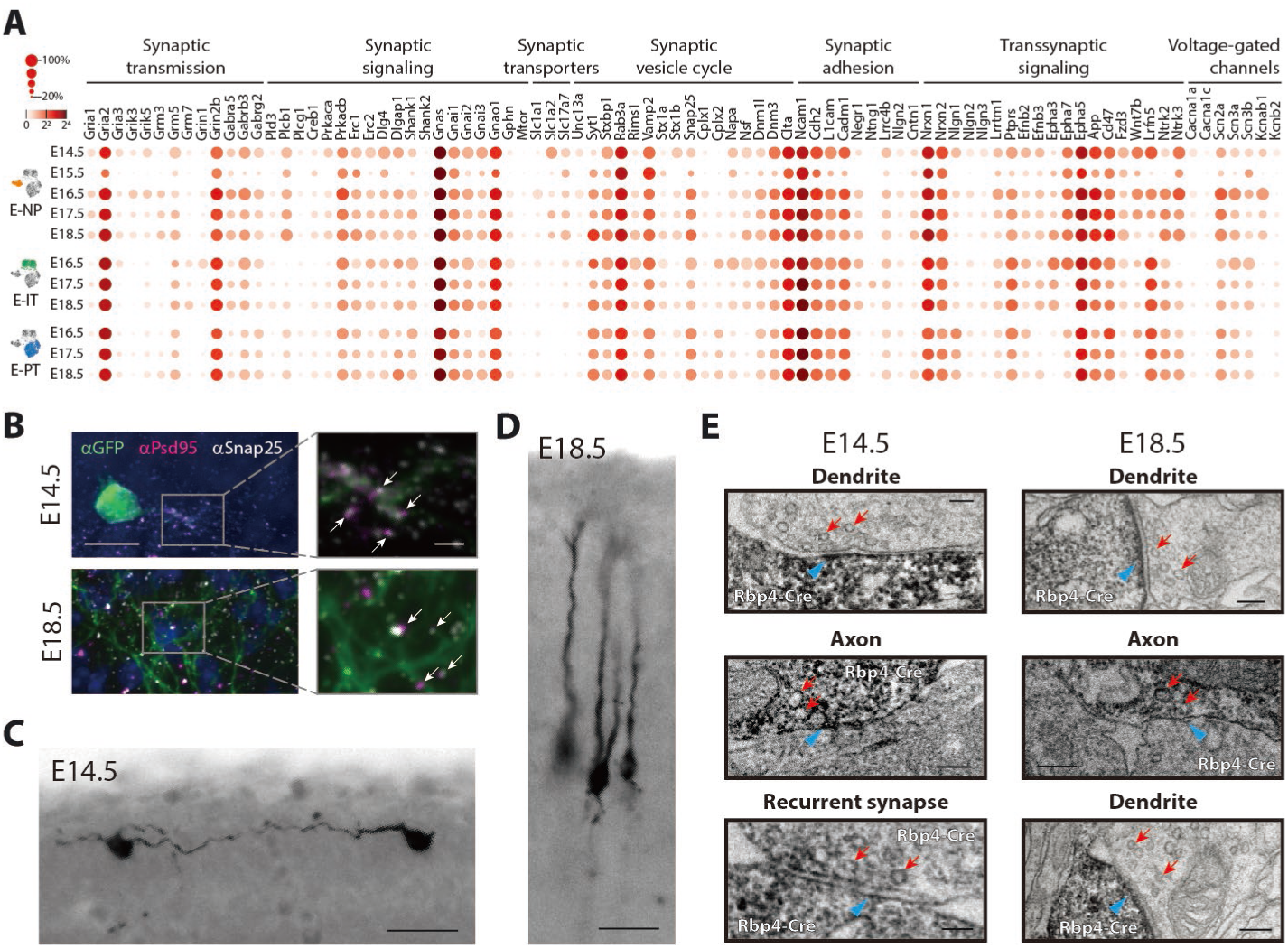
Rbp4-Cre neurons form synapses starting at E14.5. (**A**) Expression of selected genes (subset from Fig. S21) related to synaptic formation, synaptic function, and neuronal active membrane properties in all three embryonic types at all ages (Kanehisa and Goto, 2000; Siddiqui and Craig, 2011). Transcript counts shown as colored circles. Radius of circles: fraction of cells expressing the gene; color of circles: mean normalized transcripts per cell (log_2_). (**B**) Snap25 (presynaptic; magenta) and PSD-95 (postsynaptic; gray) show punctate expression within Rbp4-Cre neurons (green) at E14.5 (top) and E18.5 (bottom). Enlarged images (right) show colocalization of Snap25 and PSD-95 puncta (arrows). (**C** and **D**) Immunohistochemical staining of Rbp4-cre neurons (stained using GFP antibody in combination with secondary antibody coupled to horseradish peroxidase to catalyze the formation of DAB reaction product) at E14.5 (**C**) and at E18.5 (**D**) in tissue prepared for electron microscopy (EM) (Knott et al., 2009). (**E**) EM images of synaptic contacts involving Rbp4-cre neurons (darker cells, marked by electron dense DAB product) at E14.5 (left) and E18.5 (right), with presynaptic vesicles (red arrow) and postsynaptic densities (blue arrowhead). Scale bars: 10 μm (left, B), 2 μm (right, B), 25 μm (C), 25 μm (D), 100 nm (E).

We then explored if embryonic layer 5 pyramidal neurons form synapses, given prior evidence of pyramidal-to-pyramidal neuron synapses, early in postnatal development (Anastasiades and Butt, 2012; Yu et al., 2009). First, we performed immunostaining against both a presynaptic marker, Snap25, and a postsynaptic marker, PSD-95. Both markers showed punctate labeling on the neurites of Rbp4-Cre neurons, at E14.5 and at E18.5 (Fig. 5B). Additionally, Snap25 and PSD-95 colocalized in many puncta. Next, we performed electron microscopy to identify the presence of synaptic specializations onto or from Rbp4-Cre neurons (Fig. 5C–E). Using Rbp4-Cre mice crossed with GCaMP6s-tTA2 mice, Rbp4-Cre neurons were stained with a primary anti-GFP antibody, and visualized under electron microscopy through a 3,3′-diaminobenzidine tetrahydrochloride hydrate (DAB) reaction product catalyzed by horseradish peroxidase coupled to a secondary antibody (Knott et al., 2009). Both at E14.5 and at E18.5, we observed close junctional contacts involving Rbp4-Cre neurons, with synaptic vesicles marking the presynaptic side and electron dense structures (postsynaptic densities) marking the postsynaptic side (Fig. 5E). Some of these synapses were marked with DAB on the presynaptic side, while others had DAB-labeling on the postsynaptic side. Additionally, at E14.5, we observed synapses with DAB-labeling on both the presynaptic and postsynaptic sides i.e. recurrent synapses between two Rbp4-Cre neurons. This suggests that embryonic layer 5 pyramidal neurons do both form and receive synaptic contacts and, already at E14.5, pairs of embryonic layer 5 pyramidal neurons have recurrent synapses.

### Embryonic layer 5 pyramidal neurons are sensitive to AMPA and NMDA and show excitatory synaptic potentials

To determine if the synapses during embryonic development are functional, we applied agonists of glutamatergic synaptic transmission (AMPA and NMDA) while imaging from Rbp4-Cre neurons (Fig. 6A). On both E14.5 and E18.5, applying a mixture of AMPA and NMDA onto cortex was closely followed by a significant increase in calcium fluorescence in all imaged Rbp4-Cre neurons (Fig. 6B). In contrast, following an application of cortex buffer alone, Rbp4-Cre neurons showed no significant change in calcium fluorescence. Additionally, following application of TTX, in vivo two-photon targeted patch clamp recordings from Rbp4-Cre neurons in current clamp configuration displayed spontaneous excitatory synaptic potentials at both E14.5 and E18.5 (Fig. S22). The response to glutamatergic agonists and the presence of excitatory synaptic potentials suggest that embryonic layer 5 pyramidal neurons form functional synapses during embryonic development.

**Fig. 6.**
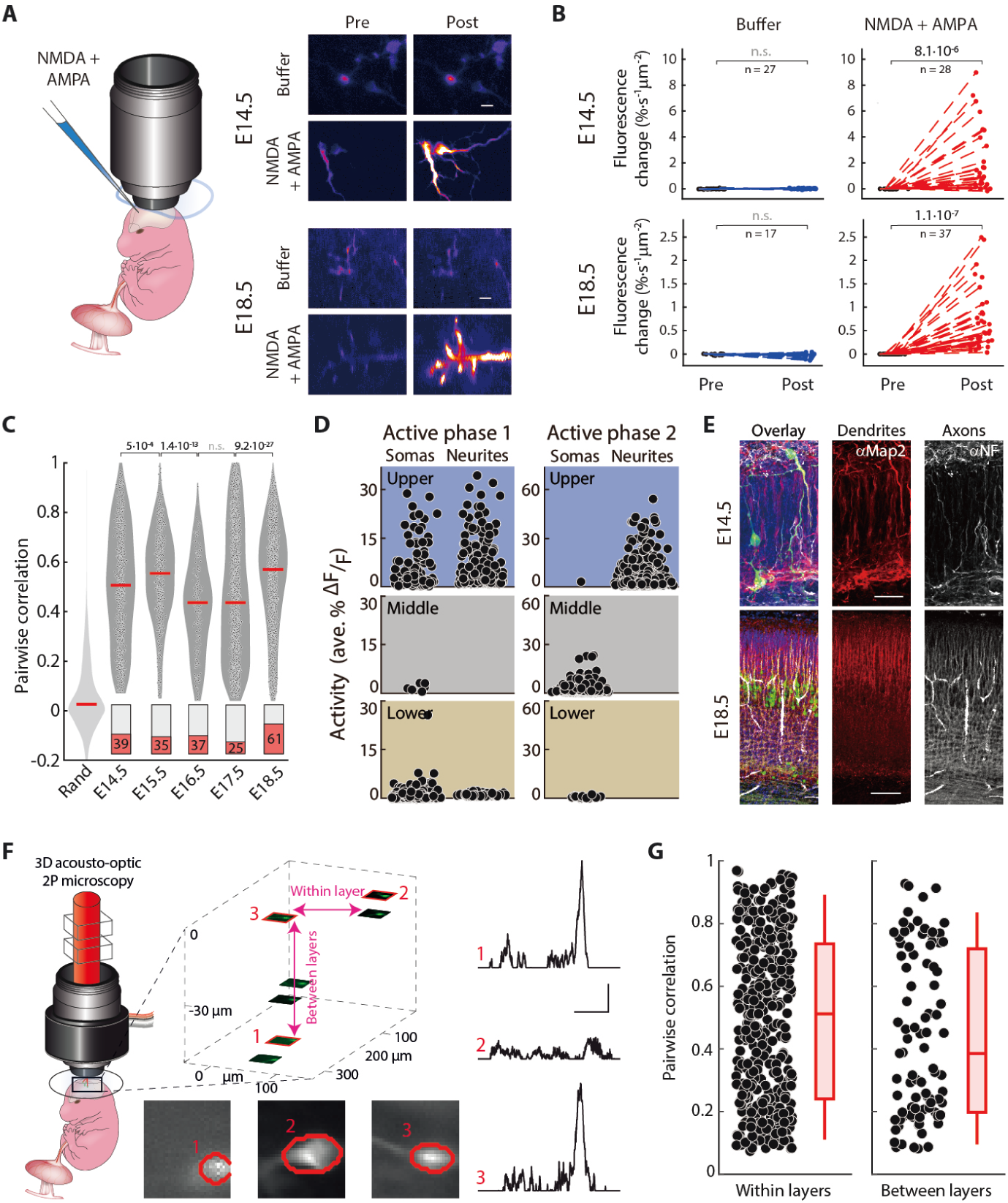
Rbp4-Cre neurons form active circuits starting at E14.5. (**A**) Left: schematic of injection of a mixture of cortex buffer with NMDA and AMPA (NMDA + AMPA) during in vivo embryonic two-photon para-uterine imaging. Right: Rbp4-Cre neurons before and after injection of either cortex buffer (control) or NMDA+AMPA at E14.5 and E18.5. (**B**) Change in fluorescence from baseline both before (Pre) and after (Post) application of either cortex buffer (blue) or NMDA + AMPA (red). Probability: Wilcoxon signed rank test; n = number of Rbp4-Cre ROIs from 3 (E14.5) and 3 (E18.5) embryos. (**C**) Correlations of spontaneous calcium activity, that are significantly greater than random, between pairs of Rbp4-Cre neurons on each embryonic day from E14.5 to E18.5. Filled circles: pairwise correlations; dark gray shading: distribution; red line: median. Random: pairwise correlations modelling the distribution of correlations from shuffling events in each Rbp4-Cre neurons; light gray shading: distribution. Bars: percent of neuron pairs with correlations greater than random (red) on each embryonic day. Probability: Wilcoxon rank-sum test. n = 425 (E14.5), 575 (E15.5), 187 (E16.5), 2517 (E17.5) and 2171 (E18.5) pairs of embryonic neurons recorded from 3 (E13.5), 9 (E14.5), 5 (E15.5), 4 (E16.5), 5 (E17.5), and 6 (E18.5) embryos. (**D**) Activity across somas and neurites, in the two active phases, within each spatial layer (blue: upper layer; gray: middle layer; beige: lower layer; as defined in Fig. 3). (**E**) Immunohistochemical staining of Rbp4-Cre neurons (stained using GFP antibody, green), dendrites (stained using Map2 antibody, red), and axons (stained using NF antibody, white) at E14.5 and E18.5, within the developing cortex, counterstained with Hoechst (blue). (**F**) Left: schematic representation of in vivo para-uterine imaging of embryonic cortex using a 3D acousto-optic two photon microscope, allowing for random-access to cells within both layers, simultaneously, at E14.5. Top middle: mean projections of three-dimensional volumes around each neuronal soma in the three-dimensional imaging field are shown, with three examples (red outline) detailed (bottom middle). Examples are selected from both layers. Right: Δf/f activity traces from example neuronal somas (as labeled on left). Cells 1 and 3 are an example of a pair between layers with high correlation. (**G**) Correlations of spontaneous calcium activity, that are significantly greater than random, recorded as shown schematically in (**F**) between pairs of neurons within the same layer (left) and pairs of neurons in different layers (right). Dots: pairs of neurons; red box-and-whiskers: distribution as box (25-75 percentile) and whisker (5-95 percentile); red line: median. Scale bars: 10 μm (A), 30 μm (top, E), 100 μm (bottom, E), 20s and 5 %ΔF/F (F).

### Activity in embryonic layer 5 pyramidal neurons is correlated

Communication between neurons can lead to correlated activity. From E14.5 to E18.5, we observed many pairs of Rbp4-Cre neurons with correlations significantly greater than expected from neurons with random activity (Fig. 6C). This correlated activity did not extend across the whole population, either through synchronous activity (Fig. S23) (Bharioke et al., 2022) or through propagating waves of activity (Fig. S24), but was apparent in pairs of Rbp4-Cre neurons. At E14.5, during the first phase of increased activity, both upper and lower layers had active neurons (Fig. 6D). Further, embryonic layer 5 pyramidal neurons extended axons and dendrites both within and between the layers (Fig. 6E). Hence, we asked if neurons communicate only within their own layer, or across layers. Using a 3D acousto-optic two-photon microscope, we recorded simultaneously from neurons in both layers, throughout the depth of embryonic cortex at E14.5 (Fig. 6F). Pairs of neurons were not only significantly correlated within the same layer but also across layers, in both cases showing correlation coefficients as high as 0.9 (Fig. 6G). Taken together, the significant pairwise correlations suggest that, from E14.5 onwards, embryonic layer 5 pyramidal neurons communicate with each other.

### Cell type specific perturbation of autism-associated genes results in persistence of upper layer and patchy disorganization of upper and middle layer neurons

Because neurodevelopmental disorders have been associated with cortical circuit dysfunction (Birnbaum and Weinberger, 2017; Del Pino et al., 2018; Ebert and Greenberg, 2013; Forrest et al., 2018; Forsyth et al., 2020; Gilman et al., 2011; Jabaudon, 2017; Mullins et al., 2016; Stoner et al., 2014), we analyzed the expression of neurodevelopmental disease-associated genes in embryonic layer 5 pyramidal neurons and compared this with expression in adult layer 5 pyramidal neurons (Tasic et al., 2018). A number of genes associated with schizophrenia (Lee et al., 2019; Ripke et al., 2014) and autism spectrum disorder (Abrahams et al., 2013) were expressed across all three embryonic types of Rbp4-Cre cells (Fig. 7A, S25). Embryonic gene expression associated with both neurodevelopmental diseases was significantly higher than the average expression across all genes, with schizophrenia-associated genes showing about three-fold higher expression and autism-associated genes showing four-fold higher expression (Fig. 7B, S25). Importantly, while schizophrenia-associated genes were expressed similarly in both embryonic and adult layer 5 pyramidal neurons, autism-associated genes showed a four-fold increase in expression in embryonic layer 5 pyramidal neurons compared to the adult (Fig 7B, S25).

**Fig. 7.**
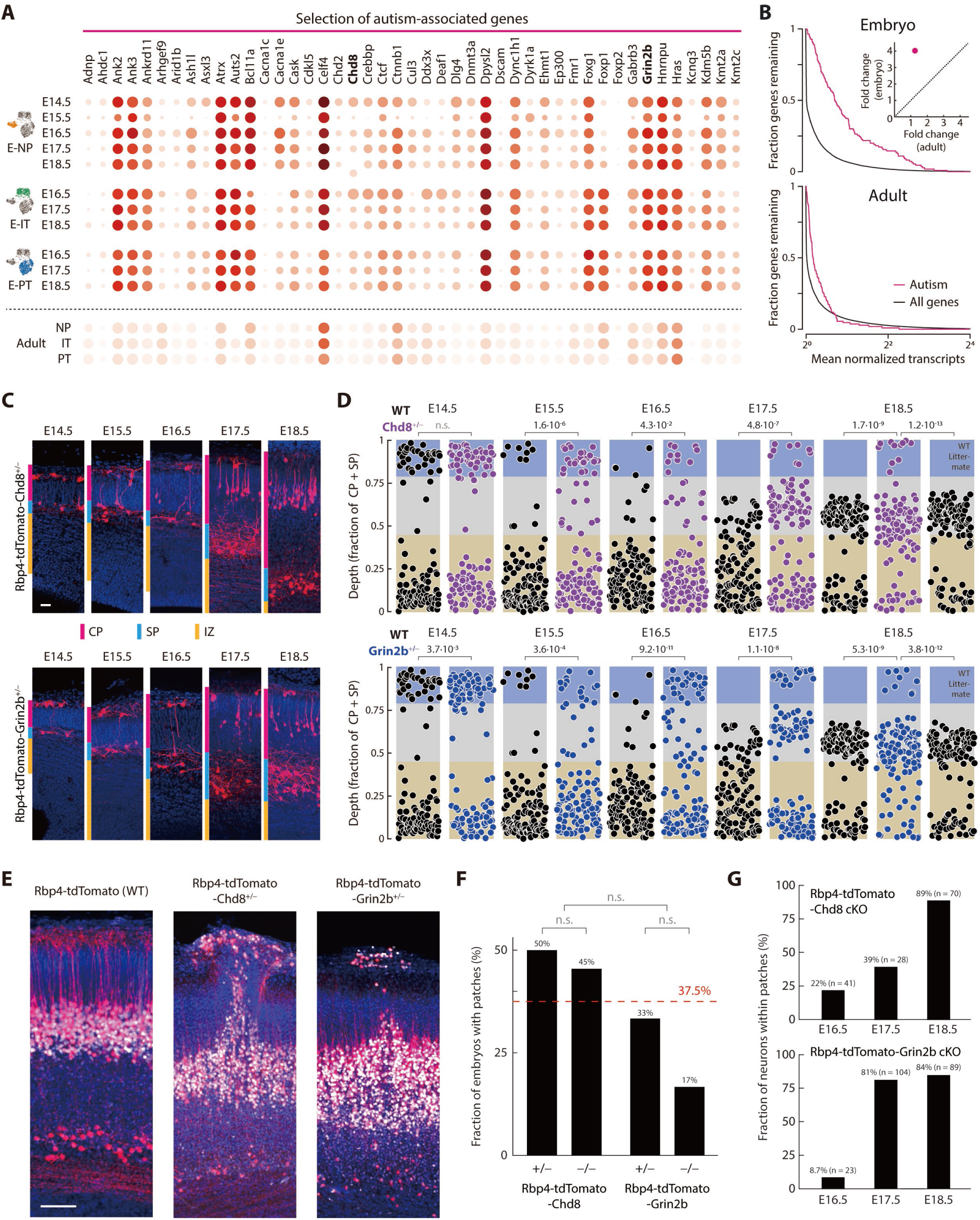
Perturbing autism-associated genes selectively in Rbp4-Cre neurons disrupts organization of layer 5 during embryonic development. (**A**) Expression of selected genes (subset from Fig. S25) related to synaptic formation, synaptic function, and neuronal active membrane properties in all three embryonic types at all ages (Kanehisa and Goto, 2000; Siddiqui and Craig, 2011). Transcript counts shown as colored circles. Radius of circles: fraction of cells expressing the gene; color of circles: mean normalized transcripts per cell (log_2_). (**B**) Fraction of genes with a mean transcript count greater than any specific mean transcript count are shown, for all genes (black), and genes associated with autism spectrum disorder (magenta) in embryonic Rbp4-Cre neurons (top) and adult layer 5 pyramidal neurons (bottom). Inset: Fold change of gene expression compared to all genes in embryos and adult. (**C**) Rbp4-Cre neurons (stained using tdTomato antibody, red) in Rbp4-tdTomato-Chd8^+/-^ (top) and Rbp4-tdTomato-Grin2b^+/-^ (blue) mice, from E14.5 to E18.5, within the cortical plate (magenta), subplate (cyan), and intermediate zone (yellow), counterstained with Hoechst (blue). See Fig. S26 for examples of Rbp4-Cre neurons for homozygous conditional knockout mice, for both genes. (**D**) Distribution of Rbp4-Cre neuronal locations as a fraction of the cortical plate and subplate thickness, from E14.5 to E18.5, in control (WT, black), Rbp4-tdTomato-Chd8^+/-^ (top, purple), and Rbp4-tdTomato-Grin2b^+/-^ (bottom, blue) mice. 125 neurons from each mouse line, sampled at random, displayed on each embryonic day. Layer boundaries derived from Fig. 3A (blue: upper layer; gray: middle layer; beige: lower layer). See Fig. S26 for distribution of Rbp4-Cre neuron locations for homozygous conditional knockout mice, for both genes. Probability: χ^2^ test comparing the fraction of Rbp4-Cre neurons in each layer between the conditional knockout mouse and control mice, on each embryonic day; p = 0.05. (**E**) Local patches of disorganization in Rbp4-tdTomato-Chd8^+/-^ and Rbp4-tdTomato-Grin2b^+/-^ mice, at E18.5, show Rbp4-Cre neurons (stained using tdTomato antibody, red) at the surface and disrupted middle layer, including both neurons expressing Cre (red) and not expressing Cre (stained using Bcl11b antibody, white), counterstained with Hoechst (blue). (**F**) Fraction of mutant mice, of each genotype, showing at least one patch, summed across embryonic days E16.5 to E18.5. Red line: Average across all four genotypes. Data from Rbp4-tdTomato-Chd8^+/-^ (14 embryos), Rbp4-tdTomato-Chd8^-/-^ (11 embryos), Rbp4-tdTomato-Grin2b^+/-^ (9 embryos), and Rbp4-tdTomato-Grin2b^-/-^ (6 embryos). Probability: Fisher exact test comparing fraction of embryos with patches in each genotype. p = 0.05, prior to Bonferroni correction for 3 tests. (**G**) Fraction of neurons within the upper layer that are located within patches of disorganization, in embryos with at least one patch. n = number of upper layer neurons on each embryonic day. Scale bars: 20 μm (C), 50 μm (E).

Therefore, we perturbed two known autism-associated genes, *Chd8* and *Grin2b* (Abrahams et al., 2013; De Rubeis et al., 2014; Iossifov et al., 2015), specifically in Rbp4-Cre neurons, and investigated their effect on the spatial organization of these neurons, during embryonic development. Since autism can be associated with both heterozygous and homozygous mutations (Busch et al., 2019; De Rubeis and Buxbaum, 2015; Frazier, 2019; Schaaf et al., 2011; Yin and Schaaf, 2017), we bred *Chd8* and *Grin2b* Cre-conditional knockout mouse lines with Rbp4-Cre and tdTomato Cre-reporter mice to produce embryos with Rbp4-Cre neurons both expressing tdTomato (Madisen et al., 2010) and being either heterozygous or homozygous knockout for the gene *Chd8* (Rbp4-tdTomato-Chd8^+/-^ and Rbp4-tdTomato-Chd8^-/-^ mice) or *Grin2b* (Rbp4-tdTomato-Grin2b^+/-^ and Rbp4-tdTomato-Grin2b^-/-^ mice). Control mice had tdTomato in Rbp4-Cre neurons but were wildtype for *Chd8* and *Grin2b*.

To reveal the spatial distribution of Rbp4-Cre neurons in the four different mutant mouse lines and controls, we collected brain sections from E14.5 to E18.5 and stained them with a tdTomato antibody (Fig. 7C, S26). Remarkably, in all four mutant mouse lines but not in the control line, we observed Rbp4-Cre neurons near the surface of the neocortex, persisting until E18.5. Indeed, the number of upper layer neurons observed from E15.5 onwards on each embryonic day was significantly higher in all four conditional mouse lines compared to control mice (Fig. 7D, S26). Therefore, a common effect of perturbing two different and unrelated autism-associated genes in layer 5 embryonic neurons was the abolishment of the transience of the upper layer. In control mice, the lower-and-upper layer configuration persisted only until E15.5. Following that day, the circuit switched to a lower-and-middle layer configuration. In contrast, in all four mutant mice, the upper layer persisted and therefore, following E15.5, embryonic layer 5 pyramidal neurons were found in all three layers.

In addition to the persistence of the upper layer after E15.5, in a subset of mutant animals, we also observed the presence of non-uniform patches of disorganized structure involving Rbp4-Cre neurons. These patches were characterized by a cluster of closely spaced Rbp4-Cre neurons, at the surface of the neocortex, together with a distortion of the middle layer, directly under the cluster of surface neurons, such that the middle layer extended closer to the neurons at the surface. We found disorganized patches in all four mutant mouse lines, on all embryonic days, from E16.5 onwards (Fig. 7E, S27). In control mice, we occasionally observed several Rbp4-Cre neurons at the surface of cortex (in 3 out of 31 embryos), but no associated disorganization of the underlying middle layer (Fig. S27). Averaging across all four lines, 37.5% of embryos showed patches of disorganization within cortex, from E16.5 onwards (Fig. 7F). Further, within the subset of mice that showed disorganized patches, at E18.5, greater than 80% of neurons at the surface of cortex were located within patches whereas, at E16.5, less than 20% of neurons were located within patches (Fig. 7G). Notably, this form of local patches of disorganization is reminiscent of patches of disorganization observed in the cortex of children with autism (Stoner et al., 2014).

## Discussion

When and how the first active pyramidal neuron circuits form is a central question in understanding the development of the neocortex (Katz and Shatz, 1996; Li et al., 2010; Verhage et al., 2000). Since layer 5 pyramidal neurons are born early in embryonic development and have the highest degree of recurrent connectivity in the adult, we argued that these can form one of the earliest pyramidal-to-pyramidal neuron circuits. Using mouse genetics and confirmed by single cell transcriptomics, we selected embryonic layer 5 pyramidal neurons and recorded from them using in vivo para-uterine two photon imaging as well as patch clamp recordings. We found that layer 5 pyramidal neurons formed pyramidal-to-pyramidal neuron circuits earlier than expected, and these circuits were organized into two distinct active circuit motifs, with a transition marked by a decrease in activity. Remarkably, selectively perturbing distinct autism-associated genes within embryonic layer 5 pyramidal neurons led to a common disruption of the transition between the two circuit motifs.

Prior work has shown that layers of pyramidal neurons are added sequentially, in an inside-out fashion, during the formation of the cortex (Angevine and Sidman, 1961). In addition to this sequential assembly, we found that at E14.5, at the inception of the neocortex, embryonic layer 5 pyramidal neurons formed two spatially distinct layers, both of which were composed of neurons of the embryonic NP type. This multi-layered organization of embryonic NP neurons at E14.5 is therefore a new motif in cortical development. Further, these neurons also participated in active circuits. The evidence for active circuits is the presence of chemical synapses, including recurrent synapses between layer 5 pyramidal neurons, the TTX-sensitive active conductances, the sensitivity of activity to TTX, the neurons’ sensitivity to AMPA and NMDA, the recorded excitatory postsynaptic potentials, and the correlated activity within and across the two layers. Therefore, the two-layered neuronal structure at E14.5 likely represents one of the first active neuronal circuits of cortex formed by pyramidal neurons.

Following this, at E15.5, the upper layer underwent programmed cell death and the activity in the remaining neurons decreased. Therefore, the two-layered neuronal structure formed at E14.5 is transient both in its structure and in its activity. From E16.5 on, the development of layer 5 pyramidal neurons was consistent with previous models of cortical development: the lower layer neurons migrated up to form a middle layer, containing precursors of all three adult layer 5 pyramidal neuron types, that further developed into the adult layer 5. Interestingly, from E17.5 onwards, we detected increased activity within dendrites, compared to the somas, potentially reflecting inputs driven, directly or indirectly, by the incoming axons of thalamic principal neurons (Antón-Bolaños et al., 2019; Guillamón-Vivancos et al., 2022; Moreno-Juan et al., 2017).

The spatial organization of the transient pyramidal-to-pyramidal neuron circuit at E14.5 is reminiscent of previously identified subplate to Cajal-Retzius circuits (Allendoerfer and Shatz, 1994; Kanold, 2019; Molnár et al., 2020; Riva et al., 2019). However, based on several lines of evidence: single cell transcriptomic identity of Rbp4-Cre neurons, in situ hybridization of marker mRNA and antibody staining of marker proteins, the neurons described in this study had neither subplate nor Cajal-Retzius cell identity (Fig. S16, S17). Rather, at E14.5, the Rbp4-Cre neurons were embryonic NP type layer 5 pyramidal neurons. Additionally, previous work has found that pyramidal neurons participate in transient synaptic circuits with subplate neurons during their migration to their final location in neocortex (Ohtaka-Maruyama et al., 2018), and that the organization of cortical circuits is instructed through subcortical inputs after E16.5 (Antón-Bolaños et al., 2019; Moreno-Juan et al., 2017). However, transient circuits between embryonic pyramidal neurons have not been previously described. We show such circuits, prior to the thalamic innervation of the cortical plate (López-Bendito and Molnár, 2003; Molnár et al., 1998; Moreno-Juan et al., 2017).

Hyperpolarization of cortical neuronal progenitors cells, and their subsequent daughter neurons, results in a change in the layer-specific identity of both upper and lower layer cortical neurons (Hurni et al., 2017; Vitali et al., 2018). Here, we restricted the hyperpolarization only to the postmitotic embryonic layer 5 pyramidal neurons. Hyperpolarized embryonic layer 5 pyramidal neurons are shifted towards the surface, however, they remain within layer 5, and continue to express a molecular marker of layer 5 identity. Therefore, manipulating activity or resting membrane potential during embryonic development acts both on cell identity, as shown previously (Vitali et al., 2018), as well as on fine circuit organization, as we have demonstrated here.

Neurodevelopmental disorders, such as autism spectrum disorder, have been associated with defects in cortical circuits (Birnbaum and Weinberger, 2017; Del Pino et al., 2018; Ebert and Greenberg, 2013; Forrest et al., 2018; Forsyth et al., 2020; Gilman et al., 2011; Jabaudon, 2017; Mullins et al., 2016; Stoner et al., 2014; Writing Committee for the Attention-Deficit/Hyperactivity Disorder et al., 2021). Notably, a recent genetic study of autistic patients suggested that, during development, autism-associated genes are strongly enriched in maturing excitatory neurons, compared to immature progenitors. Further, in human cortical organoids, mutations in autism-associated genes affect the maturation and circuit formation of deep layer cortical pyramidal neurons (Fu et al., 2022; Paulsen et al., 2022). Here, using conditional perturbations of two different autism-associated genes, we demonstrated a change in the transition between the two embryonic circuit motifs formed by layer 5 pyramidal neurons. Specifically, upper layer neurons persisted through embryonic development until E18.5, one day prior to birth. Further, an increasing fraction of these upper layer neurons clustered together, at the surface, as the embryos developed. These clusters were associated with a disorganization of the developing neocortex directly under each cluster, such that layer 5 neurons were displaced towards the surface. Intriguingly, this disorganization also involved neurons that did not express Cre, suggesting communication between the Cre-expressing subset of neurons and other layer 5 pyramidal neurons. Further, the patchy disorganization of mouse cortex that we report is reminiscent of the patchy disorganization of cortical tissue observed in children with autism (Rabelo et al., 2022; Stoner et al., 2014).

Taken together, our work identifying embryonic layer 5 pyramidal-to-pyramidal neuron active circuit motifs, together with the in vivo imaging and recording methods that we have developed, provide an opportunity to study the effects of genes associated with neurodevelopmental disorders, and in particular autism spectrum disorder, on identified circuits in the living embryo.

## Acknowledgments

We thank D. Jothimani, E. Mace, A. Herrero-Navarro, R. Morikawa, E. S. Ruthazer, N. Verma for comments on the manuscript, S. Gong for the sequence of the Cre used in the Rbp4-Cre KL100 mouse line, V. Juvin of SciArtWork for the visualization of the para-uterine imaging setup, and members of the Roska lab for discussions on the project. The project was supported by a European Molecular Biology Organization Long Term Fellowship (ALTF 1552-2015) co-funded by Marie Curie Actions, and a Human Frontier Science Program Postdoctoral Fellowship (LT000447/2016) to M.M., a Swiss National Science Foundation Synergia grant (CRSII3_141801), a European Research Council advanced grant (RETMUS N°669157, HURET N°883781), a Louis-Jeantet Foundation award, a Körber Foundation award, a Swiss National Science Foundation grant (31003A_182523), and the NCCR ‘Molecular Systems Engineering’ grant to B.R.

## Author contributions

M.M. designed para-uterine imaging, planned the experiments, managed the breeding of mouse lines, performed para-uterine surgery for calcium imaging, electrophysiology, and pharmacological manipulations, performed two-photon imaging, processed imaging data, performed and quantified immunohistochemistry, prepared tissue for electron microscopy, designed and performed single cell RNA sequencing experiments, planned single cell RNA sequencing analysis, planned in situ hybridizations, and wrote the paper. A. Bharioke designed para-uterine imaging, planned the experiments, performed two-photon imaging during calcium imaging and pharmacological manipulations, setup the imaging data pipeline, analyzed two-photon calcium recordings, designed single cell RNA sequencing experiments and analysis, planned in situ hybridizations, quantified immunohistochemistry, and wrote the paper. C.C. analyzed single cell RNA sequencing data. G.K. performed visually targeted electrophysiology recordings in embryos in vivo. V. M-J performed in utero electroporations and performed immunohistochemistry. A. Brignall processed imaging data and performed in situ hybridizations. A.G-M. performed immunolabeling of tissue for electron microscopy, fine sectioning, and electron microscopy imaging. T.U. performed and quantified immunohistochemistry. T.R. performed in situ hybridizations. S.H. performed FACS. D.P. performed single cell RNA sequencing. N.L. managed the breeding of mouse lines. B.G. managed the breeding of mouse lines. B. Rózsa developed two-photon microscopes. J.K. performed FACS. S.P. performed single cell RNA sequencing. B. Roska designed experiments and wrote the paper.

## Materials and Methods

### Animals

Animal experiments were performed in accordance with standard ethical guidelines (European Communities Guidelines on the Care and Use of Laboratory Animals, 86/609/EEC) and were approved by the Veterinary Department of the Canton of Basel-Stadt. The following transgenic mouse lines were used: Rbp4-Cre KL100 (Gerfen et al., 2013; Gong et al., 2007), Ai94(TITL-GCaMP6s) (Madisen et al., 2015) crossed with CAG-stop-tTA2 (Miyamichi et al., 2011) (GCaMP6s x tTA2), Chd8^em3Lutzy^, Grin2b^em5Lutzy^/J and Ai9(RCL-tdT) (Madisen et al., 2010). All mice were of C57BL/6 background, 55-120 days old, and maintained on a normal 12-hour light/dark cycle in a pathogen-free environment with ad libitum access to food and drinking water.

### Timed pregnancies

To determine embryonic age, we bred GCaMP6s^ki/ki^ x tTA2^ki/ki^ or Ai9^ki/ki^ males to Rbp4-Cre^tg/tg^ females on a strict schedule where males were placed in the females’ cage in the evening and plugs were checked the following morning. Pregnant females (dams) were monitored and pregnancy was confirmed visually 10 days after the detection of the plug. Experiments were then performed from E13.5 - E18.5.

For generating mice that are heterozygote and homozygote conditional knockouts for Chd8 or Grin2b as well as littermate controls, we bred, conditional Chd8^+/-^ x Rbp4^tg/-^ mice to conditional Chd8^+/-^ x Ai9^ki/ki^ mice and conditional Grin2b^+/-^ x Rbp4^tg/-^ mice to conditional Grin2b^+/-^ x Ai9^ki/ki^ mice. In addition, we also bred, conditional Chd8^+/-^ x Rbp4^tg/-^ mice to conditional Chd8^-/-^ x Ai9^ki/ki^ mice and conditional Grin2b^+/-^ x Rbp4^tg/-^ to conditional Grin2b^-/-^ x Ai9^ki/ki^ mice to increase the number of embryos homozygous for a conditional knockout of *Chd8* or *Grin2b*. To control for germline mutations (Steinmetz et al., 2017), we also used heterozygous embryos from crosses between conditional Chd8^-/-^ x Ai9^ki/ki^ mice or conditional Grin2b^-/-^ x Ai9^ki/ki^ mice to Rbp4^tg/tg^ mice.

### Para-uterine embryonic stabilization

Dams, pregnant with embryos between E13.5 and E18.5, were anesthetized with a mixture of Fentanyl-Medetomidine-Midazolam (Fentanyl (Janssen, 0.05 mg/kg); Medetomidine (Virbac, 0.5 mg/kg); Midazolam (Sintetica, 5 mg/kg)) through the initial surgery to prepare each embryo and the subsequent imaging session. A nasal mask was used to deliver humidified oxygen to the dam, at a flow rate of 0.8 l/min (Respironics EverFlo OPI pump, Phillips). For experiments where recordings were performed under Isoflurane anesthesia, dams were anesthetized via 1.75% Isoflurane (Provet) delivered using a Respironics EverFlo OPI pump, Phillips, UniVet Porta Anesthetic machine (using a Datex Ohmeda Isotec 5 continuous flow vaporizer), Groppler Medizintechnik). The desired mixture of isoflurane with humidified oxygen was generated by the vaporizer and applied to the mouse through a nasal mask (Groppler Medizintechnik), with a flow rate of 0.8 l/min.

Dams were placed on their back on a heating pad (TC1000, CWE), centered on a movable metal plate (MB1515/M, Thorlabs), and the hair on the abdomen was removed. A 1.5 cm long incision was placed along the midline of the abdomen. A heat reflective foil was placed above the abdomen to decrease heat loss. The abdominal skin and wall was retracted using the CD+ Labs System (retractor tip blunt (ACD-012); latex elastomer (ACD-011); magnetic fixator (magnet replaced by metal pin to fit plate) (ACD-002) (Southmedic Inc.)). A single uterine horn was isolated within the abdominal cavity, and the uterus surgically opened. A single embryo, within its yolk sac, was then isolated, and the yolk sac opened. The embryo was then placed in a custom 3D-printed holder. Low-melting point agar at 37 - 39 ℃ was applied around the embryo to fill the space between the embryo and the holder. A titanium headplate was glued to the embryo holder using UV glue (NOA 68, Norland Optical Adhesive), with the circular opening centered above the head of the embryo. The titanium headplate was tightly screwed to custom metal connectors, which were in turn attached to the metal plate through posts (TR30V/M, Thorlabs), thereby holding the embryo holder securely in position within the abdominal cavity of the dam, with the embryo’s body mostly immersed. HBSS (14175046, Gibco) warmed to 37 ℃ was used to fill the abdominal cavity of the dam. To improve optical access, the skin above the developing cortex was then removed, prior to imaging, creating an imaging region (Fig. S7). In addition, for embryos older than E14.5, the cranial bone was removed. A glass coverslip (10 mm round, 631-0665, VWR (Marienfeld)) was glued to the holder such that the glass covered the embryonic brain, applying minimal pressure on the brain. The metal plate supporting the dam and the immobilized embryo was then transferred to a two-photon microscope for imaging.

### In-vivo para-uterine two-photon imaging

Dams were anesthetized and maintained at a surgical depth, as assessed by the lack of spontaneous movement, as well as the absence of a paw withdrawal reflex, in response to a toe pinch. To maintain the anesthesia, top-up doses of FMM anesthesia (10 – 30% of the initial bolus) were injected into the dam, between recordings, approximately every 30 minutes. Throughout the experiment, a nasal mask was used to deliver humidified oxygen to the dam, at a flow rate of 0.8 l/min (Respironics EverFlo OPI pump, Phillips). For experiments where recordings were performed under Isoflurane anesthesia, dams were maintained under anesthesia via 1.75% Isoflurane (Provet) delivered using a Respironics EverFlo OPI pump, Phillips, UniVet Porta Anesthetic machine (using a Datex Ohmeda Isotec 5 continuous flow vaporizer), Groppler Medizintechnik). The desired mixture of isoflurane with humidified oxygen was generated by the vaporizer and applied to the mouse through a nasal mask (Groppler Medizintechnik), with a flow rate of 0.8 l/min.

Dams were monitored throughout all recording sessions using an IR camera within the recording setup (DMK 22BUC03, Imaging Source). The dam’s breathing remained rapid and shallow, without gasping, a hallmark of stable anesthesia at surgical depth (Adams and Pacharinsak, 2015). In addition, a pulse oximeter attached to a sensor applied to the rear paw (MouseSTAT Jr and Paw Sensor, Kent Scientific) was used to monitor the heart rate and blood oxygenation of each mouse during experiments. For all dams, the heart rate was maintained at less than 400 bpm (Fleischmann et al., 2016), and the blood oxygenation never dropped below 90%, when averaged over a minute.

GCaMP6s expressing neurons were imaged using three different two-photon laser scanning microscopes (Trachtenberg et al., 2002).

The first microscope was a Femtonics galvo-galvo scanning microscope equipped with a 25X water immersion objective (APO LWD 25X/1.10W, Nikon). Imaging was performed at 5 - 10 Hz within the developing embryonic cortex, at 920 nm. Each recording of spontaneous calcium activity, from E13.5 to E18.5, was 10 minutes in length.

The second microscope used was an acousto-optic (AO) microscope (Katona et al., 2012; Szalay et al., 2016) at 920 nm, also equipped with a 25X water immersion objective (APO LWD 25X/1.10W, Nikon). This microscope was used for simultaneous imaging across a three-dimensional (3D) cortical volume. Multi-cube scanning mode (Szalay et al., 2016) as well as a volume scanning mode (high-speed arbitrary frame scan) were used. For multi-cube scanning, 3D imaging regions composed of multiple imaging planes, centered around individual neurons or neurites, were selected manually within the imaged embryonic cortical volume. Each imaging plane was chosen to be approximately twice the size of the selected object in the horizontal dimensions. 5 – 7 imaging planes, generating a cube, were selected in the vertical dimension, to account for any remaining movement within the imaging volume. Recording speed varied with the number of recorded objects. Each recording was 5 minutes in length. For high-speed arbitrary frame scan mode, 9 – 11 planes were selected with a distance of 3 – 5 microns between each plane. All recordings were 10 minutes in length.

The third two-photon microscope used was a FemtoSMART resonant-galvo scanning microscope. It was equipped with an Olympus 16X water immersion objective (0.8 NA). This microscope was used for simultaneous imaging and electrophysiology.

The temperature of the embryo was supported at 37.5 °C using a temperature controller (TC1000, CWE), with the temperature probe held within the abdominal cavity of the dam. In addition, the objective was heated using an objective heater set at 50 °C (OWS-1, Warner Instruments) to further decrease the rate of heat loss from the embryo (Fig. S5).

For the in vivo imaging of Rbp4-Cre anatomy, 3D volumes within the cortex of Rbp4-Cre x Ai9 embryos at E16.5 were imaged for 5 hours, plane-by-plane, using the galvo-galvo two-photon microscope at 950 nm.

### Electrophysiology in combination with in vivo para-uterine imaging

For electrophysiological recordings we used whole-cell patch pipettes, pulled from borosilicate glass with filament (O.D.: 1.5 mm, I.D.: 0.86 mm) using a P-97 micropipette puller (Sutter Instrument Company) and filled with intracellular solution containing: 0.2 mM EGTA, 130 mM K-gluconate, 4 mM KCl, 2 mM NaCl, 10 mM HEPES, 4 mM ATP-Mg, 0.3 mM GTP-Tris, 14 mM phosphocreatine-Tris, 0.050 mM Alexa-488 and brought to pH 7.25 (with dilute NaOH or HCl) and 292 mOsm (by addition of H_2_O).

Embryos were prepared, as described for para-uterine imaging, above, with the exception of the coverslip which was cut to allow access of the patch pipette, and the opening of the dura. A ground wire was positioned in the immersion solution, near the surface of the brain (cortex buffer: 125 mM NaCl, 5 mM KCl, 10 mM glucose, 10 mM HEPES, 2 mM MgSO_4_ and 2 mM CaCl_2_). Signals (sampled at 50 kHz and low pass filtered at 10 kHz) were recorded using a National Instruments Board connected to a Multiclamp 700B amplifier (Molecular Devices).

The whole-cell patch pipette, filled with green fluorescent dye, was then lowered through the opening in the dura under positive pressure (100 mbar). Dye from the pipette filled the extracellular space, the pressure was lowered (25-35 mbar) and the pipette was advanced diagonally. Rbp4-Cre positive neurons were visualized by their expression of tdTomato. Adjacent non-Rbp4-Cre neurons were identified by the absence of red fluorescence, and by the injection of dye into the extracellular space.

The pressure on the pipette was further reduced just before touching the membrane of the target neuron, and finally released to form a gigaohm seal under visual guidance. Slow and fast pipette capacitances were compensated, and whole-cell access was achieved by applying a negative pulse of pressure on the pipette.

Current steps were applied to the patched cell, and the resulting voltage change was recorded, both in the presence and absence of TTX. Additionally, subthreshold membrane voltage was recorded following the application of TTX.

### In utero electroporation

CAG-FLEX-Kir2.1-T2A-tdTomato and CAG-FLEX-Kir2.1Mut-T2A-TdTomato constructs were generated by amplifying Kir2.1-T2A-tdTomato from pCAG-Kir2.1-T2A-tdTomato (Plasmid #60598, Addgene) and mKir2.1-T2A-tdTomato from pCAG-Kir2.1Mut-T2A-tdTomato (Plasmid #60644, Addgene) (Xue et al., 2014). Using Gibson cloning, the fragments were inserted into pAAV-CAG-Flex-mRuby2-GSG-P2A-GCaMP6s-WPRE-pA backbone from which mRuby2-GSG-P2A-GCaMP6s-WPRE was removed.

In utero electroporations were performed in embryos of females with timed pregnancies, on E12.5. The mother was deeply anesthetized with isoflurane and buprenorphine (Temgesic, 0.05 mg/kg) was injected subcutaneously to the mother for analgesia. A 2 cm incision was performed at the abdominal midline to expose the embryos and 0.1 ml of Ritodrine (Sigma-Aldrich, 12 mg/ml) was injected to avoid premature abortion. 500 nl of Kir or mKir DNA (1 mg/ml) mixed with 1% Fast Green (Sigma-Aldrich) was injected into the lateral ventricle of the brain with a nano-injector (Nanoliter 2020, WPI). For the electroporation, five square pulses of 35V were delivered through the placenta at 950 ms intervals using a 3 mm tweezertrode (BTX Apparatus) driven by an electroporator. Following electroporation, the embryos were place back within the abdomen of the dam, the surgical incision was sutured. Embryos were extracted for immunohistological staining at E18.5, following the protocol described under “Immunohistological staining of embryonic tissue” below.

### Pharmacological manipulations in combination with in vivo para-uterine imaging

For the application of pharmacological agents during either calcium imaging or electrophysiological recordings (concurrent with imaging), a pipette, pulled from borosilicate glass with filament (O.D.: 1.0 mm, I.D.: 0.5 mm) using a P-97 micropipette puller (Sutter Instrument Company), was used.

Pipette were filled with a pharmacological agent, diluted with cortex buffer (as above). The two agents used were a mixture of AMPA (5 mM) (Cat no. 0254, Tocris) and NMDA (10 mM) (Cat no. 0114, Tocris), and TTX (1 mM) (Cat no. 1069, Tocris). In control applications, only cortex buffer was added to the pipette, without any pharmacological agent.

Embryos were prepared, as described for electrophysiological recordings, above. The cut coverslip allowed access of the pipette to the surface of cortex. The pipette was advanced diagonally until the tip was close to but not touching the surface of the brain, as monitored visually via brightfield imaging through the two-photon microscope.

Calcium imaging and electrophysiology were performed from Rbp4-Cre neurons under the coverslip, within 250 µm of the application site.

The pipette was connected to a syringe via silicone tubing, to allow the application of pressure to the fluid in the pipette. The meniscus of the fluid within the pipette was monitored visually during the application of pressure, to ensure that the fluid was delivered to the embryo (DMK 22BUC03, Imaging Source). In total, 500 nL of fluid was added into the buffer between the coverslip and the brain. In calcium imaging experiments using TTX, we made recordings immediately prior to and following the application of TTX (or cortex buffer, in control experiments) to the surface of cortex. In electrophysiological experiments using TTX, we patched a cell prior to TTX application, and took recording both before and after TTX application. In calcium imaging experiments using AMPA+NMDA, we performed 10-minute recordings, and applied AMPA+NMDA (or cortex buffer, in control experiments) to the surface of cortex, 3 minutes into each recording.

### Surgical isolation of embryonic cortex for single cell RNA sequencing

Dams, pregnant with embryos between E14.5 and E18.5, were anesthetized with a mixture of Fentanyl-Medetomidine-Midazolam (Fentanyl (Janssen, 0.05 mg/kg), Medetomidine (Virbac, 0.5 mg/kg); Midazolam (Sintetica, 5 mg/kg)). Dams were placed on their back and the hair on the abdomen was shaved. A 1.5 cm long incision was placed along the midline of the abdomen. The uterine wall was surgically opened longitudinally. A single embryo, within its yolk sac, was then isolated and the yolk sac was opened. The embryo was removed and placed in ice-cold oxygenated brain slice solution (sucrose (10 mM), trehalose (125 mM), KCL (3 mM), NaH_2_PO_4_ (1.25 mM), HEPES (26 mM), Dextrose (20mM), adjusted to pH 7.3). The brain was isolated, and the embryonic cortex removed. The posterior two-thirds of both cortical hemispheres were dissociated. 10 mL of papain activation solution was prepared (0.5 M EDTA, 100 mM L-Cysteine, 100 mM 2Me-EtOH) 30 minutes before the dissociation. 20 - 40 µl of papain solution (LS003126, Worthington) was then activated with 100 - 200 µl of papain activation solution, as per the manufacturer’s instructions. Each sample of dissected embryonic cortex was transferred into 1.5 mL Eppendorf tubes, cut into smaller square pieces of 1 mm^2^, and incubated in 1 ml of activated papain solution for 40 - 45 minutes at 37℃. Dissociated cortical cells were spun down at 1300 rpm for 2 minutes to remove as much papain solution as possible. Cells were then washed three times with DMEM (D4947, Sigma-Aldrich), supplemented with GlutaMAX (35050061, Gibco) and DNase (2000 U/ml; D4263, Sigma). Cells were spun down, again, at 1300 rpm for 2 minutes, and then resuspended in 0.5 - 1mL DMEM GlutaMAX, to which B27 (12587-010, Gibco) was added. To remove debris, cells were filtered through a cell strainer (352235, Falcon).

Sorting was performed on a FACSAria Fusion Cell Sorter (BD Biosciences) using a 100 μm nozzle and a sheath pressure of 20 psi. After recording several thousand events, the gating strategy was set using the FACSDiva software (v8.0.2, BD Biosciences). Size and granularity of the population of events was determined using the forward and side scatter values, and gates were set to exclude cell debris, doublets, and clumps. A final gate was set to select cells with high tdTomato fluorescence using a 582/15 nm bandpass filter (example gating strategy shown in Fig. S1). Cell were sorted in single-cell drop mode into a tube containing DMEM without phenol red (D4947, Sigma-Aldrich), supplemented with GlutaMAX (35050061, Gibco) and B27 (12587-010, Gibco). Sorted cells were immediately processed for single cell RNA sequencing (as described below). In some cases, tdTomato-negative cells were also sorted, in order to aid in isolating Rbp4-Cre neurons (detailed below).

### Single cell RNA sequencing

Cellular suspensions (8,000 cells per lane) were loaded on a 10x Genomics Chromium Single Cell instrument to generate single-cell Gel Beads in Emulsion (GEMs). Single-cell RNA-Seq libraries were prepared using GemCode Single Cell 3′ Gel Bead and Library Kit according to the manufacturer’s manual (version CG000204_Rev_C). The Chromium Gene Expression version 3.1 (v3.1) kit was used for all samples. Reverse transcription of GEMs was performed in a Eppendorf Mastercycler X50s thermal cycler using 0.2 ml PCR tube strips (E0030124286, Eppendorf): 53 °C for 45 minutes, 85 °C for 5 minutes, held at 4 °C. After reverse transcription, the GEMs emulsion was broken and the single-stranded cDNA was cleaned up with DynaBeads MyOne Silane Beads (37002D, Life Technologies). cDNA was amplified using a Eppendorf Mastercycler X50s thermal cycler in 0.2 ml PCR tube strips using the following program: 98 °C for 3 minutes, cycled 12 × : 98 °C for 15 s, 63 °C for 20 s, 72 °C for 1 minute; then 72 °C for 1 minute, hold at 4 °C. Amplified cDNA product was cleaned up with the SPRIselect Reagent Kit (0.6 × SPRI; B23317, Beckman Coulter). Indexed sequencing libraries were constructed using the reagents in the Chromium Single Cell 3′ library kit v3.1 (PN-1000121, 10x Genomics), following these steps: (i) fragmentation, end repair and A-tailing, (ii) post fragmentation, end repair and A-tailing; double-sided size selection with SPRIselect Reagent Kit (0.6 × SPRI and 0.8 × SPRI), (iii) adaptor ligation, (iv) post-ligation cleanups with SPRIselect (0.8 × SPRI), (v) sample index PCR using the Chromium multiplex kit (PN-120262, 10x Genomics), and (vi) post-sample index double-sided size selection with SPRIselect Reagent Kit (0.6 × SPRI and 0.8 × SPRI). The barcode sequencing libraries were measured using a Qubit 2.0 with a Qubit dsDNA HS assay kit (Q32854, Invitrogen) and the quality of the libraries assessed on a 2100 Bioanalyzer (Agilent) using a high-sensitivity DNA kit (5067‒ 4626, Agilent). Libraries were sent to the EMBL Genomics Core Facility (Heidelberg) for sequencing. Libraries were loaded at 240 pM on an Illumina HiSeq4000 with 150 cycle kits using the following read length: 28 cycles Read 1, 8 cycles i7 index and 91 cycles Read 2.

### Immunohistological staining of embryonic tissue

Dams, pregnant with embryos between E13.5 and E18.5, were anesthetized with a mixture of Fentanyl-Medetomidine-Midazolam (Fentanyl (Janssen, 0.05 mg/kg), Medetomidine (Virbac, 0.5 mg/kg); Midazolam (Sintetica, 5 mg/kg)). Dams were placed on their back and the hair on the abdomen was shaved. A 1.5 cm long incision was placed along the midline of the abdomen. The uterine wall was surgically opened longitudinally. A single embryo, within its yolk sac, was then isolated and the yolk sac was opened. The embryo was removed from the yolk sac, decapitated and the brain was removed, while immersed in ice-cold cortex buffer. Brains were immediately transferred into 4% (wt/vol) paraformaldehyde (diluted in phosphate buffered saline (PBS)). Brains were kept at 4 °C, for 12 hours, and then washed for 24 hours in PBS at 4 °C. To improve antibody penetration, the tissue was then washed with PBS and cryopreserved by incubation in 30% (wt / vol) sucrose at 4 °C until the tissue sunk to the bottom of the tube. Three sequential freeze-thaw cycles (using dry ice) were applied, and the tissue was then stored at −80 °C until further processing.

After washing in PBS, brains were embedded in 3% agarose (SeaKem LE Agarose, Lonza). 150 μm thick vibratome sections (VT1000S vibratome, Leica Biosystems) were incubated for 24 hours in blocking buffer containing 10% (vol/vol) normal donkey serum (NDS) (S30-100ML, Merck), 0.5% (vol/vol) Triton X-100 (93443-100ML, Sigma), 1% (wt/vol) BSA (A4161-1G, Sigma) and 0.01% (wt/vol) sodium azide (S2002-25G, Sigma) in PBS. Primary antibodies were diluted in a buffer containing 3% (vol/vol) NDS, 1% (wt/vol) BSA, 0.01% (wt/vol) sodium azide and 0.5% Triton X-100 in PBS and incubated for 3 – 7 days at room temperature. The primary antibodies used in this study were: rabbit polyclonal anti-GFP (A-11122, Invitrogen), rat monoclonal anti-GFP (04404-84, Nacalai), goat polyclonal anti Nurr1/NGFI-Bbeta/NR4A2 (AF2156, R&D Systems), rat monoclonal anti CTIP2/BCL11B (MABE1045, Merck), rabbit anti-PSD95 (ab269863, Abcam); goat polyclonal anti-snap25 (ABIN1742235, antibody online); chicken polyclonal anti-Map2 (CPCA-MAP2, Encor Biotechnology Inc); mouse monoclonal anti-Neurofilament H (801602, BioLegend); rabbit anti-cleaved caspase-3 (ab2302, Abcam); rabbit polyclonal RFP antibody (anti-tdTomato) (600-401-379, Rockland). Secondary antibody incubation was performed for 24 hours at room temperature in the same buffer as the primary. The secondary antibodies used in this study were all from ThermoFisher unless otherwise stated: Alexa Fluor 488 donkey anti-rabbit IgG (heavy and light chains (H+L), A-21206); Alexa Fluor 568 donkey anti-rabbit IgG (H+L, A10042); Alexa Fluor 647 donkey anti-rabbit IgG (H+L, A-31573); Alexa Fluor 488 donkey anti-rat IgG (H+L, A-21208); Alexa Fluor 568 donkey anti-goat IgG (H+L, A-11057); Alexa Fluor 633 donkey anti-goat IgG (H+L, A-21082); Cy3-AffiniPure donkey anti-rat IgG (H+L, 12-165-153, Jackson ImmunoResearch Laboratories). The secondary antibody solution was replaced with a solution of Hoechst (10 μg/ml) (33342, ThermoFisher) diluted to 1:10000 in PBS for 10 minutes. Slides were then washed 3 times in PBS. Finally, coverslips (631-0974, VWR) were added onto brain slices embedded using ProLong Gold Antifade Mountant (P36982, ThermoFisher).

### In situ hybridization

Whole embryonic brains were dissected at E14.5 and E18.5 and then fixed in 4% (wt / vol) paraformaldehyde (28908, ThermoFisher) in PBS at 4 °C for 8 to 12 hours. The tissue was then washed with PBS and cryopreserved by sequential incubations in 10%, 20% and 30% (wt / vol) sucrose at 4 °C until the tissue sunk to the bottom of the tube. Embryonic brains were then embedded in Optimal Cutting Temperature (OCT) Compound (25608‒930, VWR), frozen on dry ice and stored at −80 °C. 35 µm coronal sections were cut (CryoStar NX70 Cryostat 957000, ThermoFisher), excluding the olfactory cortex, and placed on SuperFrost Ultra Plus GOLD slides (11976299, ThermoFisher). Slides were placed on a 60 °C hot plate for 1 to 2 hours for increased tissue adhesion and were then air dried at −20 °C for 2 to 4 hours, before being stored at −80°C until use.

The RNAscope Multiplex Fluorescent Reagent Kit v2 Assay (323100, ACD Bio-techne) was used to detect RNA in embryonic cortical cells according to the user manual (Section: “Fixed-frozen tissue samples”), with the exception of a shorter incubation period with Protease III. Slides were incubated with 5 drops of Protease III for 4 minutes at 40 °C to facilitate probe penetration into the cells while minimizing damage to the tissue. Combinations of 3 probes per slide were used in their respective channels (C1, C2 and C3). Probes used were RNAscope 3-plex Positive Control Probe-Mm (320881, Lot 20049A), RNAscope 3-Plex Negative Control Probe (320871, Lot 20006A), Mm-GCamP6s-01 (557091, Lot 20233B), Mm-Lhx2-C2 (485791-C2, Lot 20232C), Mm-Pou3f3 (441521, Lot 19315A), Mm-Reln (405981, Lot 20078A), Mm-Nxph4-C3 (489641-C3, Lot 20241D), Mm-Crym-C2 (466131-C2, Lot 20241D), Mm-Nfe2I3-C3 (486201-C3, Lot 20241D), Mm-Fam19a2-C2 (452631-C2, Lot 20232C), Mm-Cdh8 (485461, Lot 20176C), Mm-Zcchc12-C3 (524441-C3, Lot 20232C). Slides were stored overnight at room temperature after hybridization in 5x Saline Sodium Citrate (SSC; 20X stock solution containing 3 M sodium chloride and 300 mM trisodium citrate, pH 7.0). Opal 520 dye (FP1487001KT, Akoya Bio), Opal 570 dye (FP1488001KT, Akoya Bio) or Opal 690 dye (FP1497001KT, Akoya Bio) diluted 1:1500 in TSA buffer (provided with the Multiplex Fluorescent Reagent Kit v2) were used as a fluorescent labels. Before mounting the slides, DAPI solution (provided with the kit) was added to the slides for 1 minute. Coverslips (631-0974, VWR) were added onto brain slices embedded using ProLong Gold Antifade Mountant (P36982, ThermoFisher).

### Image acquisition for immunohistology and in situ hybridization

A spinning disc microscope (Axio Imager M2 upright microscope, Yokogawa CSU W1 dual camera T2 spinning disk confocal scanning unit, Visitron VS-Homogenizer on an Olympus IXplore Spin confocal spinning disc microscope system) was used to image immunohistologically-stained slides, using 20X (UPLSAPO20X, Olympus) and 40X (UPLSAPO40XS, Olympus) objectives. In situ hybridization slides were imaged using an Olympus FV3000-BX63L upright confocal laser scanning microscope, using 20X (UPLSAPO20X, Olympus) and 40X oil (UPLANFL N40X, Olympus) objectives. The image acquisition settings were adjusted to maximally fill the available imaging dynamic range. Data collection and analysis were not blind to the conditions of the experiments.

### Whole mouse brain fixation and slice preparation for electron microscopy

Dams, pregnant with embryos at either E14.5 or E18.5, were anesthetized with a mixture of Fentanyl-Medetomidine-Midazolam (Fentanyl (Janssen, 0.05 mg/kg), Medetomidine (Virbac, 0.5 mg/kg); Midazolam (Sintetica, 5 mg/kg)). Dams were placed on their back and the hair on the abdomen was shaved. A 1.5 cm long incision was placed along the midline of the abdomen. The uterine wall was surgically opened longitudinally. A single embryo, within its yolk sac, was then isolated and the yolk sac was opened. A single embryo, within its yolk sac, was then isolated and the yolk sac was opened. The embryo was removed from the yolk sac, decapitated and the brain was removed, while immersed in ice-cold cortex buffer. Brains were immediately transferred into 4% (wt/vol) paraformaldehyde (diluted in phosphate buffered saline (PBS)). Brains were kept at 4 °C, for 12 hours, and then washed for 24 hours in PBS at 4 °C. Brains were fixed overnight at 4°C in freshly prepared phosphate buffer (0.1 M, pH 7.4) containing 4% paraformaldehyde (15700, Electron Microscopy Sciences) and 0.2% glutaraldehyde (16200, Electron Microscopy Sciences). After fixation, brains were embedded in 3% agarose (SeaKem LE Agarose, Lonza) and vibratome-sectioned (VT1000S vibratome, Leica Biosystems) into 60 µm thick slices. Slices were placed into multi-well culture dishes containing phosphate buffer saline (PBS) or phosphate buffer (0.1 M pH 7.4) containing 1% paraformaldehyde for longer storage.

### Immunostaining for immuno-electron microscopy

As in (Knott et al., 2009), to increase penetration of the antibody, 2 freeze/thaw cycles were applied (using liquid nitrogen). Slices were thawed in a cryoprotectant solution containing 2% vol/vol glycerol (G5516, Sigma-Aldrich) and 20% vol/vol dimethyl sulfoxide (DMSO) (D8418, Sigma-Aldrich) diluted in PBS. Slices were then agitated in a solution of 0.3% peroxide (107209, Merck) diluted in PBS for 10 min, and then washed three times in PBS/BSA-c (900.022, Aurion). Slices were incubated overnight, at 4 °C, with an anti-GFP primary antibody (AB3080, Chemicon) diluted 1:500 with PBS/BSA-c. Slices were then washed three times in PBS/BSA-c for 5 min before incubation for 90 min with a secondary antibody (Biotin-SP-affinipure goat anti-rabbit IgG, F(ab’)2 (111-065-047, Jackson Immuno Research Laboratories)) diluted 1:300 in PBS/BSA-c. After three washes, 5 min each, in tris-buffered saline (TBS: 0.1 M, 0.9% NaCl, pH 8) slices were treated with a peroxidase-based enzymatic detection system (Vectastain Elite ABC kit (PK-6100, Vector Laboratories)). Slices were then washed three times in TBS (0.05 M, 0.9% NaCl, pH 7.6) (T5030, Sigma-Aldrich), before incubation for 20 minutes, at room temperature, with a mixture of 0.04% (wt/vol) 3,3-diaminobenzidine (DAB) (D8001, Sigma-Aldrich) and 0.015% (vol/vol) hydrogen peroxidase (107209, Merck) diluted in TBS. Staining was terminated by rinsing the slices three times in TBS for 5 minutes each.

### Transmission electron microscopy (TEM)

As in (Knott et al., 2009), slices were transferred into glass scintillation vials containing cacodylate buffer (0.1 M, pH 7.4) (C0250, Sigma-Aldrich) and washed three times before post-fixation for 40 minutes with 1.0% osmium tetroxide (19110, Electron Microscopy Sciences) diluted in cacodylate buffer (0.1 M, pH 7.4). After a further three washes in ddH2O, slices were incubated in 1.0% (wt/vol) uranyl acetate dissolved in ddH2O. The slices were then rinsed three times in ddH2O and dehydrated through a graded ethanol series. After the final dehydration in 100% ethanol, slices were incubated twice in 100% propylene oxide (82320, Sigma-Aldrich). Slices were further infiltrated with a 1: 1 mixture of Durcupan resin (Resin A/M (44611, Sigma-Aldrich); B hardener (44612, Sigma-Aldrich); D hardener (44614, Sigma-Aldrich); DMP 30 (13600, Electron Microscopy Sciences)) and 100% propylene oxide, before being infiltrated with pure Durcupan resin. Finally, slices were flat embedded and cured overnight at 60°C. Thin sections of 50 nm each were cut using a Leica EM UC7 ultramicrotome and transferred onto formvar support film copper slot grids (FF2010-CU, Electron Microscopy Sciences) for TEM imaging.

Images were recorded at a magnification of between 8200X and 9900X for overviews and at a magnification of 16500X for imaging from immunolabeled synapses (corresponding to a pixel size of 2.8 nm) using a Tecnai Spirit TEM (FEI, Eindhoven Company) operated at 120 kV using a side-mounted 2K × 2K CCD camera (Veleta, Olympus).

### Imaging in adult layer 5 pyramidal neurons

PHP.eB AAV-CAG-FLEX-GCaMP7s, generated from the plasmid pGP-AAV-CAG-FLEX-jGCaMP7s-WPRE (Plasmid #104495, Addgene) (Dana et al., 2019) was produced as previously described in (Jüttner et al., 2019). For systemic administration of AAVs, Rbp4-Cre mice were anesthetized with 1.75% isoflurane. 0.5 – 20 μl of purified AAV, with the volume adjusted to 50 μl by adding saline solution (0.9%), was injected retro-orbitally into the sinus using a 30-gauge micro-fine insulin syringe (Yardeni et al., 2011). A minimum of 2 × 10^10^ genome copies (GC) of virus were injected per gram of mouse weight. Animals were imaged at least 3 weeks following AAV infection.

For implantation of cranial windows, mice were anesthetized with Fentanyl-Medetomidine-Midazolam (FMM) (Fentanyl (Janssen, 0.05 mg/kg), Medetomidine (Virbac AG, 0.5 mg/kg), Midazolam (Sintetica, 5 mg/kg)). To prevent dehydration of the cornea during surgery, we applied Coliquifilm (Allergan) to the eyes. The skin was removed, the skull cleared of tissue, and a thin titanium holder was attached to the skull with dental cement (Superbond C&B) allowing for head fixation during calcium imaging (Holtmaat et al., 2009). A 4 mm diameter craniotomy was made over the left hemisphere, exposing the visual cortices and surrounding areas. After removal of the bone, dehydration of the cortical surface was minimized by repeatedly applying cortex buffer to the surface. The cortical surface was covered with a 4 mm diameter glass coverslip and sealed with UV glue (NOA 68, Norland), stabilized by an additional layer of dental cement. Buprenorphine (Temgesic, 0.05-0.1 mg/kg) was injected 20 minutes before the end of the surgery to provide extended pain relief during the immediate recovery from surgery. At the end of surgery, mice were woken using an antagonist mixture to counteract FMM anesthesia (Atipamezol (Virbac, 2.5 mg/kg), Flumazenil (Sintetica, 0.5 mg/kg)). Mice were monitored daily following surgery and allowed to recover for at least 10 days before any recordings.

Mice were head-fixed, in darkness, within the Femtonics galvo-galvo scanning two-photon setup, and Rbp4-Cre neurons were imaged under the Nikon 25X water immersion objective (1.1 NA). Imaging was performed at 5-10 Hz. The resulting recordings (used to generate Fig. S3) were then analyzed as with embryonic recordings, as detailed below.

### In vivo para-uterine blood flow imaging

Dams were anesthetized and maintained at a surgical depth, as assessed by the lack of spontaneous movement, as well as the absence of a paw withdrawal reflex, in response to a toe pinch. To maintain the anesthesia, top-up doses of FMM anesthesia (10 – 30% of the initial bolus) were injected into the dam, between recordings, approximately every 30 minutes. Throughout the experiment, a nasal mask was used to deliver humidified oxygen to the dam, at a flow rate of 0.8 l/min (Respironics EverFlo OPI pump, Phillips). Dams were monitored visually throughout blood flow imaging sessions. In addition, a pulse oximeter attached to a sensor applied to the rear paw (MouseSTAT Jr and Paw Sensor, Kent Scientific) was used to monitor the heart rate and blood oxygenation of each mouse during experiments. For all dams, the heart rate was maintained at less than 400 bpm (Fleischmann et al., 2016), and the blood oxygenation never dropped below 90%, when averaged over a minute.

Embryos were imaged under a widefield microscope (SZX16, Olympus) in the brightfield, to observe blood flow throughout the imaging region at the beginning of the experiment, after 5 hour and, 10 minutes following disruption of the umbilical cord (severing of the cord). Recordings were collected using a CMOS camera (ORCA-Flash4.0 V3, Hamamatsu). Data was recorded via the CameraLink interface, using the HCImage Live software (Hamamatsu).

### Filtering of calcium traces

On the galvo-galvo scanning microscope, two-photon recordings were imported as 3D matrices of recorded activity within a spatial field of view over time. Similarly, recordings taken with high - speed arbitrary frame scan mode on the acousto-otpic two-photon microscope were also imported as 3D matrices across time, by taking the mean projection across all imaging planes, at each time point. Following image stabilization (Pnevmatikakis and Giovannucci, 2017), custom code (Matlab R2020a, Mathworks) was used for manual selection of elliptical regions of interest (ROIs) within the imaging field of view containing individual cells or dendrites. The distance between elliptical ROIs was quantified as the Euclidean distance between the centers of each of the two ellipses. The average of the pixels within the ROI at each time point was defined as the raw activity trace for each ROI. Three separate regions of background expression were also selected within each imaging field of view.

To estimate the per pixel fluorescence within individual neuronal somas as a function of the background activity in the embryonic tissue (as in Fig. S6), the raw activity traces were integrated over time and normalized by the number of pixels within each ROI, to obtain a raw calcium activity per pixel over the recording interval. The same raw pixel fluorescence value was computed for the three background regions selected within each imaging window. The amplitude of the cellular fluorescence as a function of the per pixel background fluorescence, for each background region, was computed within each imaging window, and compared against the distribution of background fluorescence (as a function of other background ROIs).

For recordings taken on the acousto-optic two-photon microscope in multi-cube scanning mode, the mean projection across imaging planes at each time point was computed to generate a 3D matrix for each ROI across time. Following image stabilization (Pnevmatikakis and Giovannucci, 2017), custom code (Matlab R2020a, Mathworks) was used for manually selecting a single ROI, containing the imaged soma or neurite, within each mean imaging field. To identify a responsive object within the manually selected ROI, custom code was used to identify pixels which responded with similar temporal profiles. The mean pairwise correlation of each pixel with its surrounding pixels over the recorded time interval was computed (2 concentric rings of surrounding pixels; Matlab R2020a, Mathworks). The correlation of each pixel within the ROI was compared against the distribution of pairwise correlations for pixels outside the manually selected ROI, within the same mean projected imaging plane. Pixels within the manually selected ROI that had a significantly higher mean pairwise correlation were identified as part of the recorded object (>2σ). If the fraction of significant pixels within the manually selected ROI was greater than 20%, the mean response of all such pixels was defined as the raw activity trace for each recorded object.

To denoise the raw traces, recordings were first converted to Fourier domain. A threshold, set at 150% of the mean amplitude component of the trace, was then applied to the amplitude component across the entire frequency range. This filter reduces white noise within the frequency spectrum, without a bias towards any specific frequency component. For recordings on the galvo-galvo microscope, the total power within the filtered frequency spectrum of the three background ROIs, was used as a secondary threshold. If an ROI contained a power less than the mean over the background ROIs’, the filtered trace was set to zero. In contrast, if the power of an ROI was greater than the background ROIs’ power, the filtered amplitude component was inverted back to obtain a denoised trace in time. To appropriately account for spontaneous activity, a %ΔF/F trace was then computed by utilizing the mean amplitude across the entire recording as the estimate of F. For all embryonic days following E13.5, a rolling mean with a time constant of 62.5 s was then subtracted away from the filtered activity trace, to ensure that the baseline was stable across the recording length. This trace was then used for all further analyses. For recordings on E13.5, a shorter rolling mean with a time constant of 12.5 s was subtracted away from the filtered activity trace. Finally, to obtain an estimate of the total activity within each ROI, the average of the %ΔF/F activity across time was used, yielding a unit of %(ΔF/F)/s.

While event-based analyses were also performed (see below), %(ΔF/F)/s was selected for most analyses in the paper since it provides an estimate of the overall calcium activity, and is less dependent on assumptions about the calcium kinetics of each individual event (as required to detect calcium events).

For recordings made to assess the effect of TTX application, recordings were filtered as detailed above. The total activity within each ROI was determined, in a 10-minute recording immediately prior to, and a 10-minute recording immediately following the injection of TTX.

For recordings made to assess the effect of AMPA+NMDA application, the 3D matrix was image stabilized (Pnevmatikakis and Giovannucci, 2017) and the same custom code as above (Matlab R2020a, Mathworks) was used for manual selection of elliptical ROIs within the imaging field of view. Calcium activity in each ROI, per unit area, was calculated by summing across all pixels within each ROI and then dividing by the size of the ROI. The baseline fluorescence was defined as the average fluorescence within each ROI prior to the application of AMPA+NMDA, i.e. across the first three minutes of each recording. The fluorescence change, as a fraction of the baseline, was then computed and the change in the first minute of each recording (i.e. two minutes before the application of AMPA+NMDA) was compared to the change in the 5^th^ minute of the same recording (i.e. two minutes following the application of AMPA+NMDA) (as in Fig. 6B).

### Computing event-based statistics

Calcium events were defined as fluctuations of ΔF/F above baseline with a length greater than 1.5s and a minimum average activity of at least 10 %(ΔF/F)/s. The size of each event is defined as the total ΔF/F recorded from start to end. The frequency of events in each ROI was computed by dividing the total number of events by the total length of the recording. The properties of the calcium events recorded are examined in Fig. S8.

### Quantifying voltage changes in response to current steps

Every embryonic neuron, independent of recording day, responded with a maximum of a single nonlinear depolarization, at each current step, which was always located close to the initiation of the current injection. For the voltage curve recorded in response to a single current injection, the peak voltage in the first half of the current injection was defined as the peak voltage, and the mean voltage across the second half of the current injection was defined as the steady state voltage. Any active conductance was thereby represented by a peak voltage that was higher than the steady state voltage.

### Quantifying excitatory synaptic potentials

Excitatory synaptic potentials were computed from subthreshold membrane voltage recorded in the presence of TTX. Recordings, taken at 50000 Hz, were downsampled by averaging recorded values in 50 frames bins, to generate traces with a frequency of 1000 Hz. A rolling mean with a time constant of 1s in length was subtracted away from the downsampled traces, to ensure that the baseline was stable across the recording length. We termed this trace the membrane voltage with adjusted baseline, and it was used for all further analyses.

Visual inspection of the amplitude distribution of the membrane voltage with adjusted baseline demonstrated a number of peaks within a long tail of amplitude values, suggesting the presence of events of quantal size. A threshold of 5 mV was chosen, and for each event crossing this threshold, a window of 100 ms prior and 400 ms following the event was selected. This 500 ms interval was then averaged across events to obtain the average shape of an excitatory event within Rbp4-Cre neurons (separately at both E14.5 and E18.5).

### Location of cells within immunohistological sections

The bottom edge of the subplate (defined by the lower border of staining with CTIP2 (Bcll1b) and Nurr1 (Nr4a2) antibodies), the top edge of the cortical plate (defined by the outer edge of Hoechst staining), as well as the center of individual Rbp4-Cre neurons (defined by a GFP antibody) within coronal sections of the embryonic brain were manually annotated. A custom macro for Fiji (Schindelin et al., 2012) (modified from the points_to_curve_distance macro) was used to compute the perpendicular distance of each cell to the two edges. The fractional distance of Rbp4-Cre neurons from the bottom of the subplate, normalized by the total distance within the cortical plate and subplate was computed, on all embryonic days. The fractional locations for the set of Rbp4-Cre neurons across all embryonic days was clustered into three clusters by hierarchical clustering of the mean of the pairwise Euclidean norms. The boundaries of these clusters were used to define the fractional edges of each of the three layers of Rbp4-Cre neurons, within the embryonic cortex.

### Quantifying in vivo para-uterine imaging of anatomy

Due to movement between the imaging of each plane, custom code was used to register images using a rigid transformation, within each 3D stack, to a reference plane, and the registered stacks output as TIFF stacks (Matlab R2020a, Mathworks). Fiji (Schindelin et al., 2012) was then used to rotate the TIFF stacks into a projection onto the xy plane, thereby showing the location of cells within the cortical plate and subplate. Individual cells were selected that were identifiable across all time points. As with the location of cells within immunohistological sections, the bottom edge of the subplate, the top edge of the cortical plate, as well as the center of individual Rbp4-Cre neurons within coronal sections of the embryonic brain were manually annotated. A custom macro for Fiji (Schindelin et al., 2012) (modified from the points_to_curve_distance macro) was used to compute the perpendicular distance to the two edges. The fractional distance of Rbp4-Cre neurons from the bottom of the subplate, normalized by the total distance within the cortical plate and subplate was computed, and neurons classified into the individual layers using the quantitative boundaries defined from immunohistological staining. The average location of neurons within different anatomical layers could then be quantified across time.

### Quantification for in situ hybridization

All GCaMP6s-positive cells per image in cortex were manually identified. GCaMP6s-positive cells were then assessed for colocalization of markers for E-NP (Nfe2I3, Nxph4, Zcchc12, and Reln (on E14.5 only)), E-IT (Lhx2 and Fam19a2) and E-PT (Cdh8 and Pou3f3 (on E18.5 only)). GCaMP6s-positive neurons were identified by their anatomical location as being members of a specific layer (upper and lower at E14.5; middle and lower at E18.5). The fraction of GCamP6s-positive cells in each layer that were positive for each marker was then calculated on both embryonic days. Finally, the fraction of all GCaMP6s-positive cells positive for at least one of the markers for each type was computed.

### Quantification of blood flow rate

Brightfield recordings of blood vessels, at the surface of the embryonic cortex, were obtained at 100 Hz and were imported as 3D matrices of the 2D spatial field of view over time. 3D matrices were stabilized (Pnevmatikakis and Giovannucci, 2017), and identical blood vessels identified by visual inspection of the pattern of blood vessels, at the start of imaging, after 5 hours of imaging, and following the disruption of the umbilical cord. Custom code (Matlab R2020a, Mathworks) was used to draw cross-sections across a subset of the blood vessels. The amplitude of activity averaged across the pixels within the cross-section of the blood vessel was computed. This amplitude within each blood vessel was filtered by the application of a rolling mean with a time constant of 25 ms, followed by the subtraction of a rolling mean with a time constant of 3s, to denoise the recorded signal and ensure that the baseline was stable across the recording length.

The blood flow within each blood vessel results in changes in amplitude as blood cells pass the cross-section. Therefore, we quantify the number of fluctuations in amplitude across the recorded interval. We computed the fractional change in the number of fluctuations from the recording at the start of imaging to the recording after 5 hours, and the fractional change in the number of fluctuations from the recording after 5 hours to the recording 10 minutes following the disruption of the umbilical cord, per blood vessel.

### Correlation analyses

The correlation of the spontaneous activity within Rbp4-Cre neurons in each recording was computed as the pairwise Pearson correlation of the denoised activity in each pair of neurons. To determine that the observed correlations were greater than that expected from a random pair of neurons with the same overall activity, the spontaneous activity traces for each pair of neurons were reordered, 1000 times. In detail, individual events within each spontaneous activity trace (defined as deviations of the ΔF/F > 0, as introduced above) were shuffled in time, to maintain both the overall spontaneous activity level as well as the time constant of individual events (which differed across embryonic days). The distribution of pairwise correlations of both neurons’ activity across the 1000 randomized pairs defines the random distribution for each pair of neurons, and neuron pairs showing significant pairwise correlations were those with correlations significantly higher than this distribution (>3σ). Note that this distribution is specific to the level of activity within each pair of neurons. To obtain a model of the random correlations across all pairs, the distribution of correlations across all pairs of neurons that were not significantly different from their specific random distribution was used.

To calculate the synchrony across a population of Rbp4-Cre neurons within a recording (as for Fig. S24), we computed the pairwise Pearson correlation between activity within each soma (filtered as described above) and the sum of the activity in all other somas within a given recording, i.e. excluding the region of interest for which the synchrony is being computed (Matlab 2020a, Mathworks) (Bharioke et al., 2022).

### Single-cell RNA sequencing alignment and pre-processing

Analysis was performed in Python (v3.7.7). The workflow for the analysis was managed using Snakemake (v5.19.3). Sequencing results were aligned to a reference and barcodes identified using StarSolo (v2.7.5a). The reference was generated from the mouse genome (GRCm38) and annotation (GRCm38.93) and modified to include the sequences for tdTomato (Addgene plasmid # 22799) (Madisen et al., 2010) and the Cre recombinase present in the mouse strain (personal communication with S. Gong) (Gerfen et al., 2013; Gong et al., 2007).

Raw count matrices for each sample were loaded into and primarily handled within ScanPy (v1.5.1). To filter cell barcodes associated with empty droplets, count matrices were temporarily transferred to R (v3.6.1) using rpy2 (v3.3.2) and EmptyDrops (Lun et al., 2019) was applied using DropletUtils (v1.6.1). Droplets with a false-discovery rate less than 0.001 were classified as non-ambient and retained for subsequent steps.

To identify multiplets (cell barcodes associated with multiple cells), we used Scrublet (v0.2.1) to simulate the creation of multiplets from our data and score our observed cells in comparison (Wolock et al., 2019). The threshold separating likely multiplets from individual cells was estimated using a mixture of two Gaussians fit to the simulated scores, visual inspection confirmed that one Gaussian captured the singlet distribution and the other Gaussian captured the multiplet distribution. The point where these two distributions crossed was used as the threshold beyond which cells were excluded as being more likely a multiplet than a singlet. The process of simulation, identifying the threshold, and excluding cells beyond the threshold was repeated for each sample individually.

Cells were filtered as low-quality if they expressed fewer than 500 distinct genes or if the fraction of their reads originating from mitochondria exceeded 3 standard deviations from the median fraction of mitochondrial reads within the sample. To reduce the influence of outliers, median absolute deviation was used to estimate the standard deviation. Mitochondrial and ribosomal genes were filtered out of most subsequent analyses.

Count matrices across all samples were aggregated, and a Leiden clustering (leidenalg, v0.8.0) was performed (Traag et al., 2019). Unless otherwise specified, gene expression is presented as log_2_ pseudocounts of transcripts which were first normalized to a total library size of 10,000 in each cell. Three individual cells were selected at random from every cluster and manually annotated by a trained observer according to their individual transcriptomes. Additional cells were annotated when necessary. In total, 106 cells were annotated in this manner. Throughout the process, an interactive cell browser was used to assist experimenters in exploring the data (UCSC Cell Browser v0.7.11).

### Isolation of Rbp4-Cre neurons

When enriching for tdTomato-labelled cells (Rbp4-Cre neurons) with FACS, a relatively permissive gate was used to help ensure a representative set of labelled cells. Additional enrichment of the sequenced single cells was performed to further isolate Rbp4-Cre neurons (Fig. S1). First, clusters annotated as excitatory neurons or their progenitors were selected (a total of 25681 cells). Second, from these excitatory neurons, communities of cells with high tdTomato were located and isolated. Community detection was performed by applying a weighted kernel density smoothing to a UMAP embedding (60-nearest-neighbor graph, top 400 overdispersed genes) (Becht et al., 2019) where the weights are the tdTomato transcript count in each cell and the kernel bandwidth is half of the bandwidth determined by Scott’s rule. This approach was selected because (i) it is more robust to dropout and ambient RNA than applying a threshold at the single-cell level (ii) it may provide a more-principled cutoff for inclusion for the lineages contained in our data than clustering.

To validate this method we used the same kernel and UMAP projection to compare communities of cells identified as having high tdTomato expression with communities of cells which FACS selected as possible Rbp4-Cre neurons. The degree of FACS enrichment was estimated by comparing the position on the UMAP plot of cells which were positive- and negative-selected by FACS and sequenced separately by dividing the kernel density of the positive-sorted cells by the density of the negative-sorted cells at each cell position. The highest FACS enrichment was observed in communities of cells containing the highest tdTomato expression (Fig. S1).

The threshold for a cell to be included as tdTomato-labelled was 6 estimated standard deviations above the median kernel-smoothed tdTomato expression (Fig. S1). To avoid the influence of outliers, this threshold was determined using cells which had a FACS enrichment ratio of less than 1 and by estimating the standard deviation using the median absolute deviation. Following this enrichment, we obtained a population of 1098 positive-identified Rbp4-Cre neurons, which were used for all further analyses.

### Embedding and clustering of Rbp4-Cre neurons

Embeddings of Rbp4-Cre neurons were created by applying UMAP to a 60-nearest neighbor graph created directly from the top 400 overdispersed gene representation (Becht et al., 2019). Leiden clusters were calculated at a resolution of 0.2 (Traag et al., 2019).

### Statistical analysis of single cell RNA sequencing in embryonic Rbp4-Cre neurons in relation to the adult cortex

RNA sequencing data from single cells of the adult cortex were loaded as counts of reads mapping to exons (from (Tasic et al., 2018)), normalized by the effective gene length to approximate the transcript abundance, and then library-normalized to a total of 10000 normalized counts per cell. Subsequent comparisons were performed in log_2_ space using pseudocounts.

To compare gene expression of Rbp4-Cre neurons to that of cells from each adult layer, we identified the top genes discriminating the different adult layers using Wilcoxon rank sum. The number of genes selected was varied to verify that results were robust. Spearman correlations of gene expression were calculated between the Rbp4-Cre neurons and each of the adult layers.

Individual Rbp4-Cre neurons were compared to types of adult layer 5 cells using a Bayesian approach. Starting with a uniform prior, the posterior probability of each single cell was updated based on the genetic identity of the transcripts observed inside it and the conditional probability of observing transcripts with those genetic identities in each of the three adult layer 5 cell types. The joint distribution between genes and cell types in the adult was pre-processed in two ways: (i) additive smoothing was applied by adding 1 to the number of transcripts observed in each combination of adult cell type and gene before normalizing to create the joint probability distribution and (ii) the joint probability distribution was normalized such that each gene had the same marginal probability of being observed. The gene space employed for these comparisons was the intersection of the top 600 genes differentially expressed between the adult types and different numbers of genes differentially expressed by the clusters of Rbp4-Cre neurons. The fraction of cells which had a posterior probability greater than 99% (Bonferroni-adjusted for multiple comparisons) of being sampled from one of the three adult layer 5 cell types revealed that the cell type association of clusters was not sensitive to the number of genes selected within a range of 7 to 40 unique genes.

The ternary plot was created using the posterior probability of each Rbp4-Cre neuron being one of the three adult cell types, using the set of 24 unique genes, chosen as discussed above. To prevent the majority of cells from overlapping each other at the extreme corners of the triangles, the values presented on the ternary plot were curved by taking the 15th root of each cells’ probabilities followed by normalization to a total of one.

### Distribution of gene expression in Rbp4-Cre neurons

Genes associated with synaptic transmission, signaling, the synaptic vesicle cycle, and active membrane properties were identified from appropriate Kyoto Encyclopedia of Genes and Genomes (KEGG) pathways including *Glutamatergic synapse*, *GABAergic synapse* (postsynaptic), and *Synaptic vesicle cycle* (Kanehisa and Goto, 2000). Genes associated with transsynaptic signaling were identified from a review of synaptic development (Siddiqui and Craig, 2011). Spearman correlation was used for comparing correlations of transcripts across days, within each of the embryonic cell types.

The list of schizophrenia associated genes was chosen from a genome wide association study of the disease (Ripke et al., 2014), the OMIM database ([CSL STYLE ERROR: reference with no printed form.]), and supplemented with genes found to be associated with schizophrenia in a genome wide association study of multiple neurodevelopmental diseases (Lee et al., 2019). To ensure a lack of bias in the representation, the list was not revised further, except for the removal of genetic loci and genes not annotated in our dataset (primarily because of the absence of a mouse ortholog), including: *Apol2*, *Apol4*, *Cacna1l*, *Chi3l1*, *ChrnaB4*, *Daoa*, *Disc2*, *Fam5b*, *Nlgn4x*, *Sczd1*, *Sczd11*, *Sczd12*, *Sczd2*, *Sczd3*, *Sczd5*, *Sczd6*, *Sczd7*, and *Sczd8*.

The list of autism spectrum disorder associated genes was chosen by ordering genes by their SFARI gene score (from 1 to 3) and, then secondarily, by the number of reports (Abrahams et al., 2013) (accessed as of August 29, 2020). In this ordering, all genes with greater than 10 reports were chosen. To ensure a lack of bias in the representation, the list was not revised further, except for the removal of genes that were not expressed in the adult data set (Tasic et al., 2018): *Kmt5b*, *Nexmif*, *Scn2a*.

For each gene, the mean number of transcripts across (i) the population of Rbp4-Cre neurons and (ii) the population of adult layer 5 neurons (combining NP, IT, and PT types in VISp) (Tasic et al., 2018) was quantified. Expression values for each gene were presented as a circle plot, separated by embryonic type and age. To compare expression of disease-associated genes to the set of all genes, in both embryonic Rbp4-Cre neurons and adult layer 5 neurons, the distribution of mean expression was separately computed for the subset of genes associated with each disease and for the full set of genes. Distributions of disease gene expression were compared to the distribution of the full set of genes using Wilcoxon rank-sum, with a p value below 0.05 (Bonferroni corrected for 6 comparisons). A significant difference indicated that the set of genes associated with a given disease had higher than expected expression compared to the full set. For each of the diseases, the fold difference in gene expression was compared between embryonic Rbp4-Cre neurons and adult layer 5 neurons. In each case, fold difference was computed by dividing the mean expression across genes associated with the disease by the mean expression across the full set of genes.

### Statistical analyses

All statistical comparisons of distributions, except for the statistical significance of pairwise correlations (discussed under correlations, above), the statistical significance in the analysis of gene expression profiles (discussed under distributions of gene expression, above), and the analysis of distribution of neurons into layers at E14.5 and E15.5 (for Fig. S13) were performed using either the Wilcoxon rank-sum test, for unpaired data, or the Wilcoxon signed rank test, for paired data. The significance threshold used for both tests was 3σ. The comparison of the spatial distribution of neurons into layers between GCaMP6s-tTA2 and tdTomato reporter mice (Fig. S14) was performed using a chi-squared test (χ^2^ test).

---

**Fig. S1. Associated with Fig. 1 and 4.**
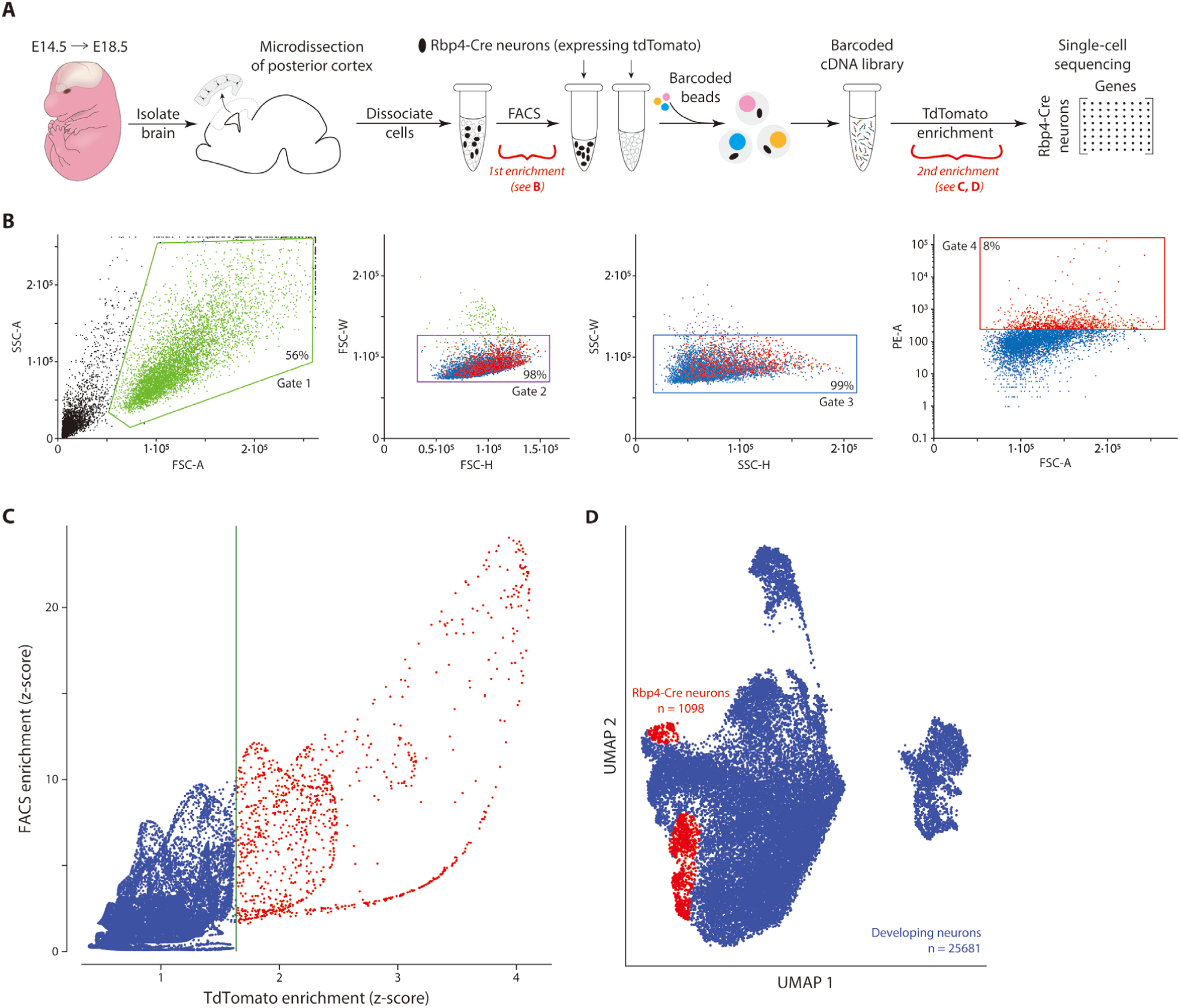
Single cell RNA transcriptomes from enriched population of tdTomato-expressing Rbp4-Cre neurons from dissociated embryonic cortical tissue. (**A**) Single cell RNA sequencing workflow overview demonstrating two distinct stages of enrichment for Rbp4-Cre neurons (red brackets). (B) Initial enrichment of Rbp4-Cre neurons was performed by isolating cells positive for the tdTomato marker from the dissociated cortical tissue using FACS. Gates were selected based on the size and granularity of sorted events. Gate 1 was chosen to exclude debris, while gates 2 and 3 were chosen to exclude doublets. Gate 4 was chosen to select cells with higher tdTomato fluorescence. Box: events filtered at each gate. (**C**) Additional enrichment for positively identified tdTomato-expressing Rbp4-Cre neurons. Relative enrichment of cells over a negatively FACS sorted population (blue) was used to define a baseline tdTomato expression level. Cells with significantly greater tdTomato expression than in the baseline were selected as positively identified Rbp4-Cre neurons (red). Green line: 6σ threshold of tdTomato expression. (**D**) UMAP embedding of all 25681 excitatory neuron transcriptomes (blue) demonstrating the location of the subset of 1098 positively identified Rbp4-Cre neurons (red).

**Fig. S2. Associated with Fig. 1.**
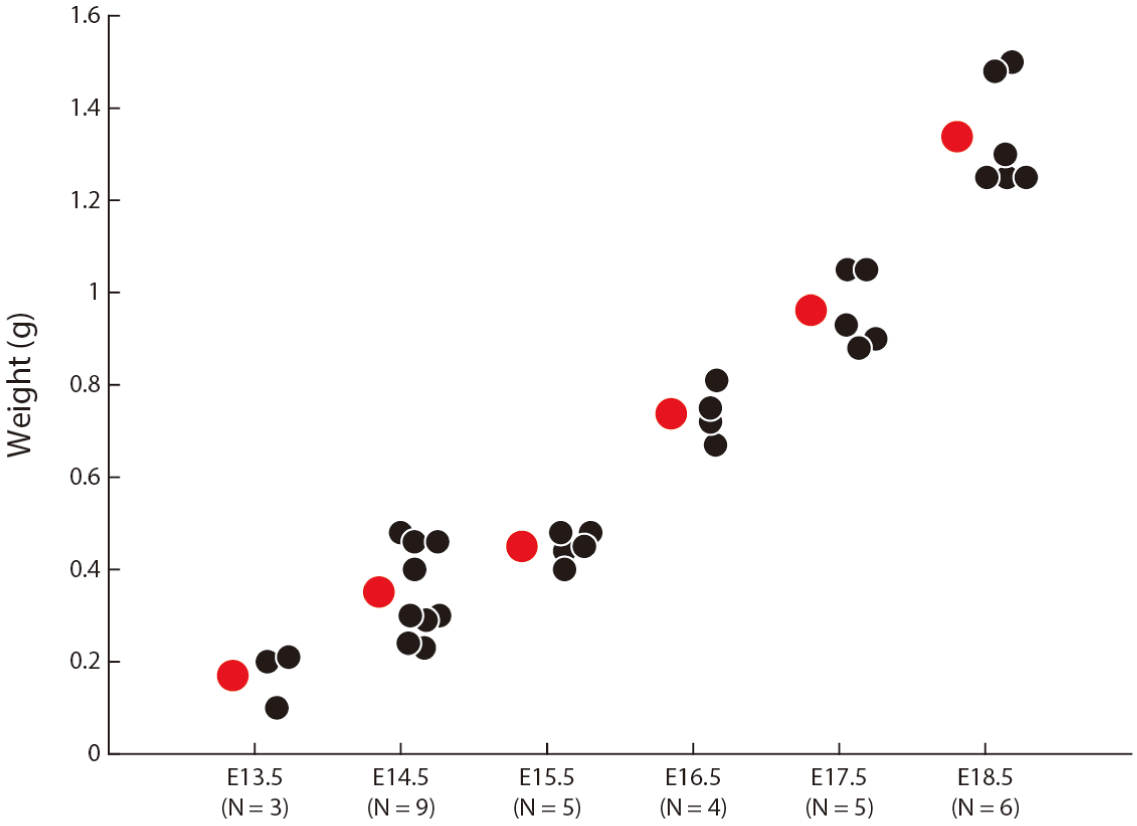
Embryonic weight on each embryonic day. Mean (red dot) of embryonic weights on each day (black dots) ranges from 0.17 g at E13.5 to 1.3 g at E18.5.

**Fig. S3. Associated with Fig. 1.**
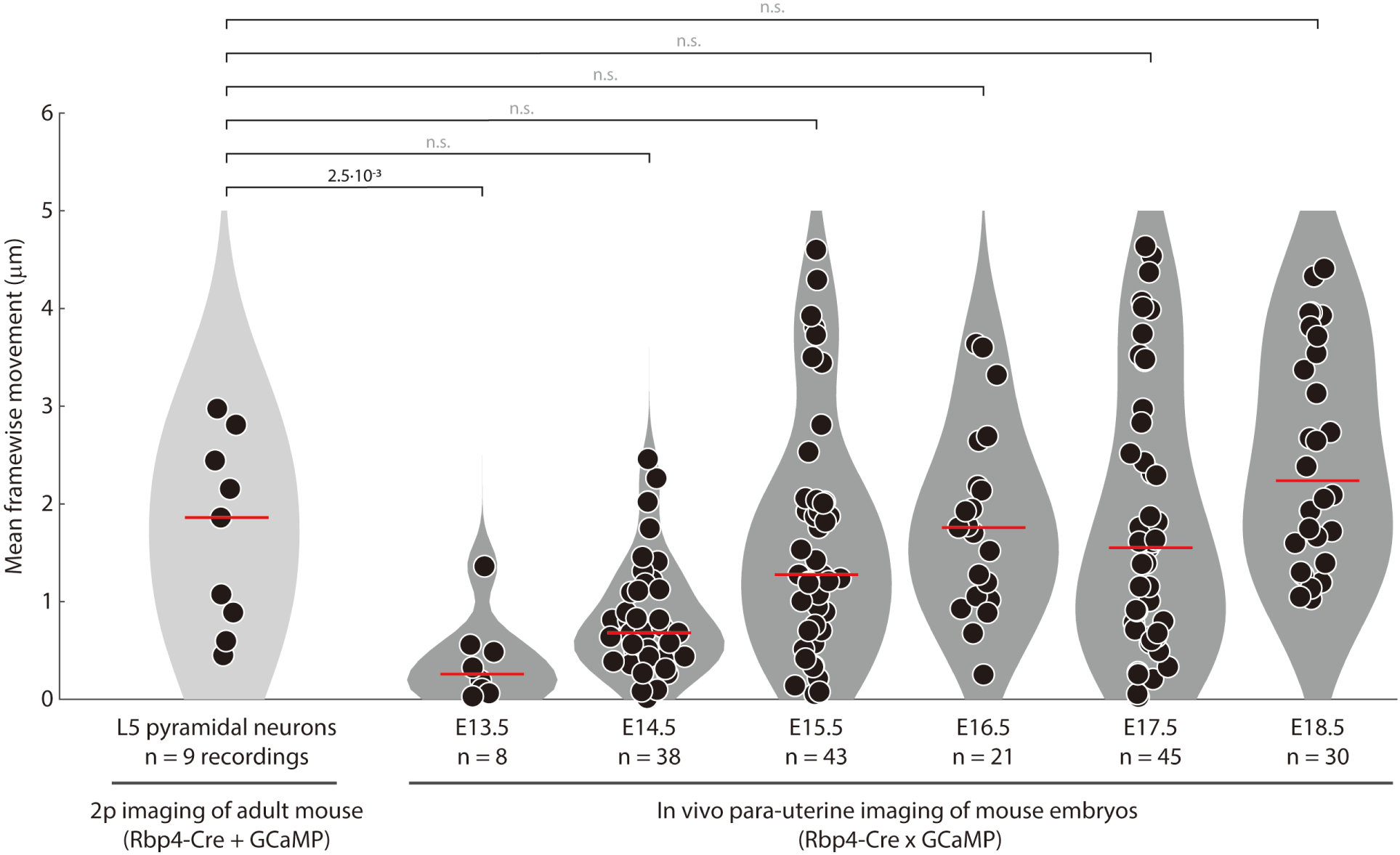
Stability of in vivo para-uterine two-photon imaging of cortical neurons in embryos is similar to stability of two-photon imaging of cortical neurons in adult mice. Movement per frame (recorded from 5 – 10 Hz), averaged per recording, computed via rigid motion correction between frames (Pnevmatikakis and Giovannucci, 2017). In vivo embryonic recordings were made as schematized in Fig. 1F. Adult recordings were made in head-fixed Rbp4-Cre mice, injected with AAV expressing cre-dependent GCaMP. Probability: Wilcoxon rank-sum test; n = number of recordings from 3 (E13.5), 9 (E14.5), 5 (E15.5), 4 (E16.5), 5 (E17.5), and 6 (E18.5) embryos and 3 adult mice.

**Fig. S4. Associated with Fig. 1.**
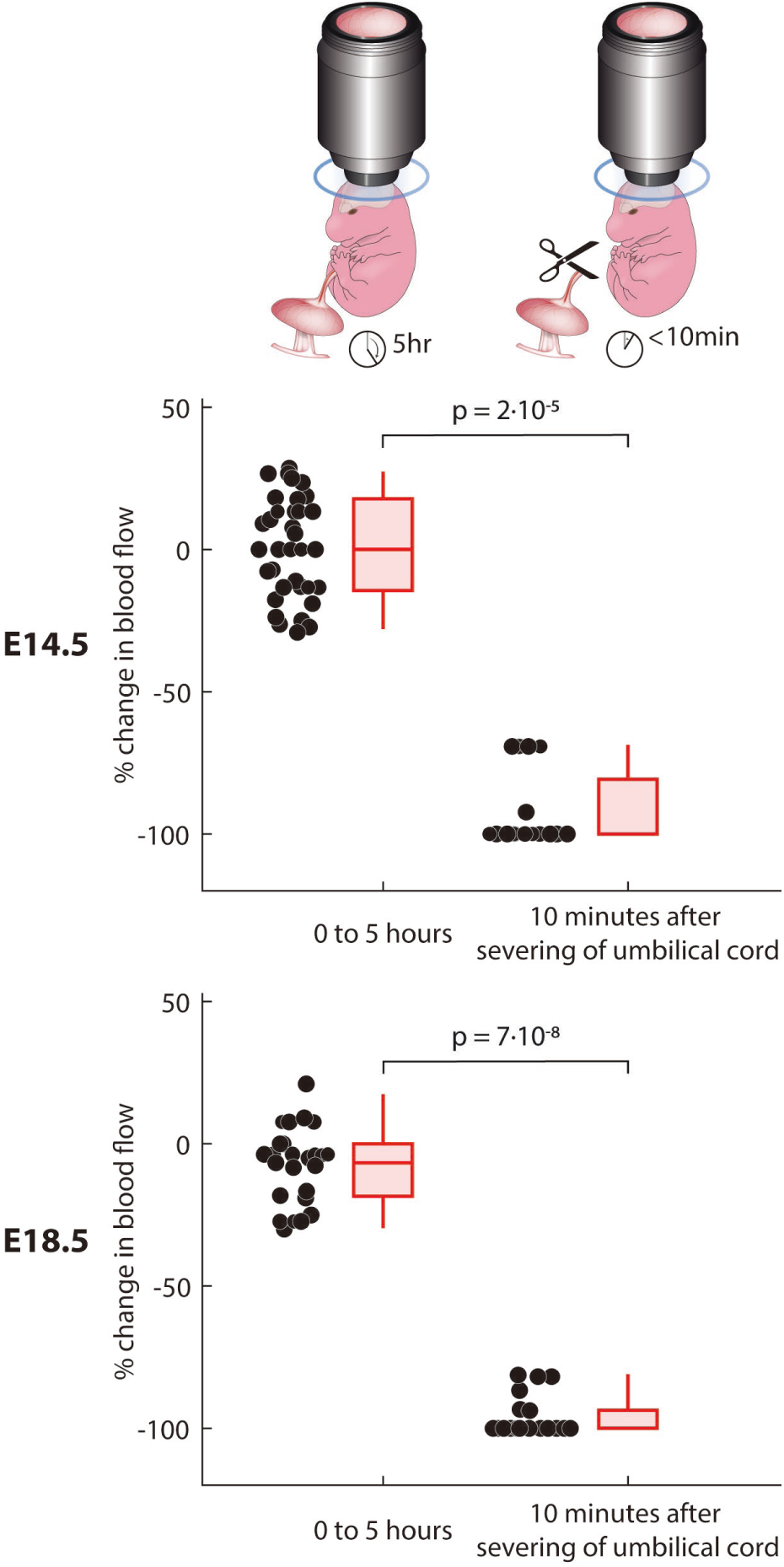
Embryonic blood flow does not change following 5 hours of imaging, but degrades rapidly following severing of the umbilical cord. Blood flow at the surface of the brain was imaged in visible light prior to and following 5 hours of imaging and the difference in blood flow is quantified (left), at E14.5 (top) and E18.5 (bottom). Blood flow at the surface of the brain was also imaged immediately prior to, and 10 minutes following the severing of the umbilical cord. The difference in blood flow is quantified (right), at E14.5 (top) and E18.5 (bottom). Box-and-whiskers: distribution of changes in blood flow across each time window, as box (25-75 percentile) and whisker (5-95 percentile); red lines: median. Probability: Wilcoxon rank-sum test.

**Fig. S5. Associated with Fig. 1.**
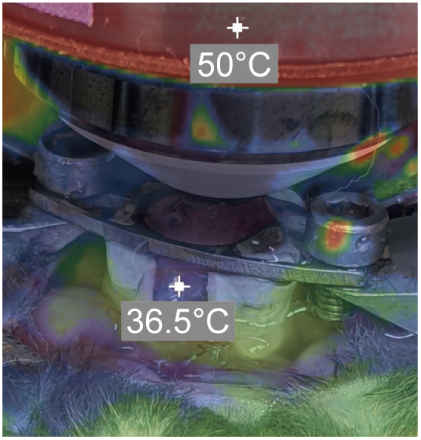
Temperature of embryo stabilized para-uterine under the two-photon microscope. Infrared image is aligned with a visible light image, where the embryo can be observed within the holder (as schematized in Fig. 1F). The 36.5°C marker labels the embryo (Refinetti, 2010; Reitman, 2018), which is visible through the opening in the holder allowing for the exit of the umbilical cord. The second 50°C marker labels the objective heater, providing a secondary source of heat during imaging. Image was taken following 5 hours of imaging.

**Fig. S6. Associated with Fig. 1.**
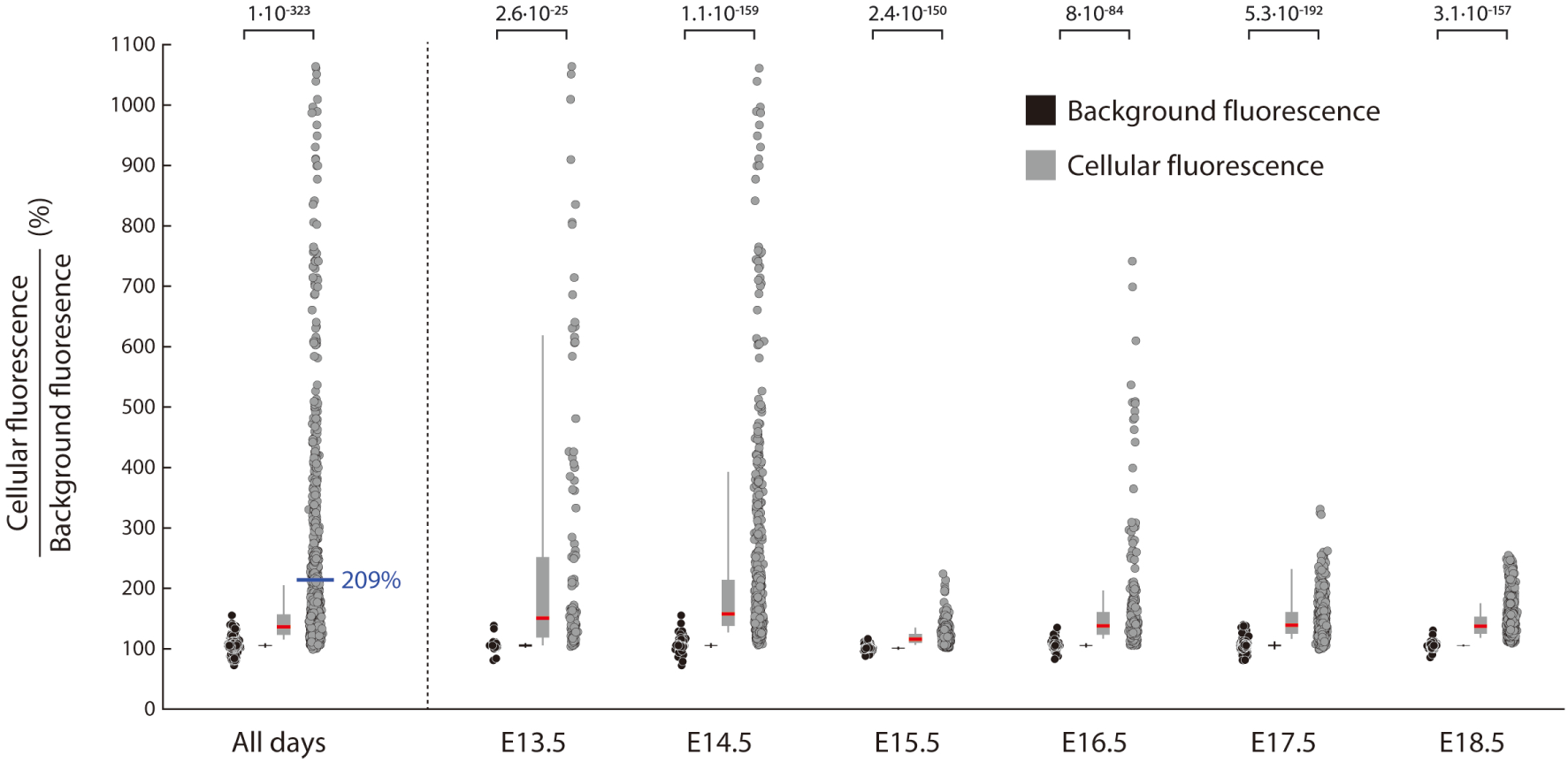
Fluorescence of Rbp4-Cre neurons is significantly increased over background fluorescence from E13.5 to E18.5. Mean cellular fluorescence, normalized by the pixel size of each cell, compared against the mean pixel fluorescence within three background regions selected within the imaging window, collected across all neurons, across all embryonic days (left) and on each embryonic day (right). Black circles: ratio of fluorescence within individual background regions in each imaging plane compared to each other; gray circles: ratio of fluorescence within individual neurons compared to each background region in the same imaging plane; box-and-whiskers: distributions across background fluorescence ratios (black) and cellular fluorescence ratios (gray) as box (25-75 percentile) and whisker (10-90 percentile); black and red line: median; blue line and text: mean. Probability: Wilcoxon rank-sum test comparing cellular fluorescence ratios to background fluorescence ratios. Recordings from 3 (E13.5), 9 (E14.5), 5 (E15.5), 4 (E16.5), 5 (E17.5), and 6 (E18.5) embryos.

**Fig. S7. Associated with Fig. 1.**
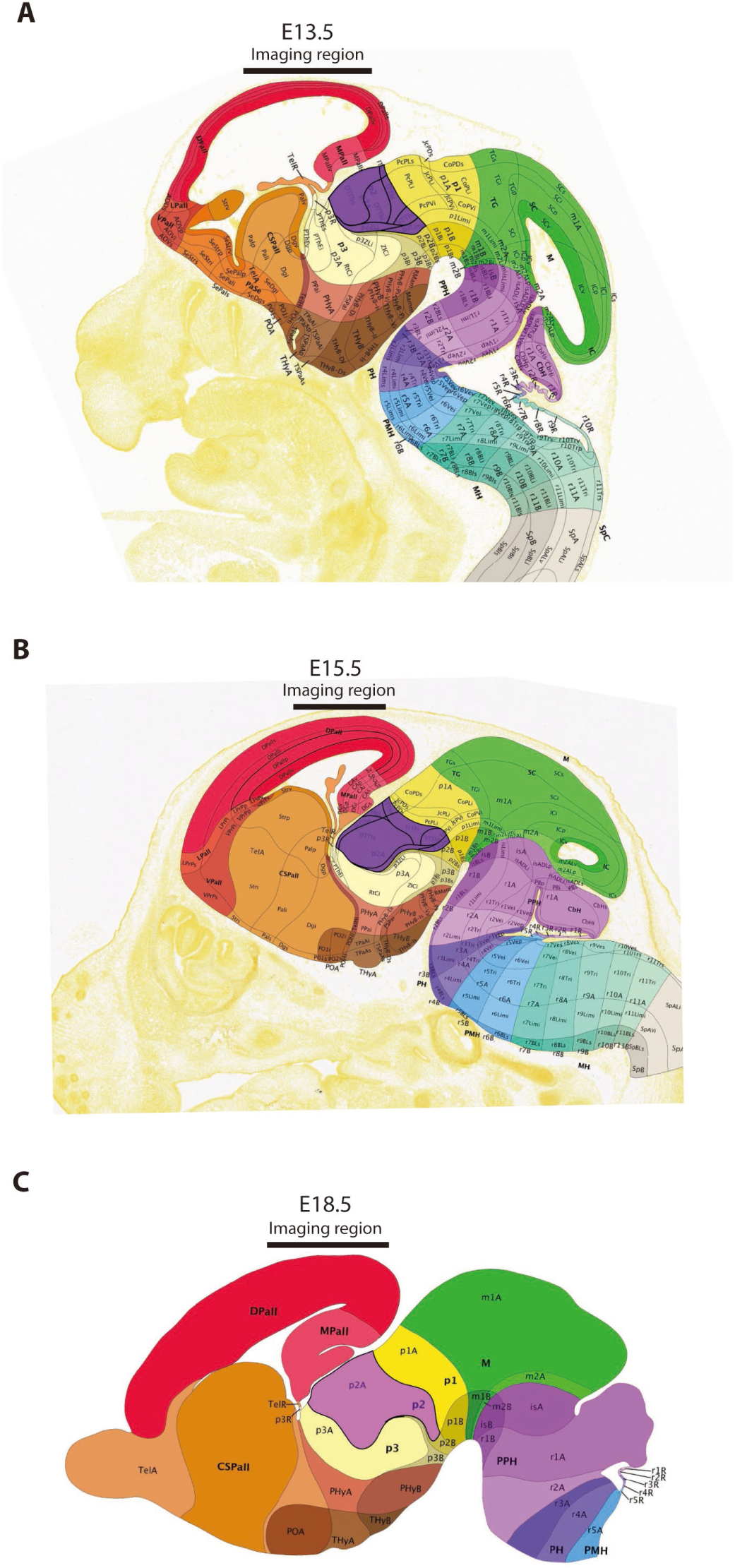
Imaging region with respect to the embryonic brain at E13.5 (A), E15.5 (B), and E18.5 (C). Imaging region was centered over the posterior dorsal pallium. Images taken from Allen Developing Brain Atlas (http://atlas.brain-map.org/) ([CSL STYLE ERROR: reference with no printed form.]).

**Fig. S8. Associated with Fig. 1.**
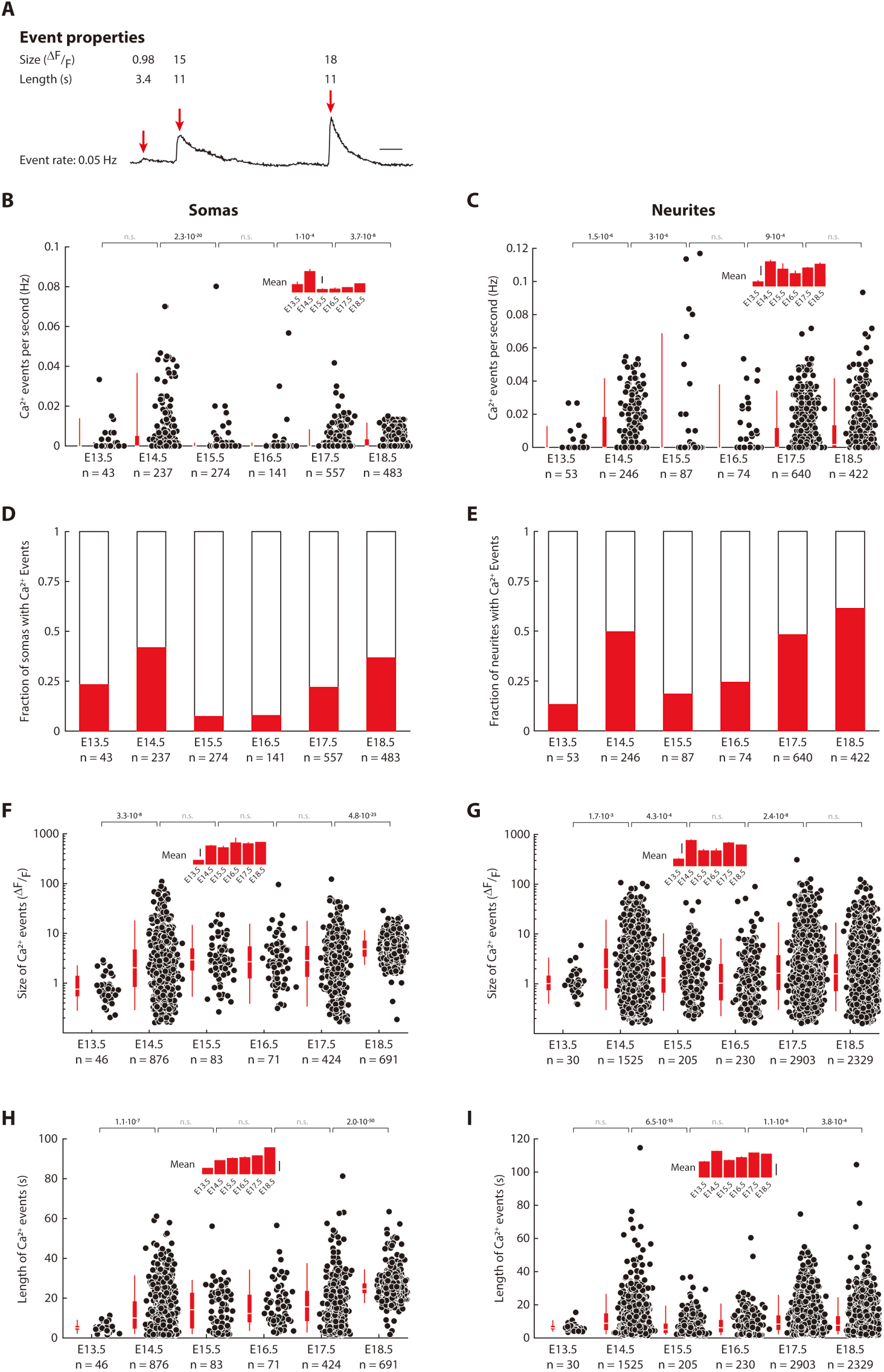
Fraction of Rbp4-Cre ROIs with calcium events, as well as the frequency, length, and amplitude of calcium events all vary across embryonic days. (**A**) Spontaneous calcium activity recorded from a single Rbp4-Cre neuron, showing three detected events, with event properties quantified for each event. (**B – I**) Quantification of event properties for Rbp4-Cre somas (**B, D, F,** and **H**) and neurites (**C, E, G,** and **I**). On each embryonic day from E13.5 to E18.5, these event statistics include the event rate (**B** and **C**), fraction of Rbp4-Cre ROIs showing spontaneous calcium events in each 10-minute recording (**D** and **E**) and, for each detected calcium event, the size (**F** and **G**) and length (**H** and **I**) of the event. (**B – E**) n = number of Rbp4-Cre ROIs (somas or neurites) recorded from 3 (E13.5), 9 (E14.5), 5 (E15.5), 4 (E16.5), 5 (E17.5), and 6 (E18.5) embryos. (**F – I**) n = number of detected calcium events. Scale bar: 5s (A), 2·10^-3^ Hz (inset, B), 4·10^-3^ Hz (inset, C), 2 ΔF/F (inset, F), 2 ΔF/F (inset, G), 10s (inset, H), 10s (inset, I).

**Fig. S9. Associated with Fig. 1.**
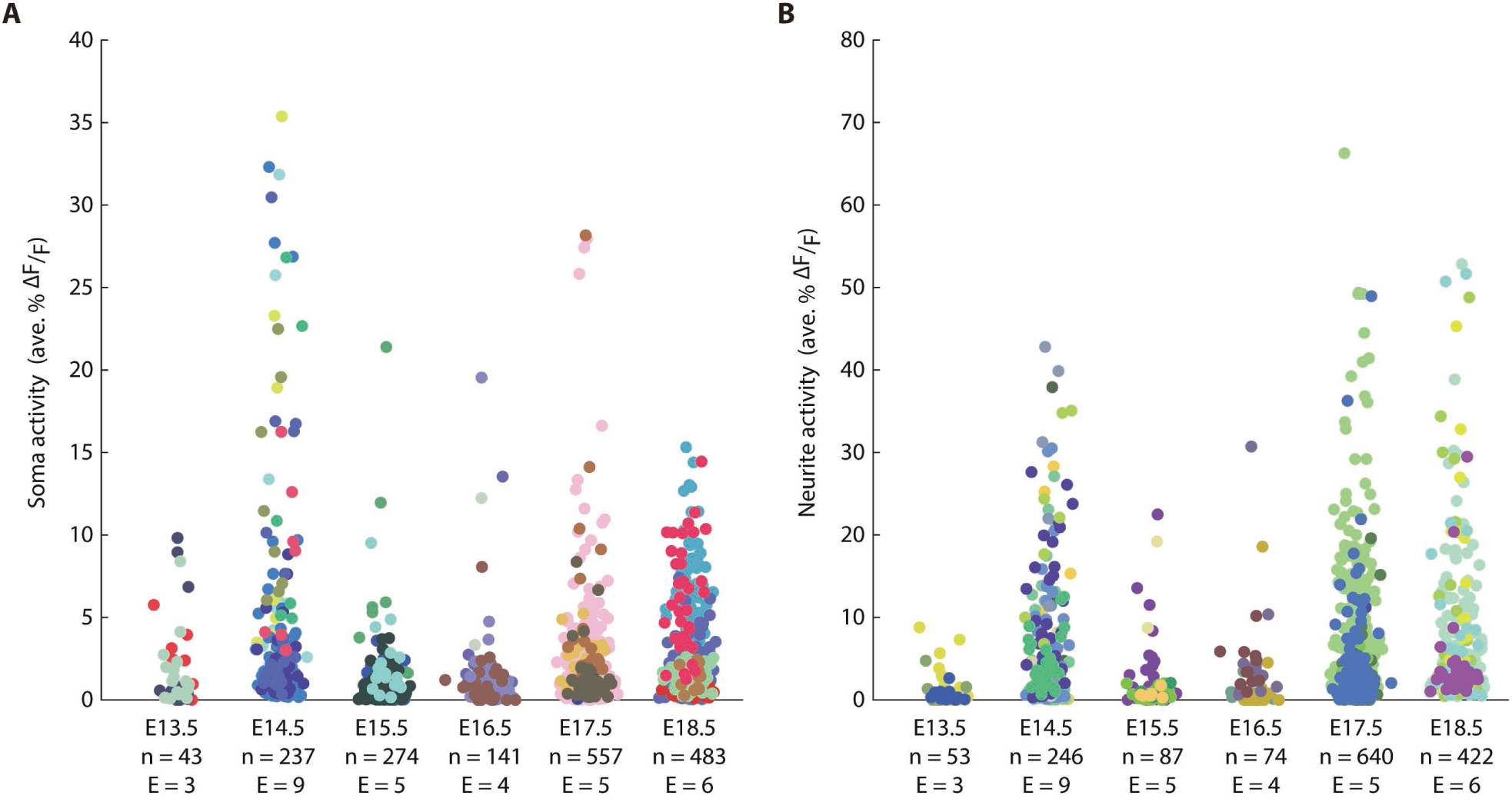
Overall calcium activity in Rbp4-Cre neurons from E13.5 to E18.5, colored by embryo. Activity in the somas (**A**) and neurites (**B**) of Rbp4-Cre neurons (data from Fig. 1J and K) with neurons recorded in the same embryo colored identically.

**Fig. S10. Associated with Fig. 1.**
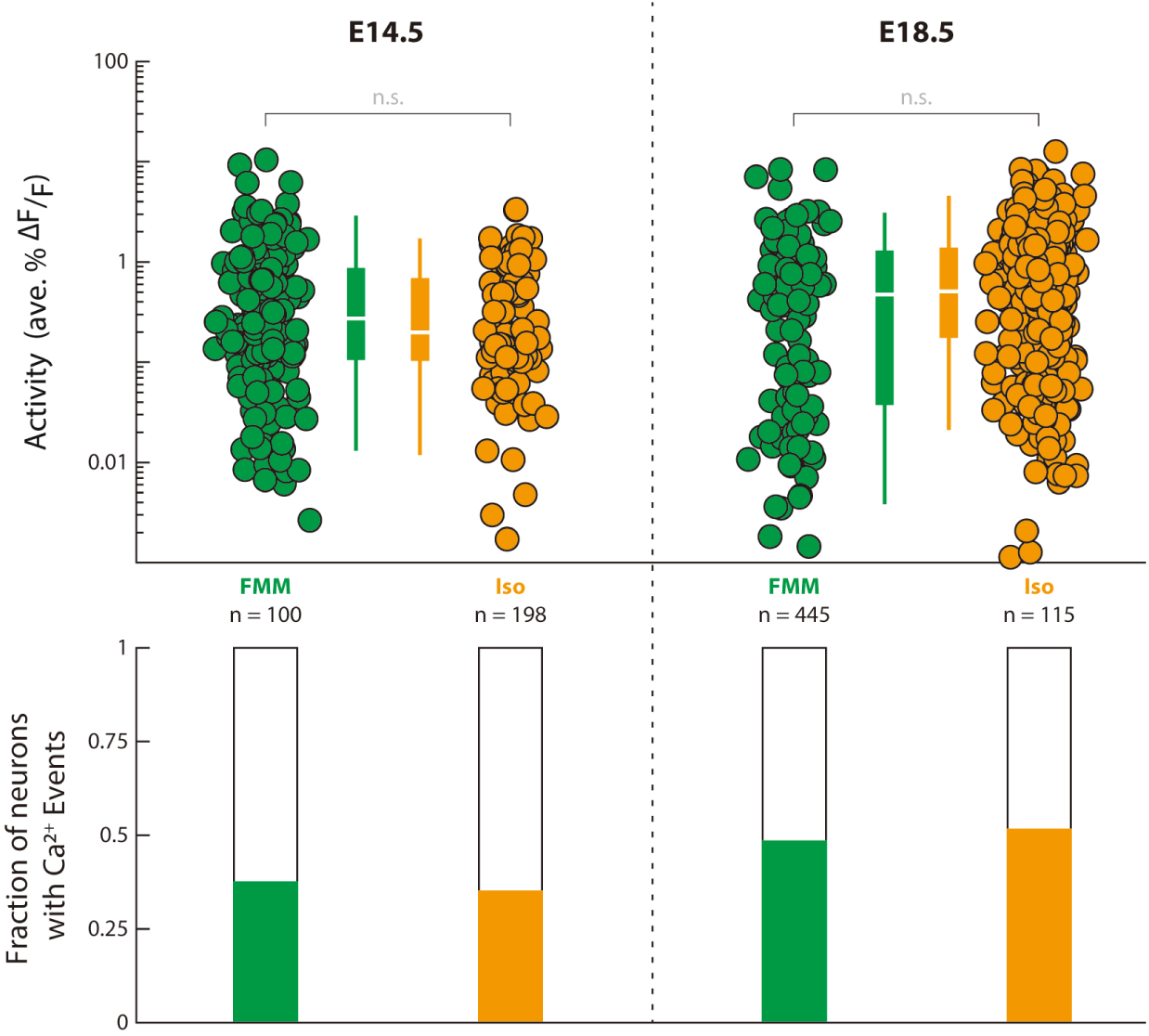
Varying the anesthesia in the dam does not change the overall calcium activity in Rbp4-Cre neurons, during both embryonic phases of increased activity. In vivo para-uterine two-photon imaging of Rbp4-Cre neurons was performed at E14.5 (left) and E18.5 (right), to characterize the activity during both active phases, with the dam anesthetized using two different anesthetics: a mixture of Fentanyl-Medetomidine-Midazolam (FMM), and 1.75% Isoflurane. Probability: Wilcoxon rank-sum test. n = number of Rbp4-Cre neurons recorded from 8 (E14.5) and 6 (E18.5) embryos.

**Fig. S11. Associated with Fig. 2.**
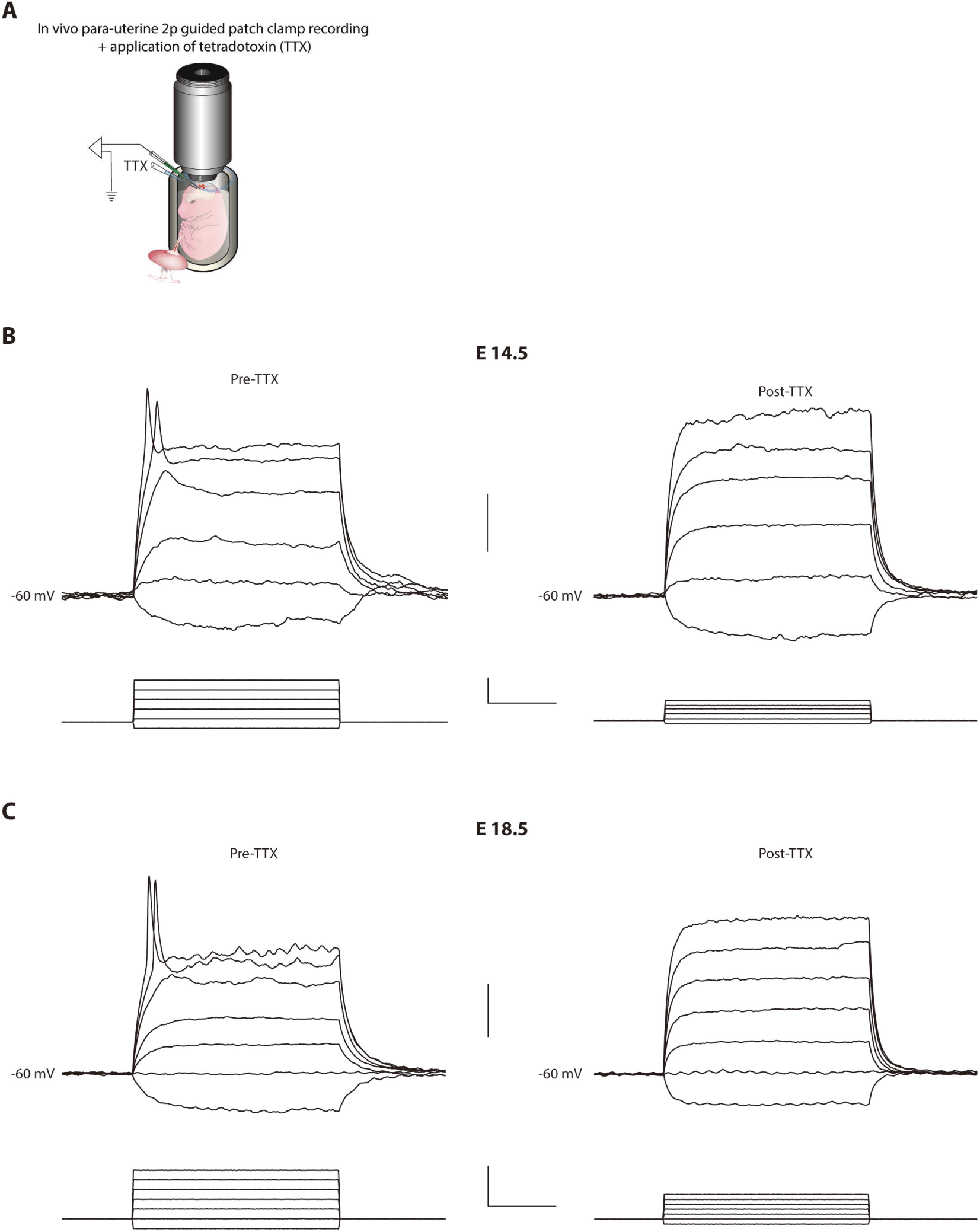
Active conductances in Rbp4-cre neurons are dependent on voltage gated sodium channels. (**A**) Schematic of para-uterine imaging combined with patch clamp recording to perform in vivo two-photon targeted electrophysiology from Rbp4-cre neurons together with application of TTX. (**B** and **C**) Voltage responses (top) of an Rbp4-cre neuron (at E14.5 (**B**) and E18.5 (**C**)) to graded intracellular current injections (bottom), recorded in the current clamp configuration before (left) and after (right) application of TTX. Scale bar: 20 mV (top, B), 10 ms and 40 pA (bottom, B), 20 mV (top, C), 10 ms and 40 pA (bottom, C)

**Fig. S12. Associated with Fig. 2.**
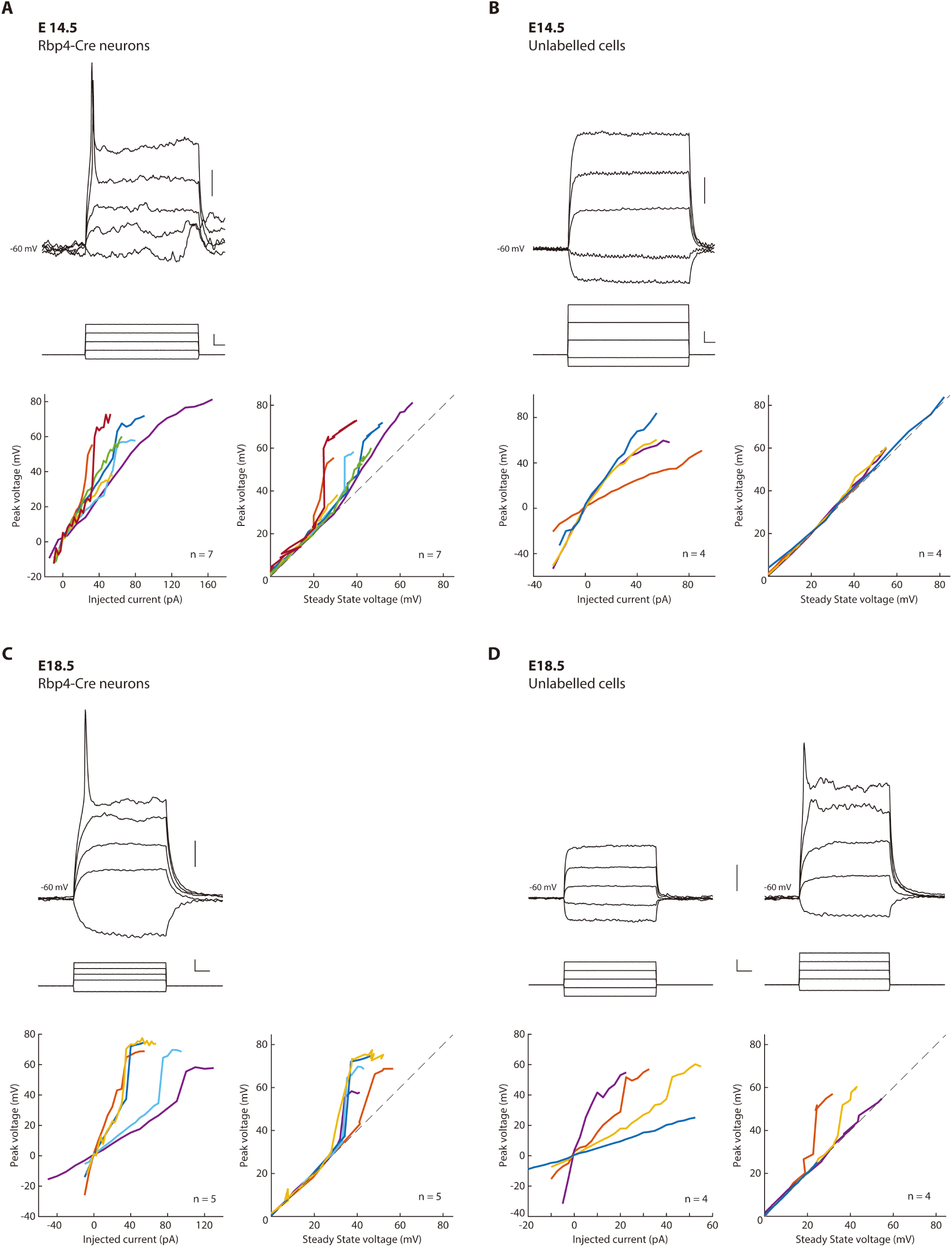
All Rbp4-Cre neurons, but not all nearby unlabelled cells, display active conductances, at both E14.5 and E18.5. (**A – D**) Voltage responses (top) of an Rbp4-Cre neuron (left) and a nearby unlabelled cell (right) at E14.5 (**A** and **B**) and E18.5 (**C** and **D**) to graded intracellular current injections (middle), recorded in the current clamp configuration. The voltage responses to current are quantified in both current-voltage curves and the peak voltage compared to the steady-state voltage (as in Fig. 2C) for both Rbp4-Cre neurons and nearby unlabeled neurons, to visualize any nonlinearity in the peak voltage response. Scale bar: 10 mV (top, A), 50 ms and 20 pA (middle, A), 10 mV (top, B), 50 ms and 20 pA (middle, B), 10 mV (top, C), 50 ms and 20 pA (middle, C), 10 mV (top, D), 50 ms and 20 pA (middle, D).

**Fig. S13. Associated with Fig. 3.**
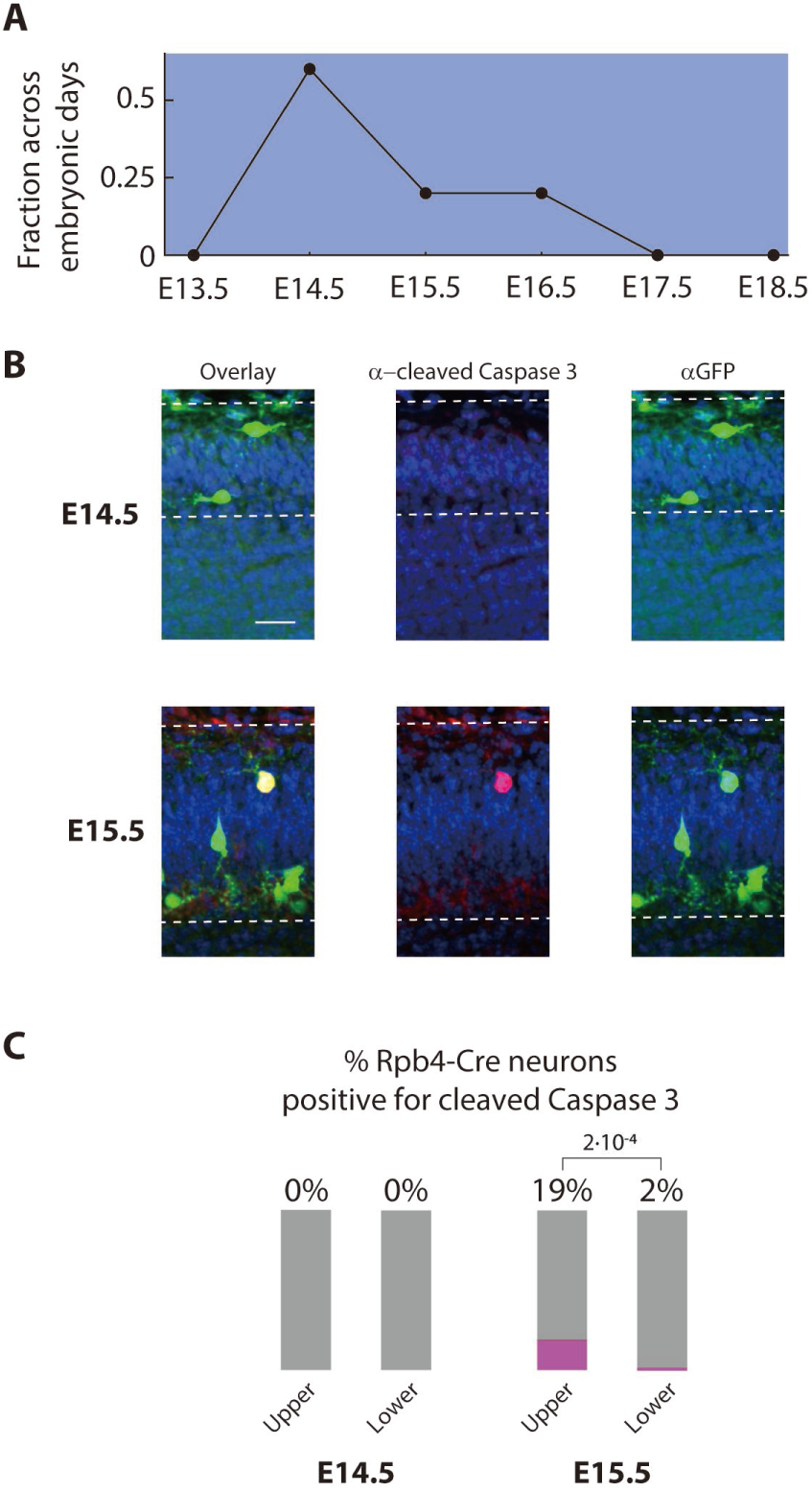
Upper layer Rbp4-Cre neurons decrease in number from E14.5 and express an apoptotic marker at E15.5. (**A**) Fraction of all upper layer Rbp4-Cre neurons found on each embryonic day from E13.5 to E18.5. Layer colored as in Fig. 3A (blue). (**B**) Immunostaining of E14.5 (top) and E15.5 (bottom) embryonic cortex to identify Rbp4-Cre neurons expressing GCaMP6s (GFP antibody, green) and cells expressing cleaved Caspase 3, an apoptotic marker (cleaved Caspase 3 antibody, magenta). (**C**) Fraction of Rbp4-Cre neurons expressing cleaved Caspase 3 at E14.5 and E15.5 in the upper and lower layers. Probability: Fisher exact test comparing the fraction of Rbp4-Cre neurons expressing cleaved Caspase 3 in the upper layer to the lower layer. Scale bar: 20 μm (B).

**Fig. S14. Associated with Fig. 3.**
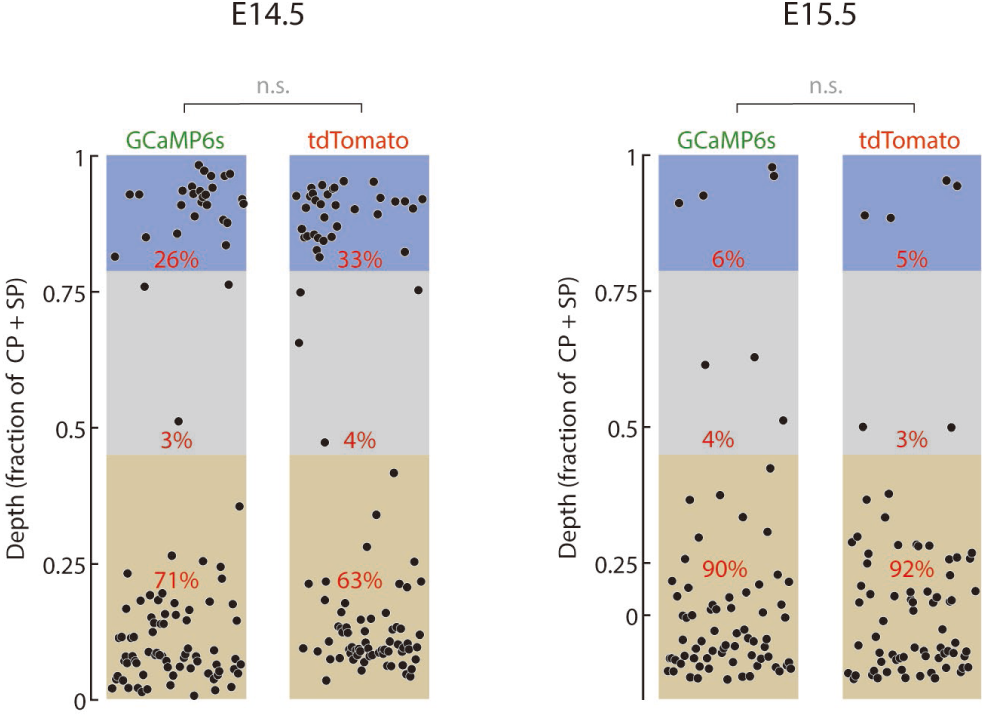
Spatial distribution of Rbp4-Cre neurons into layers is indistinguishable when expressing GCaMP6s compared to tdTomato. Quantifying the distribution of Rbp4-Cre neurons into layers at E14.5 (left) and R15.5 (right) in embryos generated by using the GCaMP6s-tTA2 reporter line, or a tdTomato reporter line. Probability: χ^2^ test comparing the fraction of Rbp4-Cre neurons in each layer for the two different reporter lines.

**Fig. S15. Associated with Fig. 3.**
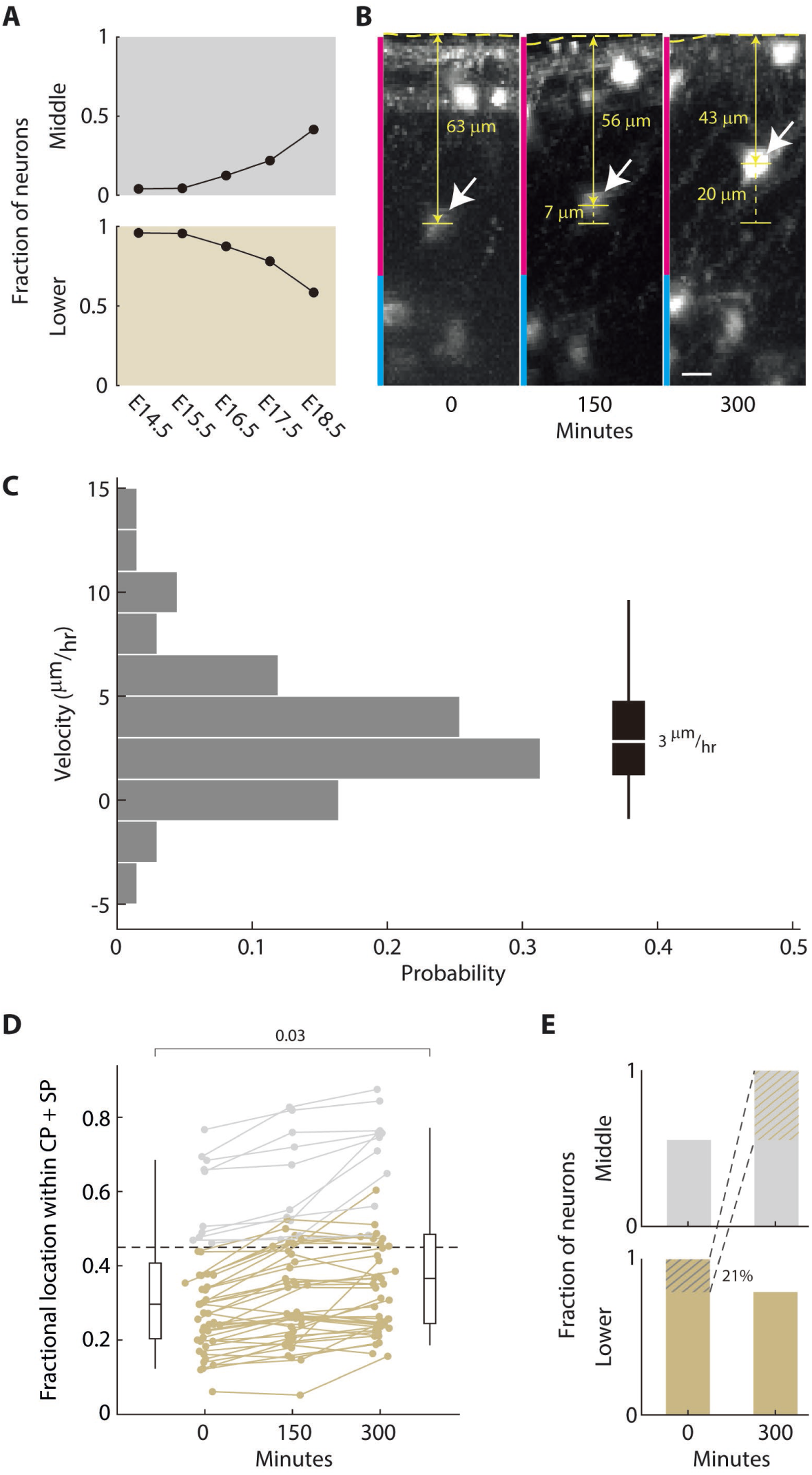
Rbp4-Cre neurons migrate from the lower layer to the middle layer. (**A**) Fraction of all middle and lower layer neurons found within each of the two layers from E14.5 to E18.5. Layers colored as in Fig. 3A (gray: middle layer; beige: lower layer). (**B**) In vivo para-uterine time lapse imaging of a population of Rbp4-Cre neurons labelled with tdTomato, at E16.5, over 5 hours, in the cortical plate (magenta) and subplate (cyan). White arrow: migrating cell; horizontal dotted line: surface of cortex; yellow line: distance of cell to surface; vertical dotted line: distance migrated. (**C**) Distribution of migration velocities for all recorded migrating neurons, averaged across 5 hours of time lapse imaging. n = 48 neurons from 2 E16.5 embryos. (**D**) Locations of layer 5 neurons in middle (gray) and lower (beige) layers (colored based on position at t = 0), followed using para-uterine time lapse imaging for 5 hours. Shown as a fraction of the combined cortical plate (CP) and subplate (SP) thickness. Box-and-whiskers: distribution of neuronal locations at the start of imaging (left) and after 300 minutes (right) as box (25-75 percentile) and whisker (5-95 percentile); black line: median; dotted line: boundary between lower and middle layer at t = 0; p-value: Wilcoxon rank-sum test; n = 48 neurons from 2 E16.5 embryos. (**E**) Fraction of Rbp4-Cre neurons within each layer at the start of imaging (left) and after 300 minutes (right), normalized to the maximum within each layer (data from (**D**)). 21% of lower layer neurons move from the lower layer to the middle layer within the imaging period (colored shading), and no neurons move in the reverse direction. Scale bar: 10 μm (B).

**Fig. S16. Associated with Fig. 3 and 4.**
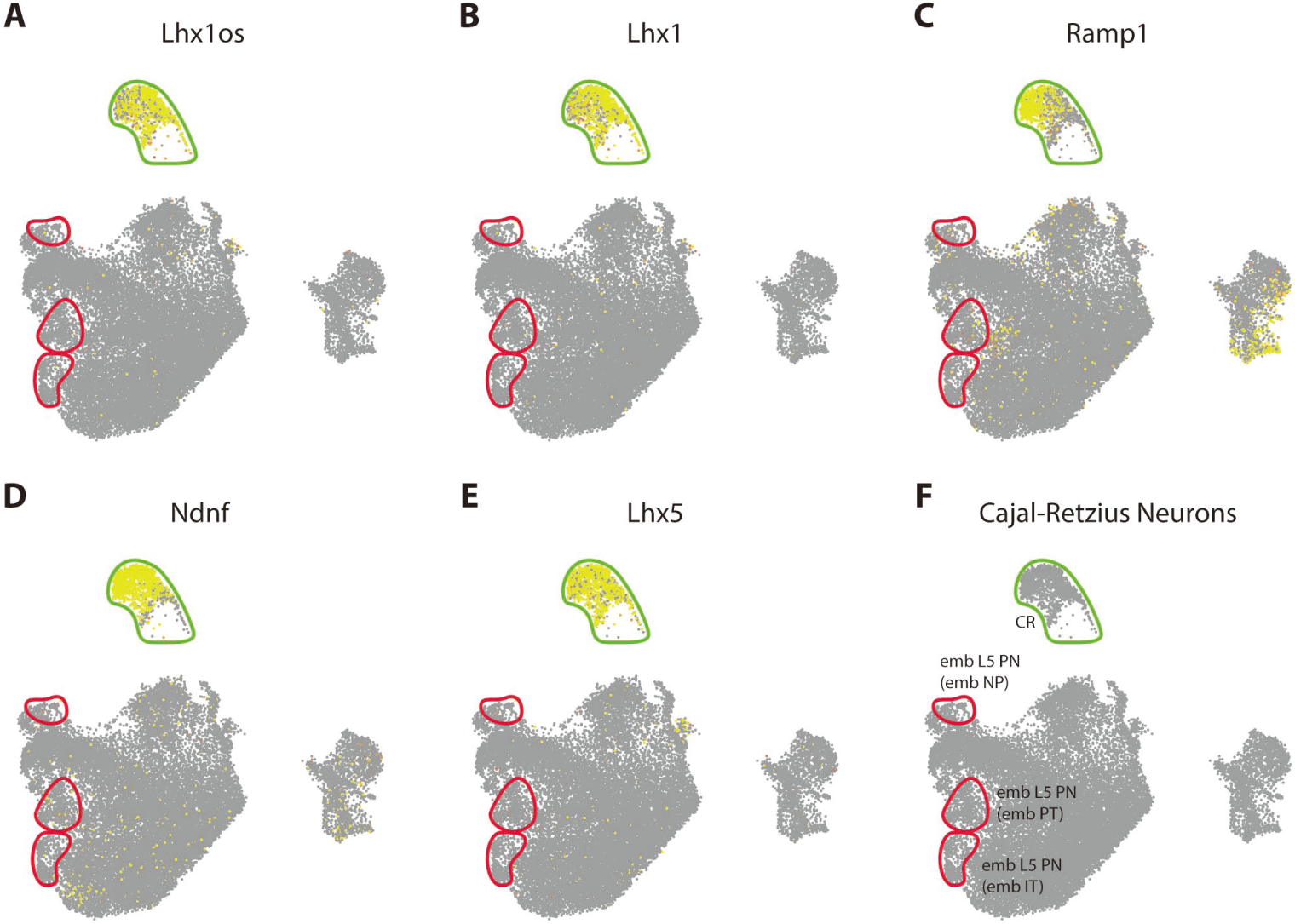
Genes associated with Cajal-Retzius cells identify a population of cells with transcriptomes distinct from Rbp4-Cre neurons. Within the UMAP embedding of all 25681 sequenced excitatory neuron transcriptomes (gray), cells expressing a number of genes previously associated with Cajal-Retzius cells (Bandler et al., 2021; Di Bella et al., 2021; Kuang et al., 2010) (**A – E**) overlap in a location (green outline) distinct from positively identified Rbp4-Cre neurons (as in Fig. S1) (red outline; embryonic L5 PN) (labeled in **F**). Clusters of Rbp4-Cre neurons are additionally labeled based on the cluster identification from Fig. 4B.

**Fig. S17. Associated with Fig. 3 and 4.**
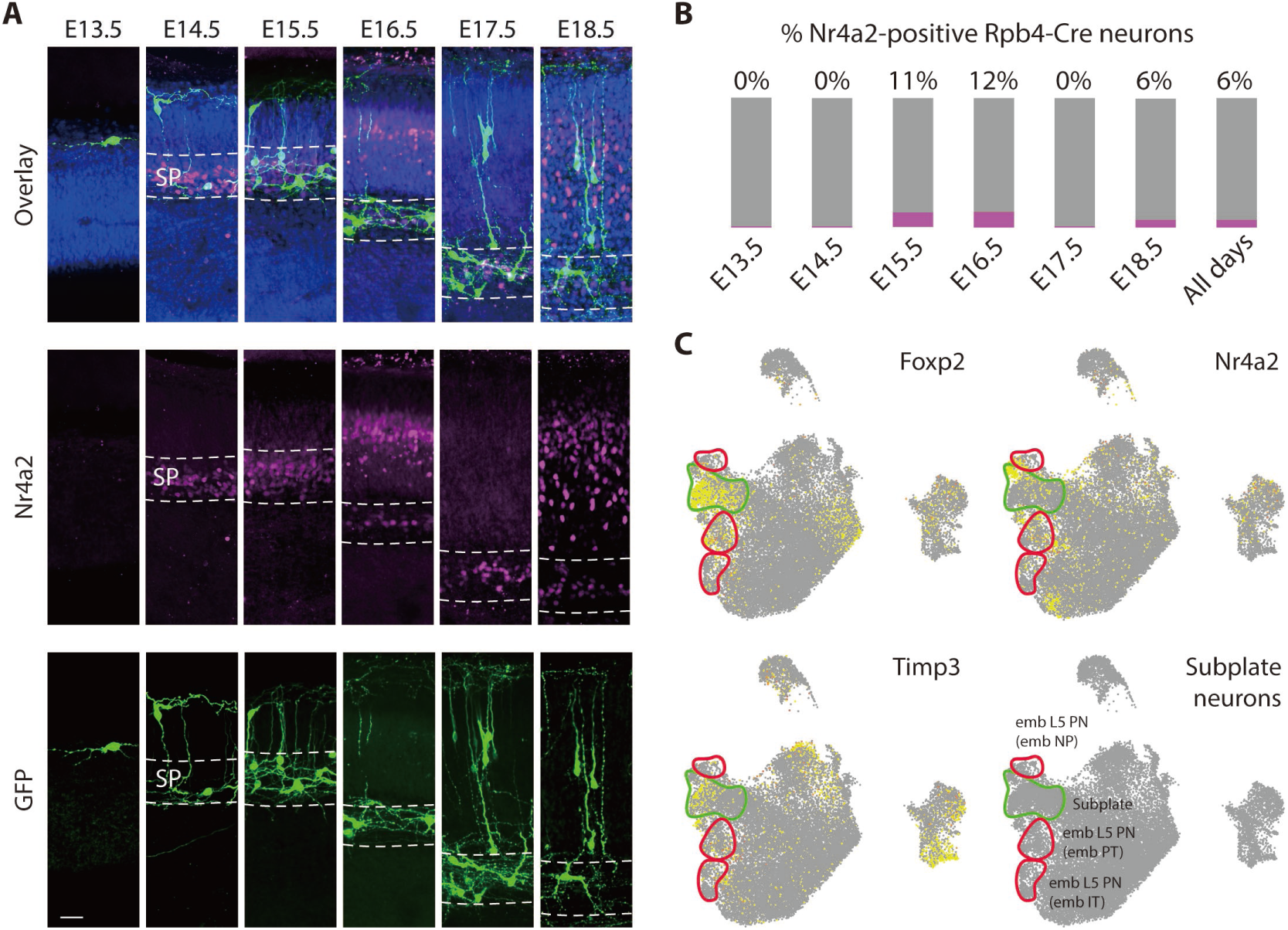
Subplate neurons are both physically close but distinct from lower layer Rbp4-Cre neurons and transcriptomically close to but distinct from all Rbp4-Cre neurons. (**A**) Rbp4-Cre neurons (green), counterstained with Hoechst (blue), show that lower layer neurons lie physically within the subplate (SP; dotted line), as localized by the expression of a common subplate marker, Nr4a2 (magenta) (**B**) Only a small fraction of Rbp4-Cre neurons express Nr4a2 from E13.5 to E18.5. (B) Within the UMAP embedding of all 25681 sequenced excitatory neuron transcriptomes (gray), cells expressing genes previously associated with subplate neurons overlap in a location (green outline) distinct from positively-identified Rbp4-Cre neurons (as in Fig. S1) (red outline; embryonic L5 PNs) (labeled in the bottom right). Clusters of Rbp4-Cre neurons are additionally labeled based on the cluster identification from Fig. 4B. Scale bar: 20 μm (A).

**Fig. S18. Associated with Fig. 3.**
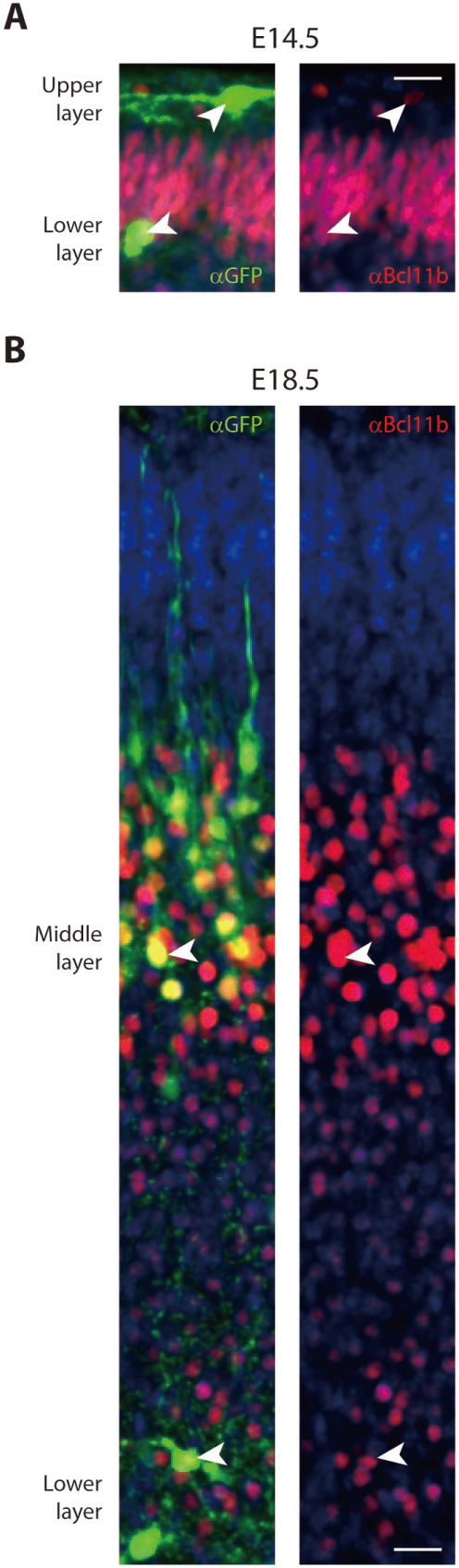
Rbp4-Cre neurons through the depth of the embryonic cortex, in the upper, middle, and lower layers, express Bcl11b. Rbp4-Cre neurons (stained using GFP antibody, green), counterstained with Hoechst (blue), show that upper and lower layer neurons at E14.5 (A), as well as middle layer and lower layer neurons at E18.5, all colocalize with Bcl11b (red), the expression of which is restricted in the adult cortex to layer 5 (Arlotta et al., 2005). Arrows: example Rbp4-Cre neurons. Scale bars: 20 μm (A), 20 μm (B).

**Fig. S19. Associated with Fig. 4.**
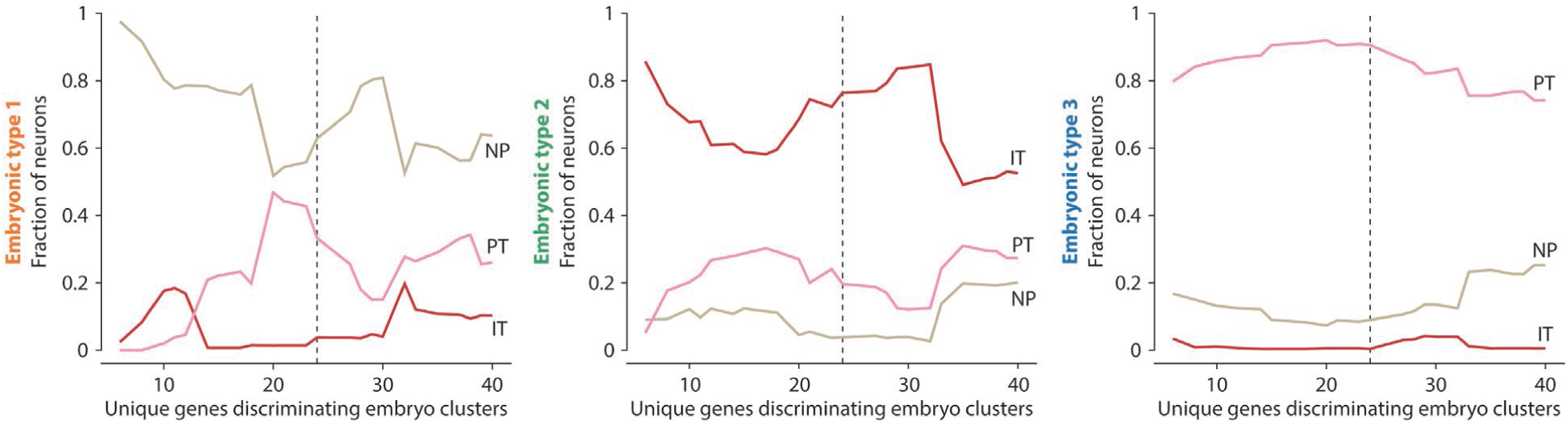
Each embryonic type associates with one adult layer 5 type, stably across an increasing number of genes. Fraction of neurons within each embryonic type (1 (left), 2 (middle), and 3 (right)) with a significant conditional probability of being sampled from the expression profile of one of the three adult cell types. Adult types: near-projecting neurons (NP) (beige), intratelencephalic neurons (IT) (red), and pyramidal tract neurons (PT) (pink) (as sequenced and identified in (Tasic et al., 2018) from VISp). Genes were selected to best discriminate the embryonic clusters, while still being differentially expressed across the three adult types. Dotted line: 24 differentially expressed genes used to generate Fig. 4B.

**Fig. S20. Associated with Fig. 4.**
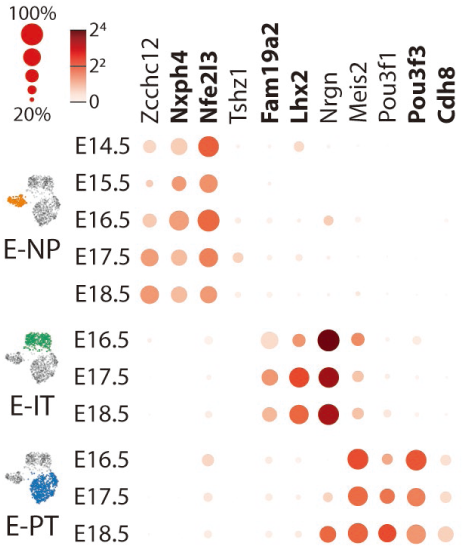
Genes selective for each embryonic layer 5 type. Transcript counts for each gene shown as colored circles in all types at all ages. Radius of circle: fraction of cells expressing the gene; color of circle: mean normalized transcripts per cell (log_2_). Bold text: genes used for in situ hybridization, shown in Fig. 4D.

**Fig. S21. Associated with Fig. 5.**
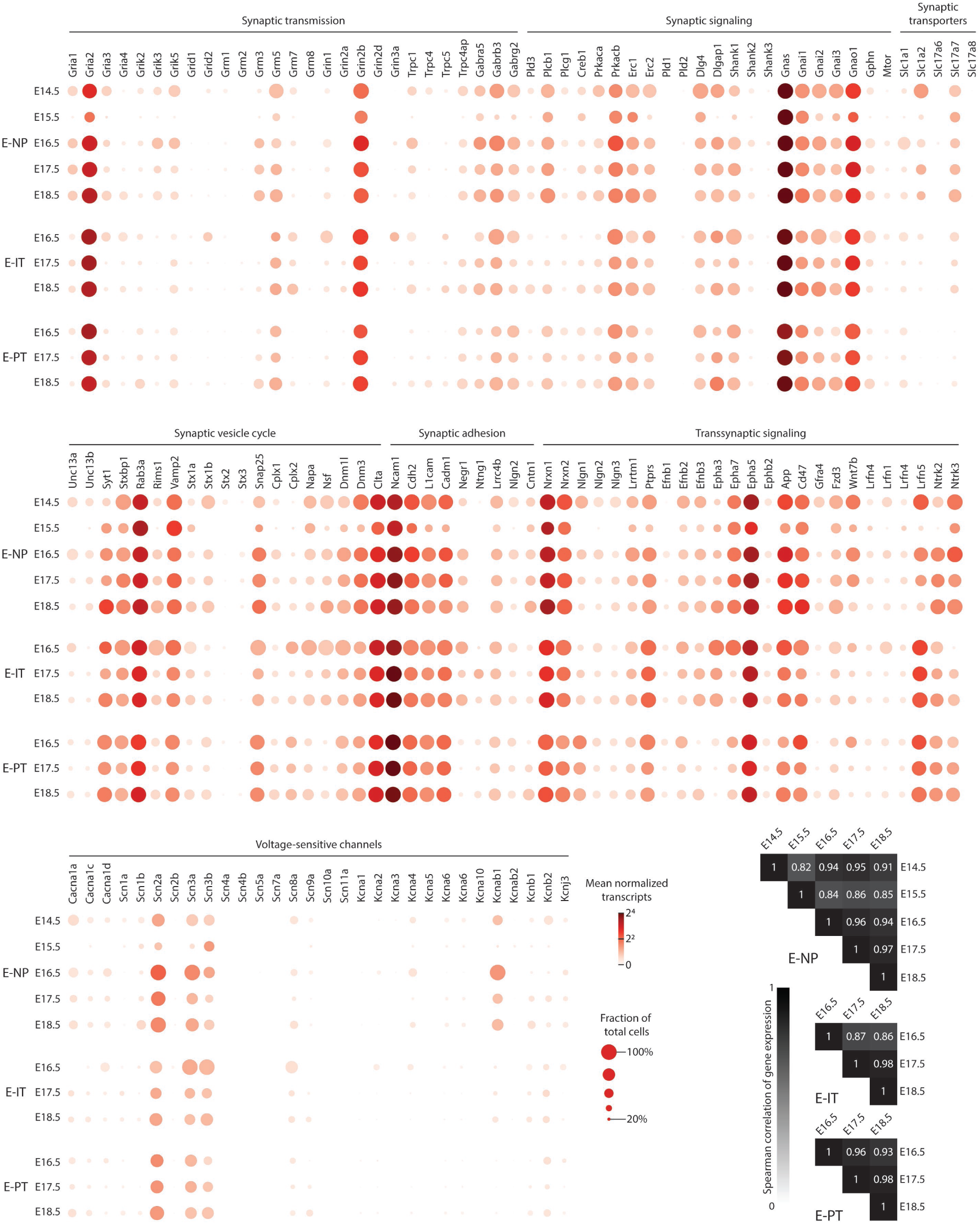
All three embryonic layer 5 pyramidal cell types express genes related to synaptic formation, synaptic function, and neuronal active membrane properties, from E14.5 to E18.5. Genes selected from (Kanehisa and Goto, 2000; Siddiqui and Craig, 2011). Transcript counts shown as colored circles. Radius of circle: fraction of cells expressing the gene; color of circle: mean normalized transcripts per cell (log_2_). Bottom right: correlation of transcript counts of all selected genes within each embryonic layer 5 pyramidal type compared across pairs of embryonic days are > 0.8 across all types and days. Colored from 0 (white) to 1 (black).

**Fig. S22. Associated with Fig. 6.**
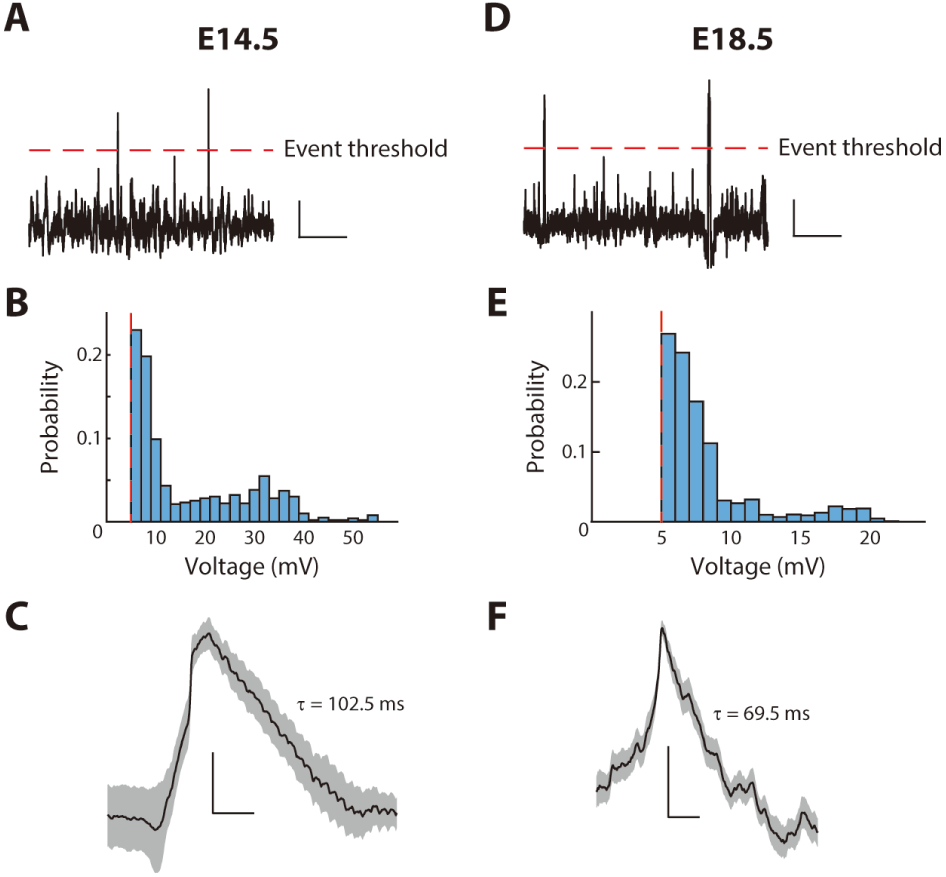
Rbp4-Cre neurons display spontaneous excitatory synaptic potentials, in the presence of TTX, in both active phases of embryonic development, at E14.5 (A – C) and E18.5 (D – F). (**A, D**) Traces recorded in current clamp configuration during in vivo two-photon targeted patch clamp recordings from Rbp4-Cre neurons, following the application of TTX to the surface of cortex. Synaptic potentials were defined as deflections greater than 5 mV (dotted line: event threshold). (**B, E**) Distribution of amplitudes greater than the event threshold. (**C, F**) Average across events, normalized to the peak voltage of each event, showing the characteristic shape of a synaptic potential. Decay time (τ) shown in ms. Scale bars: 5s and 2.5 mV (A), 50 ms and 25% of peak (C), 5s and 2.5 mV (B), 50 ms and 25% of peak (F).

**Fig. S23. Associated with Fig. 6.**
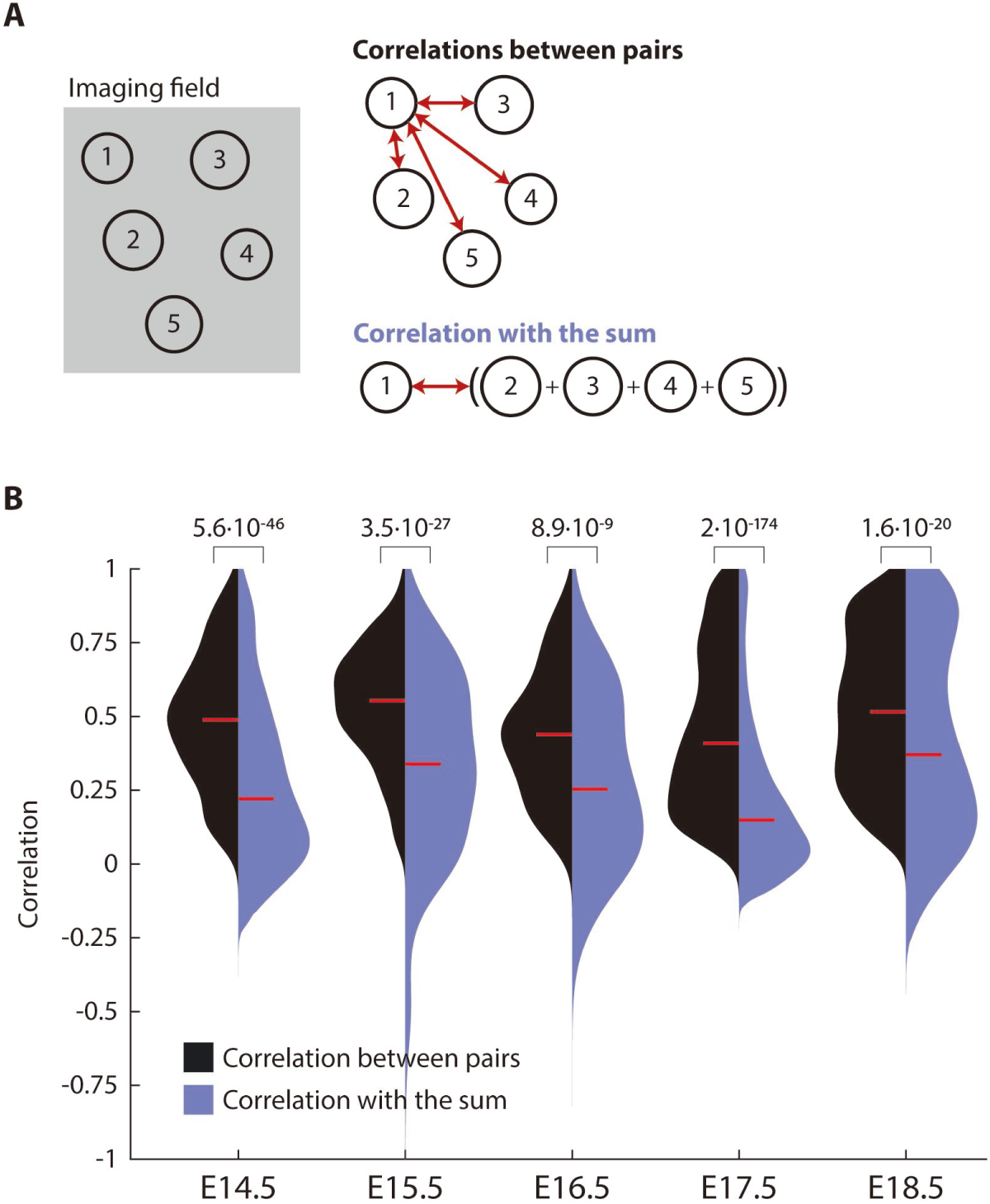
Population activity across Rbp4-Cre neurons is less correlated than activity across pairs of Rbp4-Cre neurons. (**A**) Schematic comparing the “correlation between pairs” of neurons (black), and “correlation with the sum” of neuronal activity (blue) within all neurons recorded in an imaging field (left). Pairwise correlations of neuron 1 with each other neuron among all imaging field (top right). Correlation of the activity of neuron 1 with the sum of activity in all other neurons in the imaging window (bottom right). (**B**) From E14.5 to E18.5, the distribution of correlations between pairs of neurons (that are significantly greater than random) (black shaded area) is compared to the distribution of correlations with the sum (for all recorded neurons) (blue shaded area). Probability: Wilcoxon rank-sum test. Correlations computed from recordings in 3 (E13.5), 9 (E14.5), 5 (E15.5), 4 (E16.5), 5 (E17.5), and 6 (E18.5) embryos.

**Fig. S24. Associated with Fig. 6.**
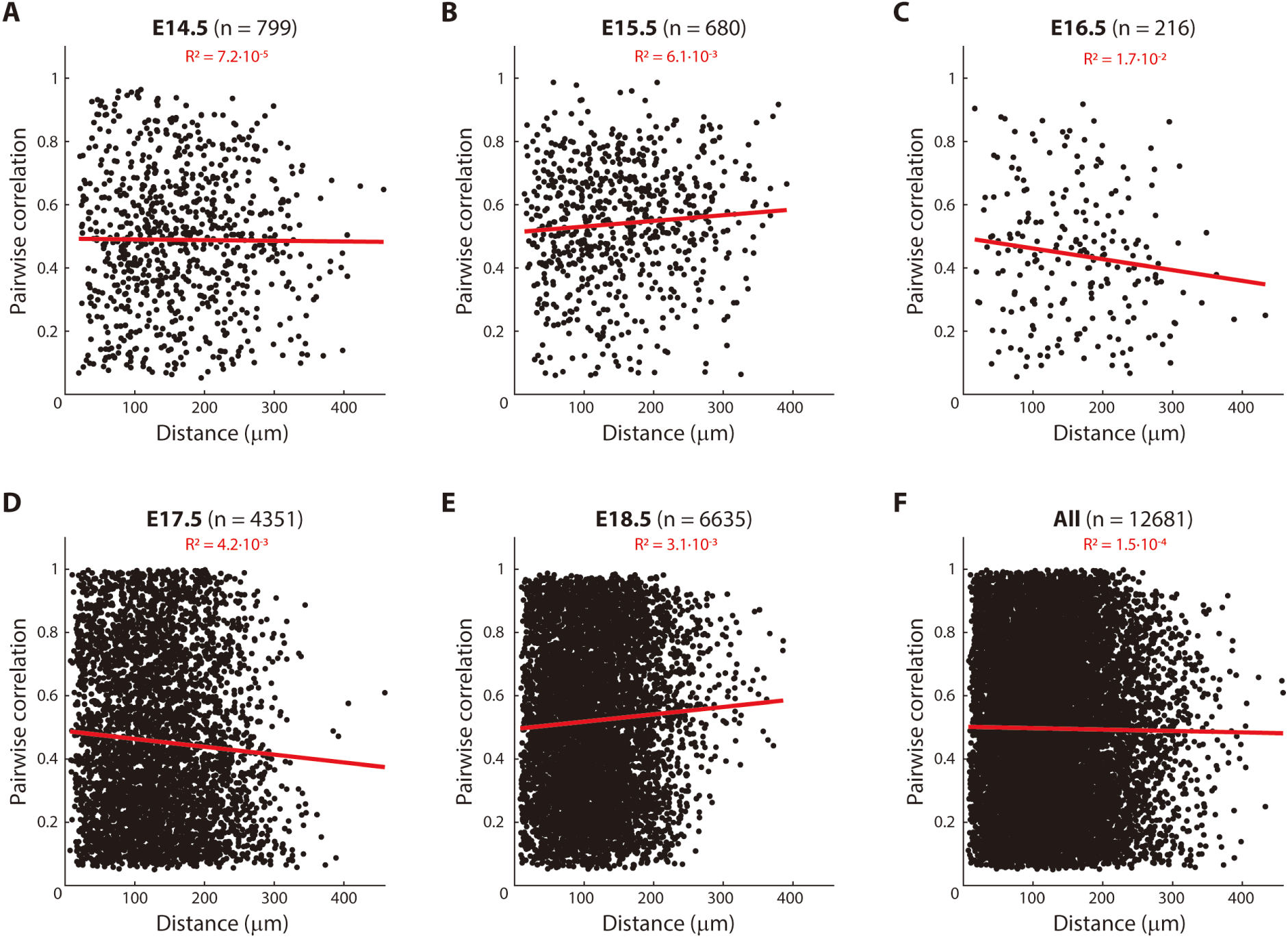
Correlated activity between pairs of Rbp4-Cre neurons does not decrease with distance. Correlations of spontaneous calcium activity, that are significantly greater than random, between pairs of Rbp4-Cre neurons on each embryonic day from E14.5 to E18.5 (**A – E**), displayed with respect to the distance between each pair of neurons. (**F**) Combining all pairwise correlations, significantly greater than random, across all embryonic days. (**A – F**) Red line: best fit trend of correlations across distance (red: R^2^ quality of fit); n = number of pairs from recordings in 3 (E13.5), 9 (E14.5), 5 (E15.5), 4 (E16.5), 5 (E17.5), and 6 (E18.5) embryos.

**Fig. S25. Associated with Fig. 7.**
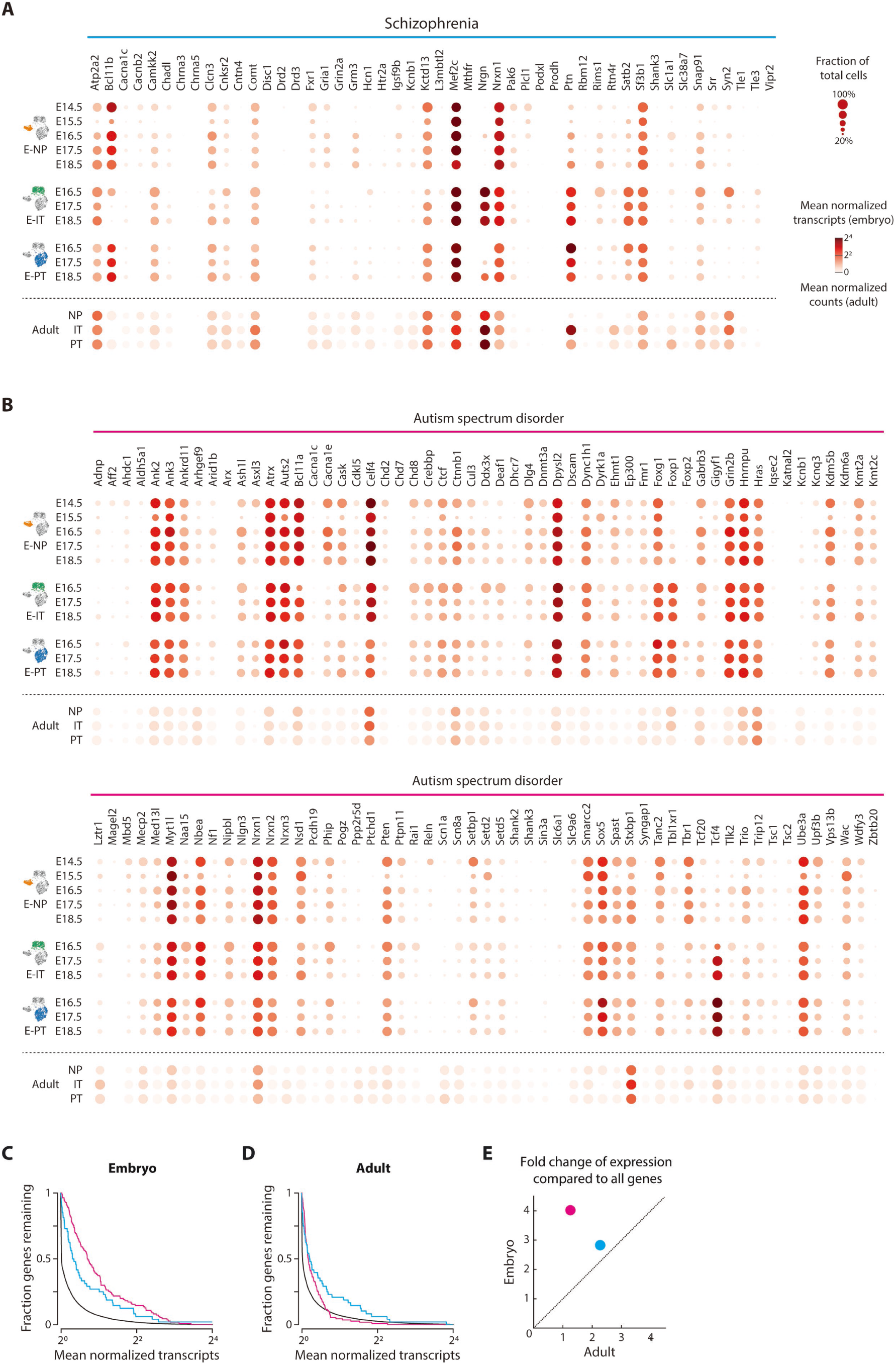
Expression of genes associated with schizophrenia and autism spectrum disorder in all three embryonic layer 5 pyramidal neuron types across embryonic days and across adult layer 5 pyramidal types. (**A**) Expression of genes associated with schizophrenia is shown for every embryonic layer 5 pyramidal neuron type across embryonic days (top) and for each adult layer 5 pyramidal cell type (bottom) (as sequenced and identified in (Tasic et al., 2018) from VISp). Genes were identified from genome-wide association studies (Lee et al., 2019; Ripke et al., 2014) and supplemented with additional genes reported in the OMIM database ([CSL STYLE ERROR: reference with no printed form.]). (**B**) Expression of 110 genes associated with autism are shown for every embryonic layer 5 pyramidal neuron type across embryonic days and for each adult layer 5 pyramidal cell type (as sequenced and identified in (Tasic et al., 2018) from VISp). Genes selected by SFARI human gene score and greater than 10 reports for each gene (Abrahams et al., 2013). (**A** and **B**) Radius of circle: fraction of cells with non-zero transcripts for each gene; color of circle: log_2_ mean normalized transcripts per cell (embryos) and log_2_ mean normalized counts per cell (adult). (**C** and **D**) Fraction of genes with a mean transcript count greater than any specific mean transcript count are shown for genes associated with schizophrenia (cyan), autism spectrum disorder (magenta), and for all transcribed genes (black) in embryonic layer 5 pyramidal neurons (**C**) and adult layer 5 pyramidal neurons (**D**). (**E**) Fold change of expression of genes associated with schizophrenia (cyan) and autism spectrum disorder (magenta) compared to all genes (for embryonic and adult layer 5 pyramidal neurons).

**Fig. S26. Associated with Fig. 7.**
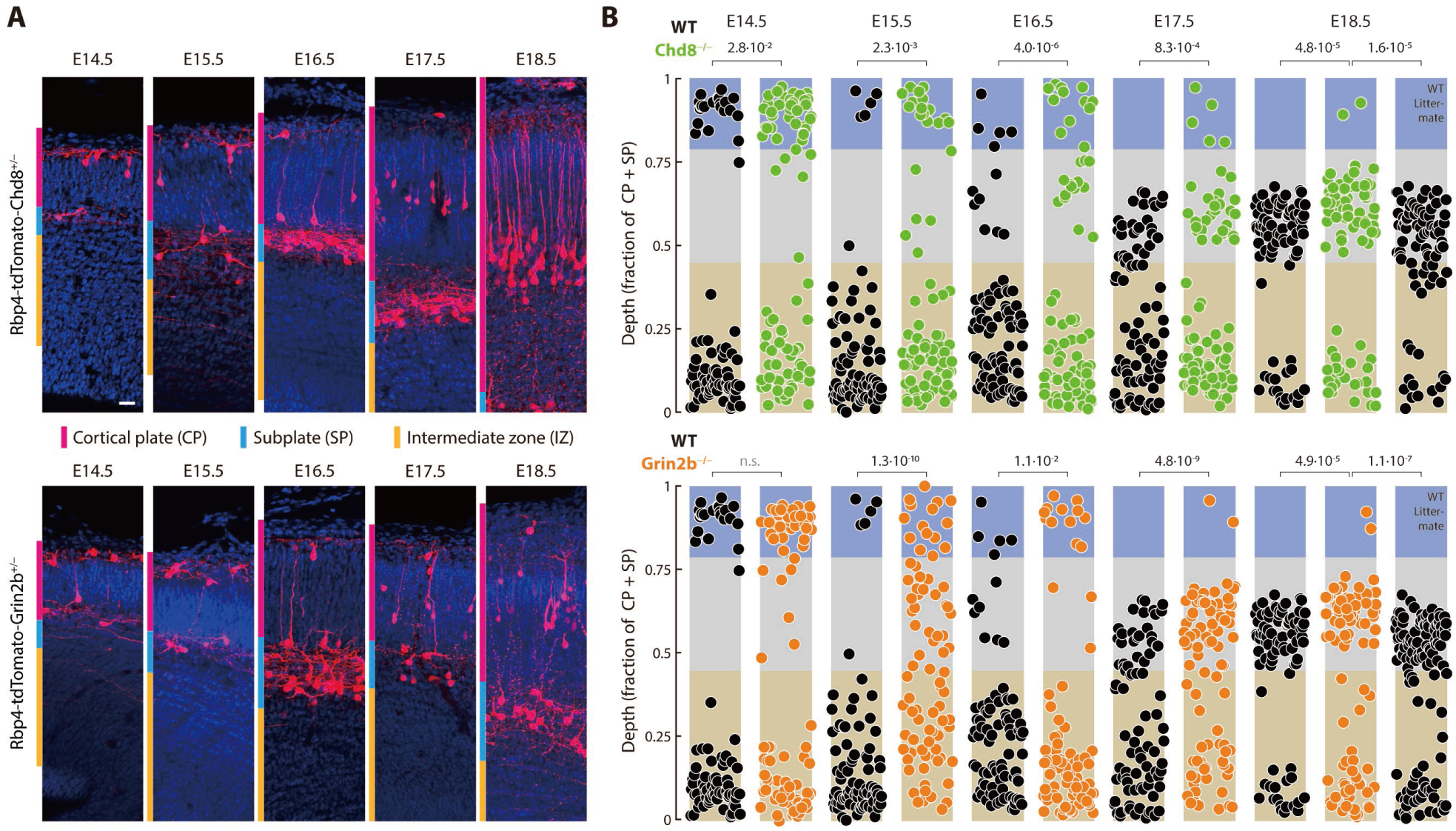
Perturbing autism-associated genes selectively in Rbp4-Cre neurons disrupts organization of layer 5 during embryonic development. (**A**) Rbp4-Cre neurons (stained using tdTomato antibody, red) in Rbp4-tdTomato-Chd8^-/-^ (top) and Rbp4-tdTomato-Grin2b^-/-^ (blue) mice, from E14.5 to E18.5, within the cortical plate (magenta), subplate (cyan), and intermediate zone (yellow), counterstained with Hoechst (blue). Scale bar: 20 µm. (**B**) Distribution of Rbp4-Cre neuronal locations as a fraction of the cortical plate and subplate thickness, from E14.5 to E18.5, in control (WT, black), Rbp4-tdTomato-Chd8^-/-^ (top, purple), and Rbp4-tdTomato-Grin2b^-/-^ (bottom, blue) mice. 85 neurons from each mouse line, sampled at random, displayed on each embryonic day. Layer boundaries derived from Fig. 3A (blue: upper layer; gray: middle layer; beige: lower layer). Probability: χ^2^ test comparing the fraction of Rbp4-Cre neurons in each layer between the conditional knockout mouse and control mice, on each embryonic day; p = 0.05.

**Fig. S27. Associated with Fig. 7.**
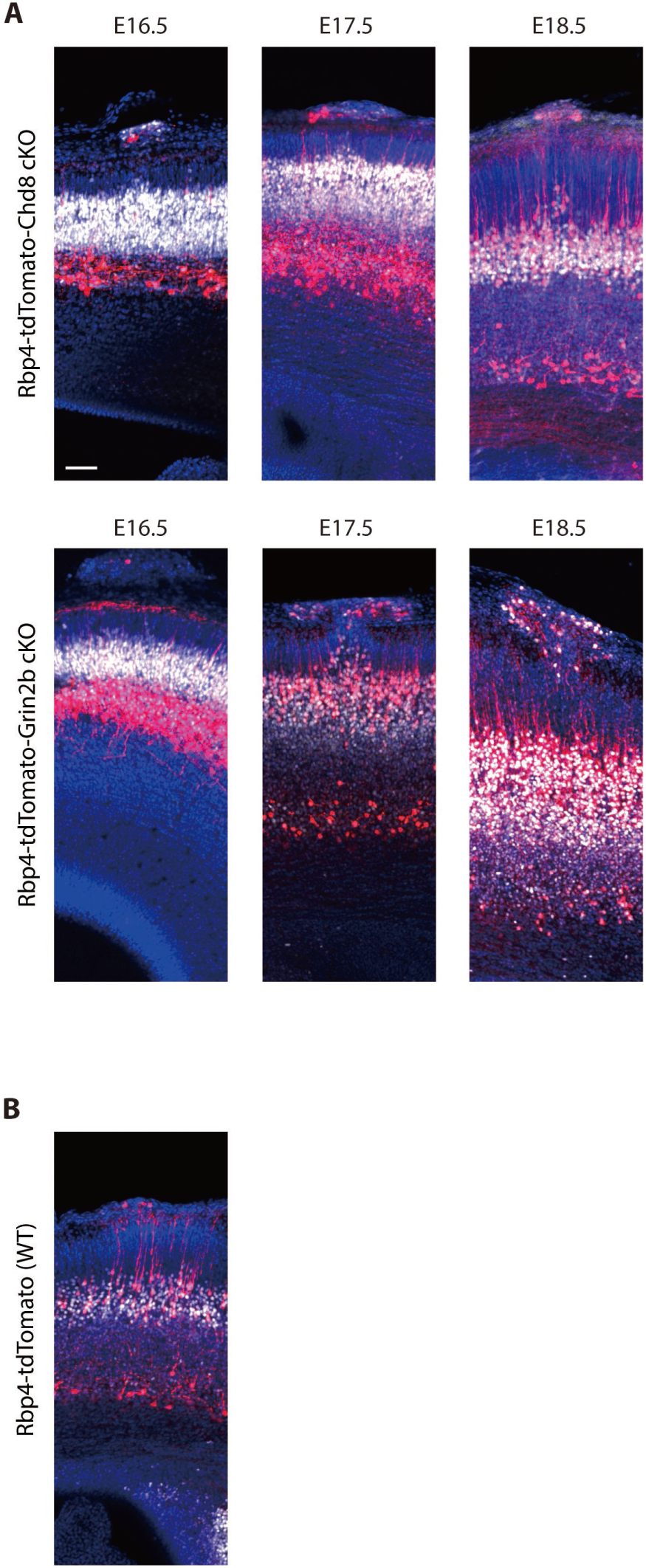
Patches of disorganization within mouse cortex. (**A**) Local patches of disorganization in Rbp4-tdTomato-Chd8 and Rbp4-tdTomato-Grin2b conditional knockout (cKO) mice (examples from each embryonic day from E16.5 to E18.5) show Rbp4-Cre neurons (stained using tdTomato antibody, red) at the surface and disrupted middle layer, including both neurons expressing Cre (red) and not expressing Cre (stained using Bcl11b antibody, white), counterstained with Hoechst (blue). (**B**) Upper layer Rbp4-Cre neurons in a control mouse, without disorganization of the underlying middle layer. Scale bar: 20 µm (**A, B**).

**Video S1. Associated with Fig. 1.**

**Blood flow at the surface of embryonic cortex at E14.5 (2X real-time).** Video was taken after 5 hours of para-uterine imaging and 10 min after the disruption of the umbilical cord.

**Video S2. Associated with Fig. 1.**

**Blood flow at the surface of embryonic cortex at E18.5 (2X real-time).** Video was taken after 5 hours of para-uterine imaging and 10 min after the disruption of the umbilical cord.

**Video S3. Associated with Fig. 1.**

**In vivo para-uterine two-photon imaging of GCaMP6s at E14.5 to show activity in Rbp4-Cre neurons’ somas (10X real-time)**. Imaging GCaMP6s in somas of Rbp4-Cre neurons at E14.5 shows calcium activity. Motion correction was applied to the video, and the fire colormap in FIJI was used to visualize changes.

**Video S4. Associated with Fig. 1.**

**In vivo para-uterine two-photon imaging of GCaMP6s at E14.5 to show activity in Rbp4-Cre neurons’ neurites (15X real-time)**. Imaging GCaMP6s in somas of Rbp4-Cre neurons at E14.5 shows calcium activity. Motion correction was applied to the video, and the fire colormap in FIJI was used to visualize changes. Arrowheads: neurites just prior to calcium events.

**Video S5. Associated with Fig. 1.**

**In vivo para-uterine two-photon imaging of GCaMP6s at E18.5 to show activity in Rbp4-Cre neurons’ somas (30X real-time)**. Imaging GCaMP6s in somas of Rbp4-Cre neurons at E18.5 shows calcium activity. Motion correction was applied to the video, and the fire colormap in FIJI was used to visualize changes.

**Video S6. Associated with Fig. 1.**

**In vivo para-uterine two-photon imaging of GCaMP6s at E18.5 to show activity in Rbp4-Cre neurons’ neurites (15X real-time)**. Imaging GCaMP6s in somas of Rbp4-Cre neurons at E18.5 shows calcium activity. Motion correction was applied to the video, and the fire colormap in FIJI was used to visualize changes. Arrowheads: neurite arbor just prior to calcium events.

**Video S7. Associated with Fig. 1.**

**In vivo para-uterine two-photon imaging of GCaMP6s at E15.5 to show the lack of activity in Rbp4-Cre neurons (15X real-time)**. Imaging GCaMP6s in somas of Rbp4-Cre neurons at E15.5 shows calcium activity. Motion correction was applied to the video, and the fire colormap in FIJI was used to visualize changes.

